# Tabula Sapiens reveals transcription factor expression, senescence effects, and sex-specific features in cell types from 28 human organs and tissues

**DOI:** 10.1101/2024.12.03.626516

**Authors:** The Tabula Sapiens Consortium, Stephen R Quake

## Abstract

The Tabula Sapiens is a reference human cell atlas containing single cell transcriptomic data from more than two dozen organs and tissues. Here we report Tabula Sapiens 2.0 which includes data from nine new donors, doubles the number of cells in Tabula Sapiens, and adds four new tissues. This new data includes four donors with multiple organs contributed, thus providing a unique data set in which genetic background, age, and epigenetic effects are controlled for. We analyzed the combined Tabula Sapiens data for expression of transcription factors, thereby providing putative cell type specificity for nearly every human transcription factor and as well as new insights into their regulatory roles. We analyzed the molecular phenotypes of senescent cells across the entire data set, providing new insight into both the universal attributes of senescence as well as those aspects of human senescence that are specific to particular organs and cell-types. Similarly, we analyzed sex-specific gene expression across all of the identified cell types and discovered which cell types and genes have the most distinct sex based gene expression profiles. Finally, to enable accessible analysis of the voluminous medical records of Tabula Sapiens donors, we created a web application powered by a large language model that allows users to ask general questions about the health history of the donors.

## Introduction

Single cell transcriptomic atlases are providing substantial new insights into human biology, including the molecular definitions of cell types, the cell type specific expression of disease-related genes, and the relationships of shared cell types across tissues, particularly those of the immune system^1–6^. These atlases are powerful reference tools whose applications in biology are just beginning to be explored. Many questions which could previously only be addressed at the bulk tissue level can now be answered with cell type specificity, thus yielding crucial new insights^6–8^.

Our previous efforts focused on creating a reference human cell atlas, called the Tabula Sapiens 1.0, which consisted of data from 24 different organs and tissues from a diverse set of donors^1^. In Tabula Sapiens 2.0, we have now doubled both the number of cells and the number of donors with multi-organ contributions, thus increasing the number of tissues and donors for which genetic background, age and epigenetic effects are controlled for. Specifically, we report data from nine new donors comprising one donor with 20 tissues, one donor with 18 tissues, one donor with 15 tissues, one donor with 2 tissues, and five donors with 1 tissue each. The updated combined Tabula Sapiens dataset contains more than 1.1 million cells from bladder, blood, bone marrow, eye, ear, fat, heart, kidney, large intestine, liver, lung, lymph node, mammary, muscle, ovary, pancreas, prostate, salivary gland, skin, small intestine, spleen, stomach, testis, thymus, tongue, trachea, uterus, vasculature. Single cell transcriptomes were obtained from live cells using both droplet microfluidic emulsions (1,093,048 cells) as well as FACS into well plates (43,287 cells).

To illustrate the utility of this reference dataset, we conducted several genome-wide expression studies. Our investigation mapped which transcription factors are active in specific cell populations, revealing cell-type associations for over a thousand regulatory proteins and uncovering previously unknown aspects of their functional contributions. Additionally, we examined aging-related cellular characteristics throughout our complete collection, which revealed which common gene expression modules used by all senescent cells alongside organ- and cell-specific aging patterns. We further explored how gene activity differs between males and females across every cell population in our study. This analysis revealed that biological sex influences gene expression primarily through cell-level regulatory mechanisms rather than through tissue-level differences. The data set is publicly available and we have also created an AI based tool to explore the clinical histories of the donors.

## Results

### Overview of Tabula Sapiens 2.0

The Tabula Sapiens 2.0 is an integrated map of 28 tissues collected across 24 donors. Nine new donors and four new tissues were collected and analyzed together with the original Tabula Sapiens 1.0 dataset (**Fig. 1, Supp. Fig. 1-Supp. Fig. 6**). All tissues and organs, with the exception of the respective reproductive organs, were profiled from both male (n=11) and female (n=13) donors. The donor’s age ranges from 22 to 74 years old, offering one of the most comprehensive molecular profiles of human tissues across the adult lifespan (7 donors under the age of 40, 11 donors between 40 and 60, and 6 donors over 60 years of age). A detailed table with metadata can be found in **Supp. Table 1**.

**Figure 1.**
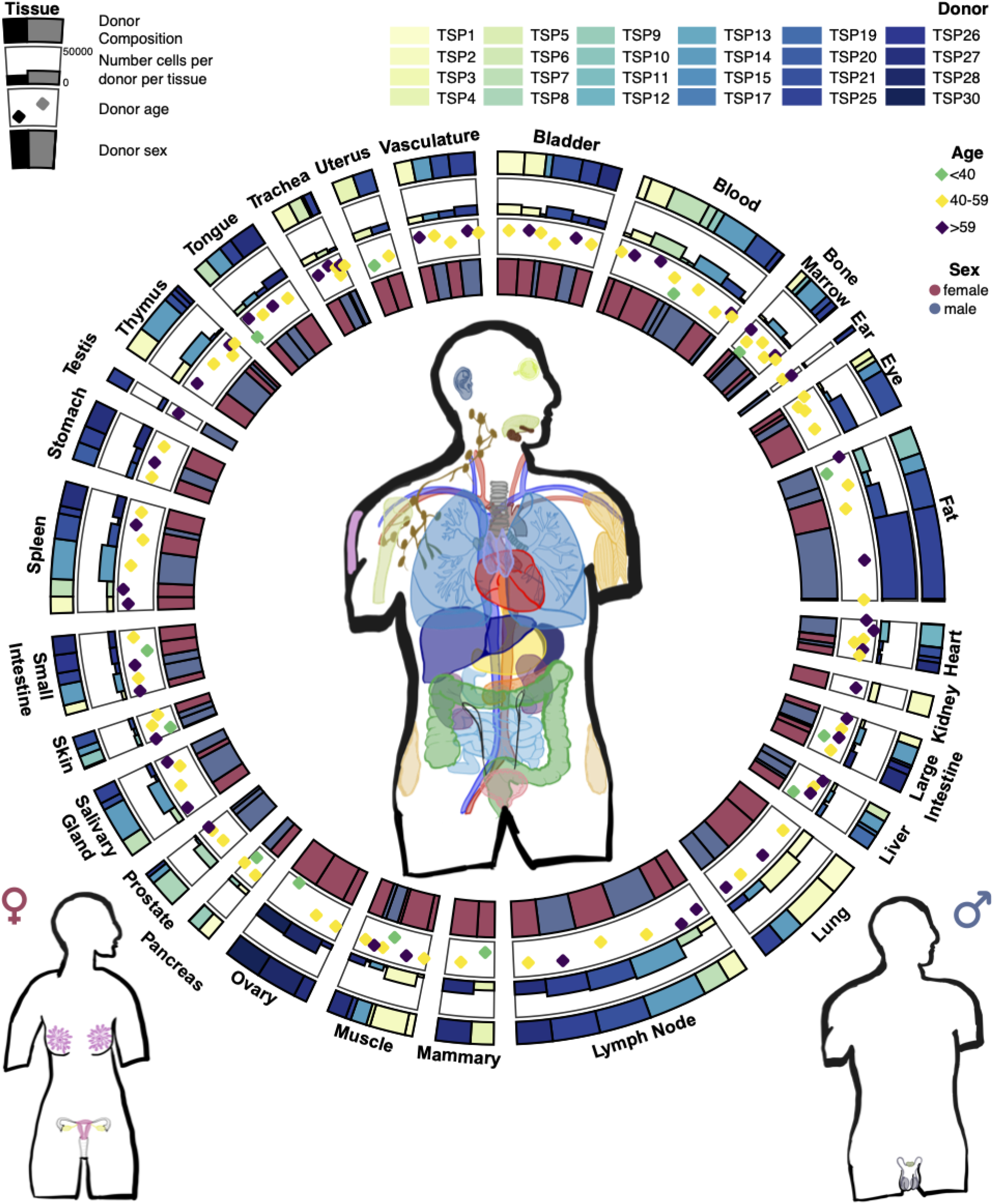
Overview of tissue donor demographics and tissue composition in Tabula Sapiens 2.0. The central anatomical illustration highlights the location of various tissues sampled from male and female donors, organized into distinct anatomical systems. Surrounding the anatomical figures are circular arrangements of charts showing the donor demographics of samples across various tissues. The inner stacked bar chart color-coded by donor sex for samples across every tissue (blue: male, pink: female). The diamond plot shows the donor age for samples across every tissue (green diamond: under 40, yellow diamond: 40-59, purple diamond: over 59). The bar plot shows the number of cells per donor for samples across every tissue with tissue color coded by donor ID. The outer stacked bar plot shows the donor composition for samples across every tissue. Tissues are labeled around the circular chart, providing an accessible visual summary of the tissue distribution and donor demographics.

For each organ and tissue, a group of experts used a defined cell ontology terminology to annotate cell types consistently across the different tissues, leading to a total of 701 distinct cell populations representing cell type combined with tissue of origin designation. The full dataset can be explored online using CELLxGENE^9,10^.

### Transcription factor analysis across cell types

A fundamental aspect of cell state and cell identity is the regulation of gene expression by transcription factors. Several large-scale consortia have systematically mapped human transcription factor (TF) activity at the tissue level. The ENCODE Project^11^ established standardized pipelines for genome-wide TF binding site identification using ChIP-seq and DNase-seq footprinting, generating high-resolution maps of regulatory DNA occupancy. The Roadmap Epigenomics Consortium^12^ expanded this work by creating reference epigenomic maps for 127 cell lines and tissues, integrating histone modifications, DNA methylation, and chromatin accessibility to contextualize TF function. The GTEx Consortium^13^ developed statistical approaches to infer individual-specific TF activities from RNA-seq data across 49 tissues, identifying genetic variants (QTLs) that modulate TF regulatory effects. Complementing these efforts, the FANTOM Consortium used deepCAGE technology to profile transcription initiation events and reconstruct TF interaction networks governing cellular differentiation. While these foundational studies focused on bulk tissue methods, more recent work has used TF overexpression in single cells to reveal insights into TF-driven human developmental trajectories^14^.

Of the 1639 putative transcription factors (TF) in The Human Transcription Factors database^15^ (HTF) we investigated the 1637 TFs which appear in GRCh38 gene annotations. We computed the mean of the log-normalized expression of each transcription factor across the 175 cell types detected in our droplet microfluidic dataset. All but two TF’s are expressed in at least one cell type in this data set. SHOX and ZBED1, both from the pseudoautosomal region 1 of the X and Y chromosomes, have zero raw droplet counts in every donor. The Human Protein Atlas^16–18^ (HPA) reports RNA and protein detection of SHOX in a broad range of tissues with enrichment in basophils, as well as expression in fibroblast cell lines. ZBED1 is found in a small number of cells across all cell types in our plate data and the HPA also reports ubiquitous cell type expression. We therefore have been able to establish the cell type specificity of all known human transcription factors.

In order to test whether gene expression of these transcription factors was also accompanied by activity, we used the SCENIC software package to infer TF activity in single cells of each cell type separately. SCENIC identifies TF’s target genes through co-expression-based GRN inference algorithms and presence of binding motifs in proximity to target genes. Activity of these TF regulons is then scored by enrichment of the identified target gene set in each cell. Of 1390 TFs that met SCENIC’s entry criteria we found 839 regulons in at least one cell type. The remaining 551 did not survive SCENIC’s filters at various stages; this does not mean they didn’t have significant co-expression with others genes but that their co-expression was below an internal threshold. Across the 38 broad cell classes, SCENIC identified varying numbers of active TF regulons (mean=80.3, sd=59.6). Individual regulons were found in up to 31 cell types with most regulons, however, showing high cell type specificity, being found on average in 3.6 cell types (sd=4.0) **(Supp. Fig. 7)**. Overall this provides evidence of activity by downstream gene expression for roughly half of the human TF’s and does not rule out activity for the remainder.

To establish transcription factor cell type and tissue specificity, we computed the specificity statistic τ ^19,20^ across all donors **(Methods, Supp. Table 2)**. When computed on 5 individual samples we found r = 0.76 to 0.81 sample-to-sample correlation (**Supp. Fig. 8**). This statistic was developed for microarray and bulk tissue RNA-seq studies, and has recently been used to determine cell-type specificity in single-cell mRNA-seq data studies such as the *Drosophila* cell atlas^21^ and a mouse brain atlas^22^. As seen in bulk mRNA-Seq studies^13^, we found the τ distribution dips at τ ≅ 0.85. This same 0.85 dip occurs in the *Tabula Muris Senis* and *Drosophila* single cell atlas datasets; and also when τ is computed on a tissue basis or on a broad cell type basis in our data. Thus we defined the 890 TF’s above this as cell type specific **(Fig. 2A)**. We defined the 745 TF’s with τ < 0.85 as non-specific and used the 100 lowest τ TF’s to explore the space of ubiquitous TF’s. To visualize TF expression we placed the TFs in τ order and grouped expression values by 38 broad cell class categories to create an expression heatmap (**Fig. 2B)**.

**Figure 2:**
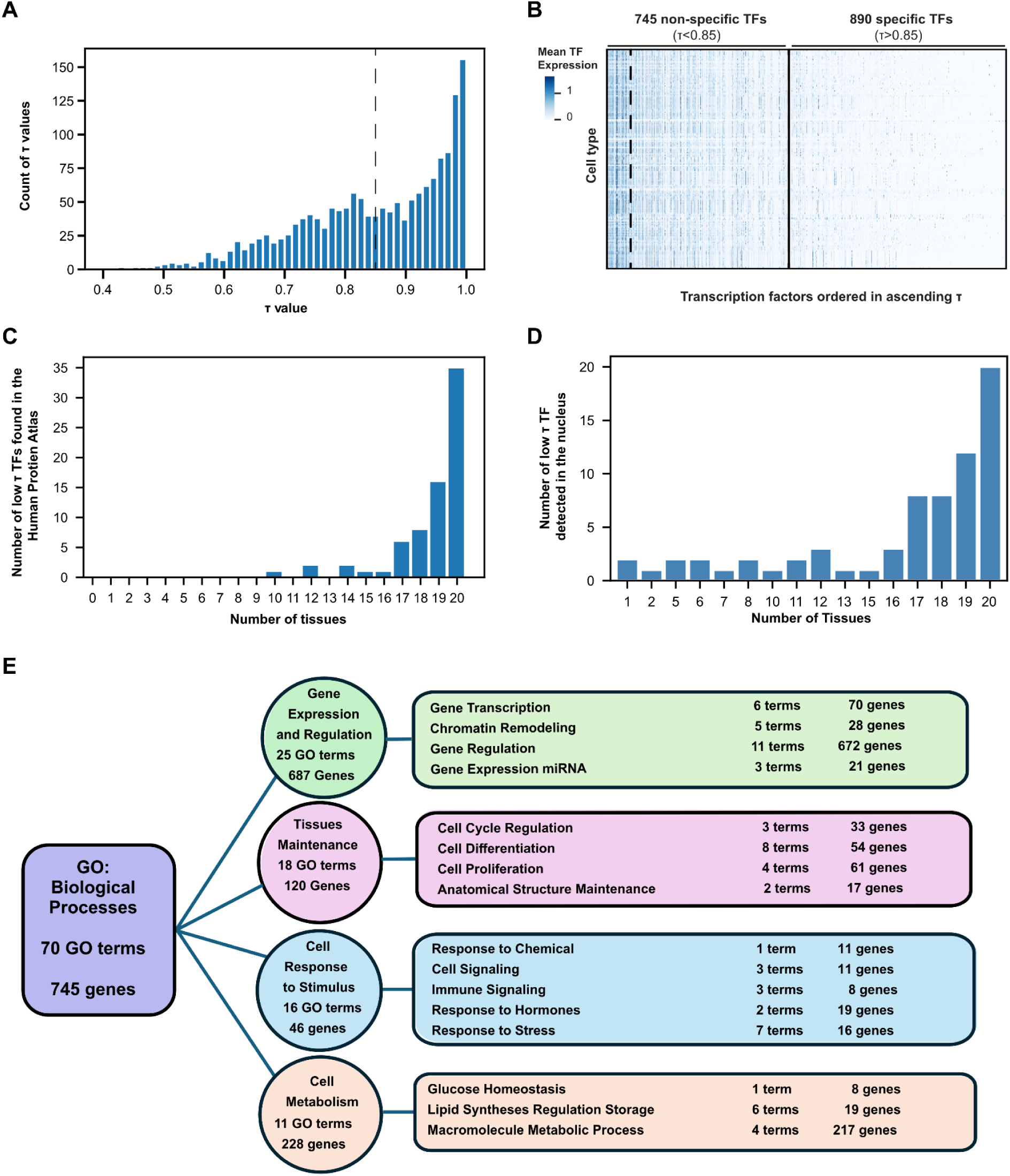
Transcription factor expression common across cell types. A) Histogram of the distribution of τ values for 1635 transcription factor genes computed across 175 distinct cell types in 28 Tabula Sapiens 2.0 tissues, marked with specificity threshold at τ > 0.85. B) Heatmap of the mean of log normalized expression in each cell type of 1635 TF’s in the 175 cell types. The mean expression values range from 0.0 to 4.9 but for better visualization in the heatmap they have been clipped to the 99.0 percentile resulting in the heatmap scale from 0 to about 1.0. Cell types are in alphabetical order by broad cell type. Transcription factors are arranged in τ value order from from 0.4 to 1.0 τ. The vertical bars on the heatmap mark the lowest 100 τ ubiquitous TF’s, 745 τ < 0.85 non-specific TF’s and the 890 τ > 0.85 cell type specific TF’s. C) Distribution of the 72 ubiquitous sc-mRNA expressed TF’s detected as expressed proteins by immuno-staining in 20 common tissues in the Human Protein Atlas. D) Distribution of nuclear expression of 69 ubiquitous TF’s across 20 tissues. E) Summary of the Enricher GO analysis of the 745 non-cell type specific transcription factors outlined in black in A. Enrichr returned 70 GO terms that were organized by broad cell function categories shown in the circles with the number of GO terms and genes in each category. Sub-categories are shown in the rectangles with the number of GO terms and genes in each.

Among the 100 TF’s labeled as ubiquitous, we found many well studied factors known to be central to universal cell function such as stress response, proliferation and metabolism. Examples of these include *NFAT5*, *NCOA1*, *FOXJ3*, *FOXK2*, *ATF4*, *JUN*, *FOS* and *STAT1.* To validate the existence and roles of the ubiquitous TF’s, we searched the HPA for evidence of protein expression. Of the 100 ubiquitous TF’s, 72 were tested in the HPA; every single one of these was detected in at least one tissue and the majority (71%) were detected in 19 or 20 of the 20 tissues tested. (**Fig 2C, Supp Fig. 9A)**. In the 32 cell types, all 72 low τ TF’s tested were detected in many cell types. The fraction detected in more than 80% of cell types was 0.69, vs 0.40 for all 699 TF’s. **(Supp Fig. 9C-D).** To further confirm the likelihood these TF’s are active, we queried the HPA for the subcellular location of expression of the 72 ubiquitous TF’s; the HPA detected expression in the nucleus for 69 of these across a broad set of tissues Altogether there is evidence of protein expression for 100% of the ubiquitous TFs found in the HPA and evidence of activity based on nuclear localization for 96% of them (**Fig 2D**).

We explored the biological function of the full set of 745 non-specific TF’s (τ < 0.85) by performing Gene Set Enrichment Analysis (**Fig. 2E, Methods, Supp. Table 3)** and grouping the GO terms into four broad cellular function categories: Gene Expression and Regulation, Tissue Maintenance, Cell Response to Stimulus, and Cell Metabolism. Within the Gene Expression and Regulation category, we found 25 Gene Ontology (GO) terms and sub-grouped these into gene transcription, gene regulation, gene expression of miRNA and chromatin remodeling as needed for the cell to synthesize mRNA and miRNA. Gene regulation comprises 11 GO terms and dominates the number genes driving these terms, 672, by an order of magnitude.

Transcription factors for maintenance of tissues are enriched in our data in 18 GO terms. We grouped these into cell cycle regulation, cell differentiation, cell proliferation and anatomical structure maintenance. Cell cycle regulation enriched for 33 genes in three GO terms. These genes include *TFDP1* and *TFDP2* which form complexes with E2F transcription factors to control cell cycle^23,24^ and *E2F6* which specifically blocks cell cycle entry^25^. The cell differentiation group includes the two broad terms negative regulation of cell differentiation (GO:0045596) and positive regulation of cell differentiation (GO:0045597), as well as terms identified as specific for a variety of cell types. There are 61 transcription factors enriched in the cell differentiation category. Of these, *ARNT*, *ARNTL*, *CEBPB*, *CREB1*, *CREBL2*, *ETS1*, *FOXO1*, *FOXO3*, *HIF1A*, *PPARD*, *SMAD3*, *STAT1*, *STAT3*,*STAT5B*, *XBP1*, *ZBTB16*, *ZFPM1* and *ZNF16* are enriched in three or more terms. Cell proliferation has four GO term a): regulation of transforming growth factor beta2 production (GO:0032909), b) hemopoiesis (GO:0030097), c) regulation of cell population proliferation (GO:0042127), and d) positive regulation of myoblast proliferation (GO:2000288). These terms have enriched genes, including the 3 ATP-1 binding factors *JUN*, *JUNB*, and *JUND*; 6 STAT family transcription factors (which respond to growth factor receptors) and *KLF4*, *KLF10*, *KLF11* (which can negatively regulate cell growth). The anatomical structure maintenance group contains two GO terms with 17 enriched transcription factors. These include all 3 PBX TF’s of the TALE family that interact with the HOX domain^26^; and 3 of the 4 TEAD factors known in humans to regulate tissue size and function by coupling external signals to control cell growth and specification^27^.

Response to a cell’s environmental stimuli is found in five groups of GO terms: chemical signaling, cell signaling, immune signaling, hormone response, and general stress response. Cell response to stress encompasses responses to fluid stress, laminar flow stress, oxidative stress, hypoxia, and chemical stress. Note, some transcription factors may be induced beyond natural levels due to processing tissue into live single-cells for analysis. Among 16 stress response genes, *ATF3*, *ATF4*, *ATF6*, *CEBPB*, *DDIT3*, *HIF1A*, *KLF2*, *KLF4*, *NFE2L2*, *RBPJ*, and *TP53* are enriched in association with two or more GO terms. *KLF2* and *KLF4* are suggested to be part of the Golgi stress response^28^. Also enriched is *SREBF2*, which regulates cholesterol synthesis in response to a broad set of stresses including inflammation, heat shock and hypoxia^29^. Oxidative stress response is thought to be regulated by enriched *NFE2L2* (NRF2)^30^ through the KEAP1-NRF2-ARE pathways^31^, as well as by *HIF1A*^32^. GO terms for cell signaling fall into three groups, cell signaling, IL6 and IL9 immune signaling, and peptide hormone signaling. They share the common STAT signaling pathway whereas cell signaling selectively utilizes NFAT TF’s and hormone signaling utilizes NCOA TF’s^33^.

Within the 10 GO terms enriched for cell metabolism four involve regulation of macromolecule synthesis, five terms regulate lipid synthesis, and one maintains glucose homeostasis. The four macromolecule synthesis terms include 217 TF’s, more TF’s than any set of terms other than gene regulation.

Within the non-specific TFs we observed many widely expressed zinc finger DNA binding family TF’s. Of these, the Human Transcription Factor Database shows many are only computationally predicted, or have no defined motif, and lack experimental validation. Often the literature only mentions differential expression in the context of disease, with no known function. It is striking that so many broadly expressed transcription factors exist and are not well characterized, including many that are ubiquitous across cell types..

We then examined transcription factor expression levels for the 890 cell type specific TF’s (τ>0.85). We discovered both previously known as well as novel relationships between specific TF’s and the cell types in which they are expressed (**Fig. 3A)**. Each TF was grouped with the cell type of its highest mean expression and plotted in broad cell class alphabetical order. Vertical lines show the ubiquitous TF’s are redistributed throughout cell types, while the horizontal bars reveal highly expressed TFs in each cell type or in a broad cell class. This provides putative cell type association for the specific TF’s, many of which previously have only been identified within the context of bulk tissue expression.

**Figure 3:**
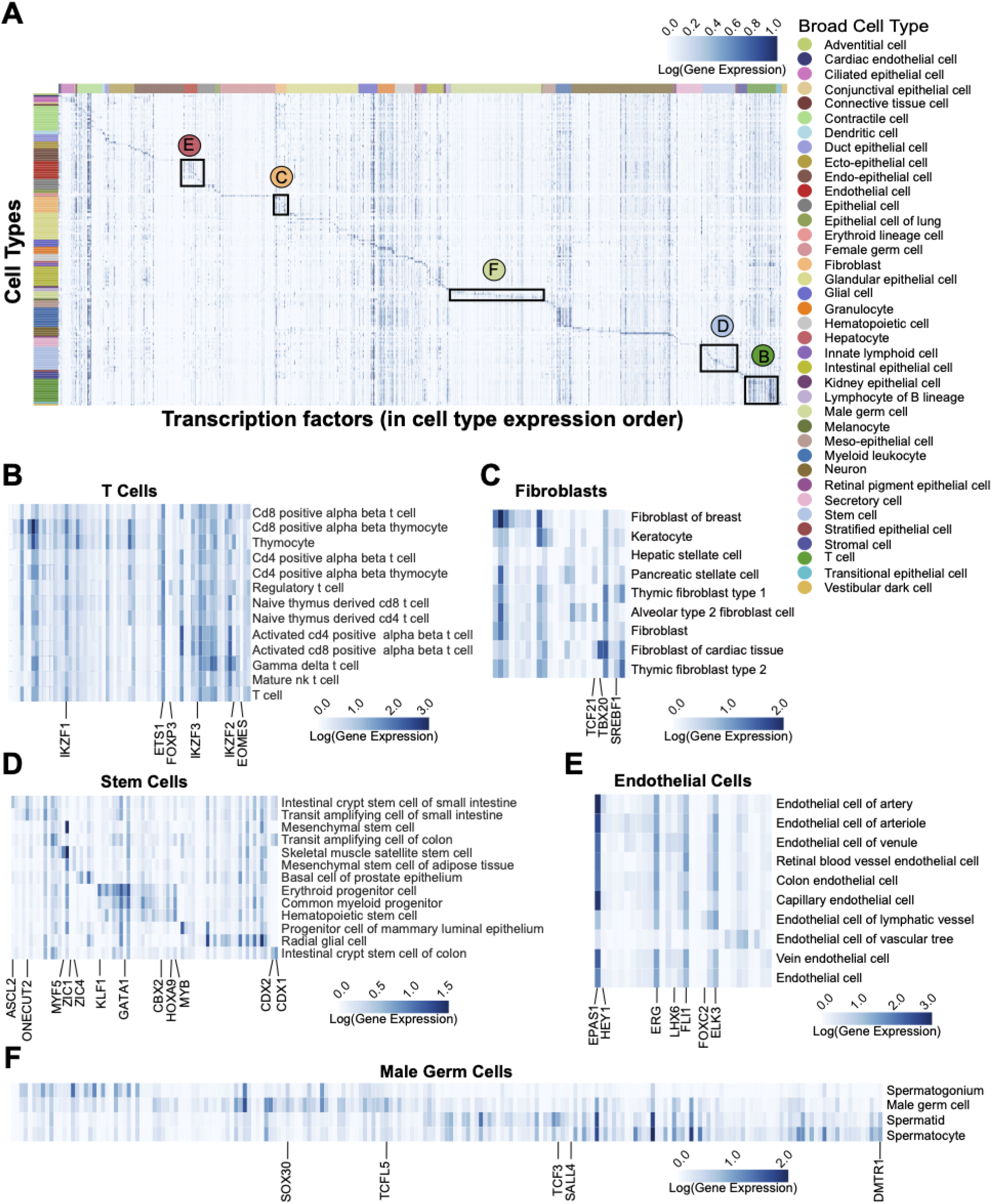
Transcription Factors Specific to Cell Types. A) Expression heatmap showing the mean of the log normalized expression values of 1635 human transcription factors in 175 cell types using all 28 Tabula Sapiens 2.0 tissues. For better visualization in the full heatmap (A) the max values are clipped at the 99.5 percentile. The cell type rows are organized alphabetically by broad cell type and denoted by color on the annotation bar. The TF columns are arranged according to the cell type for which they show the highest expression. Black outlines highlight examples of common broad cell types and refer to sub-figures B-F, which are not clipped at the 99.5 percentile in order to show the full range of expression. B) T-Cells. C) Fibroblasts. D) Stem Cells. E) Endothelial Cells. F) Male Germ Cells.

Within the T-cell broad cell class we identify five TF’s with τ > 0.85 *FOXP3*, *EOMES*, and three Ikaros TF’s (*IKZF1*, *IKZF2*, and *IKZF3*), suggesting they are specific to T-cells (**Fig. 3B**). *FOXP3* expression is the defining factor for *FOXP3* Treg cells^34^. In these cells it coordinates with *ETS1*, which we see expressed across all T-cell types^35^. *EOMES* highest expression is in Mature NK T cells and CD8 positive alpha beta T cells^36^. The three Ikaros family TF’s are well known to regulate all types of lymphocyte cell development^37,38^. We find them expressed throughout T-cell types.

Gene expression profiles in fibroblast cells are heterogeneous and often more organ or environment specific than cell type specific^39–42^. Thus, over 80% of cells in this broad category have been typed only generally as ‘fibroblast cell’ (**Fig. 3C**). We do find three fibroblast/organ specific TF’s where we have specific fibroblast cell types. In the lung, *TCF21* has highest expression in alveolar type-2 fibroblast cells where it regulates lipofibroblast differentiation and serves as a marker for these cells^43^, *TBX20* plays a key role in development and maintenance of healthy cardiac tissue^44^ and *SREBF1* expression in thymic fibroblasts is known to be required for activated T-cell expansion^45^.

Our data show nine stem cell type specific TF’s involved in the generation of fat, bone, blood, and gut cells (**Fig. 3D**). *ZIC1* and *ZIC4* have their highest expression in mesenchymal stem cells of adipose tissue, where *ZIC1* is known to direct stem cell fate between osteogenesis and adipogenesis^46^. We also see *ZIC1* and *MYF5* expressed in skeletal muscle satellite stem cells where they regulate myogenesis^47^. *ZIC1* is suggested to be involved in brown fat precursor differentiation or in white-to-brown transdifferentiation^48^.

Gut stem cells are known to be maintained by *CDX2* and *CDX1*^49^, and these two regulatory TF’s have τ > 0.85 in our data, with their highest expression found in transit amplifying cells and intestinal crypt stem cells in both the small intestine and colon. Specific to small intestine crypt stem cells we find expression of *ASCL2*, a master controller of intestinal crypt stem cell fate^50^. *ONECUT2*, a downstream target of *ASCL2*, has its highest expression in transit amplifying cells and crypt stem cells. However, *ONECUT2* expression is not restricted to the broad stem cell type in our data as it has similar expression levels in all the small intestine enterocyte cell types. *ONECUT2* is also suggested to regulate differentiation of M cells of Peyer’s patches^51^. Three TF’s with τ > 0.85 have their highest expression in hematopoietic stem cell types. Consistent with the literature, *MYB* is found in decreasing amounts in hematopoietic stem cells, then myeloid progenitors and then erythroid progenitors. It also shows some expression in the intestines^52^. It is worth noting that genes most famous for their roles in early development, such as the Homeobox family and the Yamanaka factors, are also known to be expressed in adult cell types and are present in our data. For example *HOXA9* and *CBX2*, necessary for embryonic development and hematopoiesis^53–55^, show the decreasing expression from hematopoietic stem cell type to progenitor cell types. We also see TF’s *KLF1* and *GATA1* increasing expression in sequence towards development of erythroid progenitor cells^56,57^.

The broad endothelial cell class heatmap (**Fig. 3E)** reveals 18 TF’s with > 0.85 τ. Of these, *HEY1*, shows higher expression in the arterial vascular cell types where it may mediate arterial fate decisions^58^. Similarly, *LHX6* has been extensively studied in neuron development^59^, but in our data is expressed in vein and venules endothelial cells suggesting a regulatory role in venous fate. *FOXC2* is expressed highest in lymphatic cells where it is known to regulate formation of the lymphatic system^60^. Together these TF’s appear to regulate the fate of all the three types of vascular structures, arterial, venous and lymphatic. Finally, we see *ERG* expression known to stabilize vascular growth through regulation of VE-caderhin^61^ and known to maintain vascular structure by repressing proinflammatory genes^62^. We see several less cell type specific TF’s with high expression which are also known to regulate stability of the vascular tree. *FLI1* and *ERG* prevent endothelial to mesenchymal transition^63^. *ELK3* suppresses further angiogenesis^64^. *EPAS1* (HIF2A) regulates physiological response to oxygen levels. Interestingly, some *EPAS1* alleles contribute to high altitude athletic performance^65,66^.

Finally, we highlight male germ cells which have 204 TF’s with their highest expression in the four cell types in this broad cell class (**Fig. 3F**). In addition, 50% of these are > 0.85 τ testis specific, consistent with findings in *Drosophila*^21^. Over 50% of these specific TF’s begin with the letter ‘Z’ indicating a zinc finger motif. This appears to be a rich area for further study as most of these TF’s are computationally derived and/or understudied, especially in humans, as most work has been done in mice. Spermatogenesis is driven by specific TF’s through four stages, going from male germ cell to spermatogonia via mitosis, then to spermatocyte and spermatid through meiosis, to become sperm^67^. As is known, our data shows specific expression of *SALL4* in male germ cells where it maintains this pool of cells^68,69^. *DMRT1* is known to specifically coordinate male germ cell differentiation and our data shows its expression growing from male germ cells to the spermatogonia phase^70,71^. Spermatogonia proliferation is then driven by expression of *TCF3*^72^. *SOHLH1* is regulated by *DMRT1* and known to be active in the later stages of spermatogonia mitotic division^71^. Although in our data, *SOHLH1* has its highest expression in oocytes, we see it decreasing as male germ cells progress to spermatogonia (**Supp. Table 4).** Spermatogonia then execute major transcriptional changes to become primary spermatocytes where they progress through two rounds of meiotic division to become spermatids. This requires a complex set of transcriptional changes, which are known in mice to be coordinated by *TCFL5* and *MYBL1*^73^. In our data *TCFL5* has its highest expression in spermatocytes. *MYBL1* has a reasonably high τ value of 0.95 but it has higher mean expression in several immune cell types. Outside of immune cells, its highest expression is in spermatocytes (**Supp. Table 4**). Finally, transition from spermatocyte to early spermatid is known to be initiated by expression of specific TF *SOX30*^73,74^. In our data *SOX30* expression is highest in spermatocytes and also high in spermatids.

### Molecular profiles of senescent cells across human tissues

Cellular senescence is a state of irreversible cell cycle arrest, which can be triggered by various cellular and environmental stresses^75,76^. At a molecular level, cyclin-dependent kinase inhibitors, particularly p16-INK4a (encoded by the gene *CDKN2A*), play a crucial role in controlling the initiation and maintenance of cellular senescence^77,78^. At a cellular level, senescence has been characterized by its unique features such as enlarged cell morphology, increased senescence-associated beta-galactosidase activity (SA-β-Gal), formation of senescence-associated heterochromatin foci (SAHF), and production and secretion of inflammatory cytokines, growth factors, and proteases, known as senescence-associated secretory phenotype (SASP)^79,80^. Nonetheless, the phenotype of cellular senescence is highly variable and heterogeneous, with mechanisms not universally conserved across all senescence programs^81–87^. The accumulation of senescent cells has been shown to affect various aspects of aging^88^ and their clearance has been shown to delay various age-related degenerative pathologies^89,90^.

One of the key challenges in studying senescence is its heterogeneity. Studies using senescent cell culture models have demonstrated this variability, showing that cells can exhibit diverse phenotypes depending on the cell type and the specific inducer of senescence^84,85^. For instance, SASP has been shown to vary based on cellular context^86,87^. This *in vitro* heterogeneity points to the possibility that senescent cells in living organisms are similarly diverse, with distinct cell phenotypes. Compounding the issue, many classical markers of senescence, which were established using cell culture models, have limited utility *in vivo*. To tackle this issue, researchers have developed guidelines such as the Minimum Information for Cellular Senescence Experimentation in Vivo (MICSE) and the SenNet guidelines for identifying senescent cells in human tissues. These provide recommendations for evaluating senescence markers directly within tissues and in live animal models^91^.

Given the heterogeneity of senescence markers and the methodological challenges in detecting them, we defined senescent cells based on the expression of *CDKN2A* and the absence of *MKI67* (a cell proliferation marker). To deploy a multi-pronged approach for identification of senescent cells as suggested in SenNet guidelines, we performed a differential expression analysis for genes representing canonical hallmarks of senescence in senescent cells as compared to non-senescent cells across all broad cell types in Tabula Sapiens 2.0. Although our operational definition of senescence drew exclusively on the cell-cycle-arrest hallmark (*CDKN2A*+ and *MKI67*-), the resulting differential-expression profile revealed that at least one hallmark-specific gene exhibited the expected direction of change for at least two additional hallmarks in most cell types (**Supp. Fig. 10A**). Thus, the *CDKN2A*+ *MKI67*- population not only satisfies the cell-cycle arrest criterion but also displays molecular signatures consistent with multiple complementary hallmarks, aligning with the framework proposed by SenNet.

To systematically characterize the phenotype of senescence cells of different cell types across various human tissues, *CDKN2A*+ *MKI67*- cells were identified across 25 tissues from 21 donors in the Tabula Sapiens 2.0 dataset (**Methods**). We detected 48,114 senescent cells (∼4.4% of all cells) spanning 145 cell types grouped into 34 broad cell type categories^92^ (**Supp. Fig. 11A & 11B**). This represents the largest and most comprehensive study of senescent cells in human tissues to date, providing an unprecedented view of their distribution and diversity across multiple tissue types and cellular contexts. We detected a small increase in senescent cells from medium (40-60y) and old donors (>=60y) as compared to young donors (<40y) (**Fig. 4A**). As expected, we observed a heterogeneity in the burden of senescent cells across different tissues, with eye, bladder, and tongue containing the highest, and the heart, muscle, and ovary containing the lowest proportion of senescent cells (**Fig. 4B**). The low burden of senescent cells in the heart, quadriceps muscle, and gastrocnemius has been observed previously^93^. Comparing the proportion of senescent cells in non-reproductive organs between male and female donors, we observed a slightly higher senescence cell burden in the spleen and lungs from female donors as compared to male donors (**Fig. 4B**). Next, we looked at canonical markers associated with the cellular senescence phenotype in our dataset. In addition to *CDKN2A*, previously known senescence-associated DNA damage marker *H2AX*, SASP regulators such as *TGFB1* and *NFKB1*, and SASP markers such as *HMGB1* and *TIMP2* were also enriched in senescent cells as compared to non-senescent cells (**Supp. Fig. 11C**).

**Figure 4:**
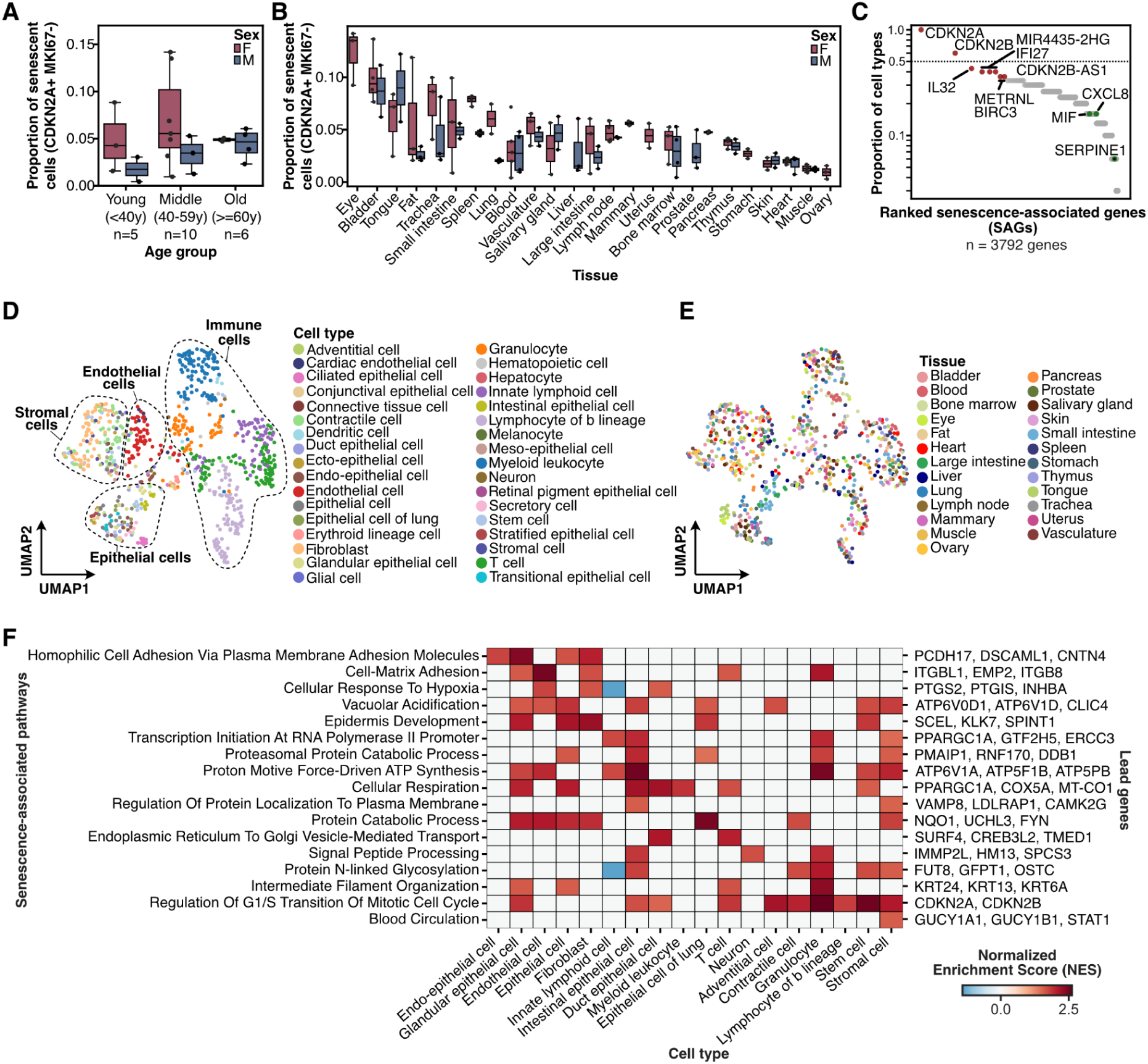
Molecular profiles of senescent cells across human tissues. **A)** Box plot showing the proportion of senescent cells, defined as *CDKN2A*+ *MKI67*- cells, across donors from three age groups. The donors within each age group were split by sex. The error bars represent the 95% confidence interval. **B)** Box plot showing the proportion of senescent cells, defined as *CDKN2A*+ *MKI67*- cells, across 25 tissues from 21 donors. The donors within each age group were split by sex. The error bars represent the 95% confidence interval. **C)** Dot plot showing the cell type prevalence for 3792 senescence-associated genes (SAGs). Cell type prevalence represents the number of cell types where a gene was upregulated in senescence cells as compared to non-senescent cells of that cell type from any tissue. A gene is considered to be upregulated in a cell type if the mean log_2_ fold-change is greater than 0.5. The top 15 most universal SAGs are highlighted in red, while known senescence-associated secretory phenotype (SASP) genes are highlighted in green. **D-E)** A Uniform Manifold Approximation and Projection (UMAP) plot showing pseudo-bulked transcriptomes of senescent cells, averaged for donors, tissues, and cell type combinations, and clustered based on the expression of senescence-associated genes alone. **D)** The pseudo-bulk transcriptomes are colored according to the cell type they represent. **E)** The pseudo-bulk transcriptomes are colored according to the tissue of origin. **F)** Heat map showing normalized enrichment scores (NES) for senescence-associated pathways across broad cell types. Enrichment scores were derived from gene set enrichment analysis using ranked enrichment between senescent and non-senescent cells for each broad cell type. Senescent associated pathways were filtered for FDR < 0.05 to retain pathways significantly enriched in at least one broad cell type.

To understand the universality of senescence phenotype, we first identified genes that were upregulated in senescent cells across individual tissues and cell types, and used these genes to quantify the cell type and tissue prevalence of senescence-associated genes (SAGs) (**Methods, Supp. Fig. 11B**). To find SAGs for individual cell types, we selected genes significantly upregulated in senescence cells of that cell type from any tissue in at least 50% of the donors (log_2_ fold change > 0.5, adjusted p-value < 0.001, **Supp. Table 5**). We ranked the 3792 SAGs, which were upregulated across 30 broad cell types in 25 tissues based on their cell type prevalence (**Methods, Supp. Fig. 10B**). Interestingly, after *CDKN2A*, *CDKN2B* was the most universal SAG, which was upregulated in ∼60% of the cell types (i.e. 18 out of 30) (**Fig. 4C**). Surprisingly, SASP markers such as *IL6* and *IL1B*, as well as regulators such as *NFKB* and *TGFBI*, did not meet our selection criteria and were therefore not included as SAGs. Among the known SASP factors, the neutrophil-attracting chemokine *CXCL8*, as well as the pro-inflammatory cytokine *MIF*, were enriched in senescent cells for ∼16% of the cell types (i.e., 5 out of 30), and *SERPINE1* was enriched in senescent cells of only two cell types (**Fig. 4C**). Gene set enrichment analysis of SAGs revealed an enrichment in ontology terms associated with cellular respiration, mitochondrial translation, protein transport, and regulation of apoptotic processes **(Supp. Fig. 11D)**.

The most universal SAGs detected in this study include the gene *IL32*, encoding for the proinflammatory cytokine Interleukin-32. *IL32* h was found to be enriched in senescent cells across 43% of the cell types is present in higher mammals but not in rodents, and has recently been shown to trigger cellular senescence in cancer cells^94^. This suggests that Interleukin-32 could be an important mediator of paracrine-induced senescence across multiple human tissues (**Fig. 4C**). Interferon stimulated gene *IFI27* was also enriched in senescent cells across 40% of the cell types (**Fig. 4C**). *IFI27* also known as *ISG12a* is known to possess potent anti-proliferative and tumor suppressive properties and has been reported to be enriched in senescent cells in models of down syndrome and Alzheimer’s disease^95^. Additionally, *IFI27* has also been shown to show a strong and sustained expression during TNFα-induced senescence^96^. Other genes that were enriched across multiple tissues include *BIRC3* gene, which not only regulates apoptosis, but also modulates inflammatory signaling, mitogenic kinase signaling, and cell proliferation (**Fig. 4C**). *BIRC3* gene expression has previously been described as a potential survival factor for senescent glioma cells (GBM), where targeting the product of *BIRC3* gene was shown to act as a senolytic strategy that triggered apoptosis of GBM cells^97^.

Collectively, these results point towards the absence of a fully universal senescence program, which has been suggested recently by others^81,83^. Nonetheless, we sought other broadly expressed markers that might reflect more common senescence features. To gain a more comprehensive view of senescence phenotypes across various cell types and tissues, we clustered the transcriptomes of senescent cells from various donors, tissues, and cell types using only the 3792 SAGs. We observed that this set of SAGs resulted in the senescent transcriptomes clustering together largely by related cell types, and independently of the tissue of origin or the donor (**Fig. 4, D and E**). This indicates that while there is no universal senescence phenotype, there are shared effects which are broadly shared across certain cell types. To systematically characterize the heterogeneity of senescence phenotypes across all cell types, we identified senescence-associated gene modules co-expressed within the same cell types, tissues, and donors, and summarized enrichment scores for these phenotypes for all cell types (**Methods, Supp. Fig. 11B**). Seventeen gene expression modules, each representing distinct biological processes, were variably enriched across different cell types (FDR q-value < 0.05, **Fig. 4F**), and it appears that the various different forms of human cellular senescence can be explained by combinations of these programs, along with some degree of cell-type specific changes.

Gene set enrichment analysis of senescence-associated pathways revealed that senescence programs are notably modulated by cell types and their lineage context. We found that physical cohesion pathways, such as strong homophilic cell-adhesion driven by protocadherin genes such as *PCDH17* and *DSCAML1*, were enriched in senescent barrier and structural cells such as epithelial cells and fibroblasts (**Fig. 4F**). Similarly, genes associated with epidermal development such as *SCEL*, *KLK7*, and *SPINT1* were enriched in senescent epithelial cells and fibroblasts (**Fig. 4F**). Cell-matrix adhesion-related genes, such as *ITGB1* and *ITGB8*, which encode β-integrins, were significantly enriched in senescent epithelial cells, fibroblasts, and endothelial cells. (**Fig. 4F**). Changes in production of cell adhesion molecules including β-integrins have been described in senescent cells before^98,99^. Interestingly, genes related to transcription initiation at RNA polymerase II promoter pathway, such as *PPARGC1A*, *GTF2H5*, and *ERCC3,* were found to be upregulated in senescent innate lymphoid cells and granulocytes, likely representing a compensatory response to meet the enhanced transcriptional demands of energy-intensive SASP production, metabolic reprogramming, and the maintenance of cellular viability despite growth arrest^100^ (**Fig. 4F**). Secretory-pathway pressure was further underscored by the selective enrichment of ER-to-Golgi vesicle transport genes, such as *TRAPPC3* and *VCP*, in senescent intestinal epithelial cells and signal-peptide processing genes, such as *IMMP2L* and *HM13*, in senescent neurons and granulocytes. Together with the strong proteasomal and general protein-catabolism scores seen in granulocytes, lung epithelium, and stromal cells, these data reaffirm the centrality of proteostatic stress in senescence^101,102^.

We found that metabolic rewiring pathways were variably enriched in senescent cells across a wide variety of cell types. Genes associated with proton motive force-driven ATP synthesis as well as broader cellular respiration, such as *ATP6V1A*, *ATP5F1B, MT-CO2*, and *COX7A2* were enriched in senescent cells of intestinal epithelium, myeloid leukocytes, and stem cells, echoing mitochondrial dysfunction that couples bioenergetic decline to SASP induction^103,104^ (**Fig. 4F**). These results align with the growing appreciation that senescent cells can up-regulate oxidative phosphorylation to sustain the energy-intensive senescence-associated secretory phenotype (SASP)^105,106^. Mitochondrial maintenance thus appears not to be a bystander but an active participant across immune, epithelial, and progenitor compartments. Genes involved in hypoxia responses such as *PTGS2*, *PTGIS*, and *INHBA* were predictably enriched in endothelial and fibroblast populations that dwell in oxygen-gradient niches.^107^ (**Fig. 4F)**. Vacuolar acidification associated genes such as *ATP6V0D1* and *ATP6V1D* were enriched concurrently in endothelial, epithelial, stromal and stem cells, reinforcing that enlarged, hyper-acidic lysosomes are a hallmark of senescence, and their prominence across both differentiated and progenitor cell types further cements the universality of lysosomal remodeling in senescence^108^. Lymphoid cells showed an enrichment for the G1/S cell cycle arrest, with increased *CDKN2A* and *CDKN2B* enrichment paralleling the accumulation of p16^+^ immune cells that evade clearance during ageing. In contrast, myeloid cells showed high MiDAS and proteasome activity related genes, implying a hyper-metabolic, SASP-potent phenotype distinct from the quiescent lymphoid state (**Fig. 4F**).

Collectively, we discovered distinct and cell type-specific fates for these cells that undergo cell cycle arrest. This constellation of pathways underscores that cellular senescence is a multifaceted state combining cell-cycle arrest with strategic rewiring of adhesion, metabolism, proteostasis, and organelle function, tailored to each cell types’s role in tissue homeostasis and age-related pathology. Our results show that while there is not a single type of senescence, there are broad programs shared by several classes of cell types across tissues.

### The impact of sex on gene expression across tissues and cell types

With the comparative sex information from male and female donors in the Tabula Sapiens 2.0 dataset, we systematically investigated differences in gene expression across a wide range of cell types. Sexual dimorphism in gene expression has been well-documented at the tissue and species level^109–113^, with implications for differential disease susceptibility, drug responses and biological processes between males and females^114–118^. These differences are influenced by variations in sex chromosomes, hormonal signaling, transcription factor activity, and environmental influences^119^. One classic example of sex-dependent gene expression is the *XIST* gene^120^, which plays a key role in X chromosome inactivation in females, ensuring dosage compensation between sexes. Other X-linked genes, such as *DAX1*^121^, *KDM6A*^122^ exhibit sex-specific expression patterns that impact processes like histone modification, gene regulation, and reproductive tissue function. Y-linked genes such as *SRY*^121^ and *TSPY*^123^, are crucial for male sex determination and spermatogenesis respectively. These genes are exclusively expressed in males. Recent studies also underscore tissue-specific gene expression contributing to sex differences. For example, genes like *CYP3A4* and *STAT3* are more highly expressed in females in the liver, affecting drug metabolism and pharmacokinetics^124^. Additionally, *LEP*, with higher expression in females, regulates energy balance, appetite, and fat distribution^125^, while differences in *PPARG* expression between sexes influence immune responses, including the regulation of inflammation and T cell differentiation, in addition to its roles in adipogenesis, insulin sensitivity, and lipid metabolism^126,127^.

Despite progress in understanding sex effects on gene expression across tissues, gaps in understanding remain at the cellular level, particularly regarding variability within cell types. We examined the prevalence of sex-biased gene transcripts across 23 donors, 12 tissues and 20 broad cell type groups from Tabula Sapiens 2.0 datasets (**Methods, Supp. Fig. 12**). Out of a total of 61,852 gene transcripts evaluated, we identified a diverse set of sex-biased genes varying across different cell types and tissues. Sex-biased genes that are specific to tissue-cell type resolution partially overlap with those identified at the tissue resolution and are almost equal in number in male and female (**Fig. 5A)**. These sex-biased genes showed a skewed pattern of tissue-cell type sharing (**Fig. 5B**), which chromosome X and Y-related genes such as *UTY*, *RPS4Y1*, *XIST*, *LINC00278, EIF1AY* and *DDX3Y* were among the most shared across tissue-cell type pairs, consistent with the pattern found in Genotype-Tissue Expression (GTEx) project at tissue resolution^117^.

**Figure 5:**
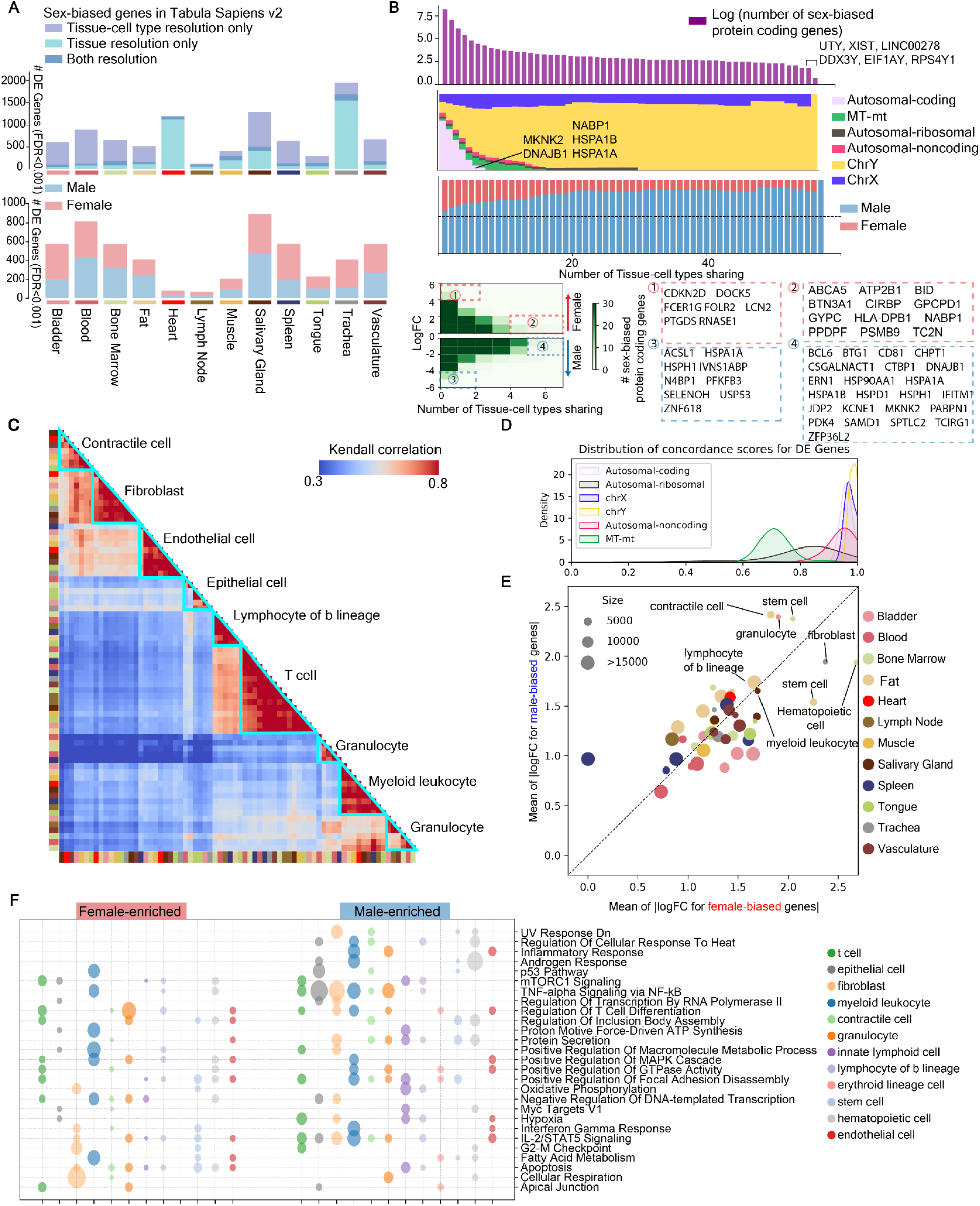
Sex-biased gene expression across tissue-cell types. **A**) Bar graphs displaying the number of sex-biased genes identified in the Tabula Sapiens 2.0, at both the tissue resolution and tissue-cell type resolution. Top: the total number of sex-biased genes across tissue-cell types in Tabula Sapiens 2.0, and overlap with sex-biased genes found at the tissue resolution. Bottom: The proportion of male- and female-enriched genes identified at the tissue-cell type level, and displayed for each tissue. **B**) Top: A bar plot showing the log-transformed number of sex-biased protein-coding genes and the extent to which they are shared across tissue-cell types. The number represents the tissue-cell types sharing that gene, with values greater than the indicated threshold. Middle: Heatmaps showing the distribution of gene types (e.g., autosomal-coding, X/Y-linked) alongside the number of tissue-cell types sharing them. Bottom left: The distribution of log-transformed fold changes (logFC) for male- and female-enriched genes, grouped by tissue-cell type sharing. Bottom right: A detailed gene list for the numbered regions from the heatmap. **C**) Clustering heatmap of Kendall correlations for sex-biased gene expression across major cell types. The inner bar denotes the specific cell type, while the outer bar represents the tissue of origin. The clustering highlights patterns of similarity in sex-biased gene expression between cell types, revealing which tissues and cells share similar sex-specific expression profiles. **D**) Distribution of concordance scores for differentially expressed genes, categorized by autosomal-coding, X/Y chromosome-linked, and mitochondrial genes. **E**) Scatter plot showing the mean logFC for female- and male-enriched genes across cell types. The size of the dots represents the number of sex-biased genes identified in each tissue-cell type. **F**) Bubble plot of representative ontology terms for major cell types. The bubble size corresponds to the adjusted p-value of the enriched terms. The color coding for cell types and tissues is consistent throughout the figure.

Studies on sex differences using bulk tissue GTEx data have revealed small but statistically significant effects of sex on gene expression, driven by sex chromosomes and hormones across various tissues, and suggest that sex correlates with tissue cellular composition^117^. Analysis of sex differences within Tabula Sapiens also reaches similar conclusions at both the pseudobulk tissue and cell type level (**Supp. Fig. 13**). Notably, we observe marked differences in the cell-type specificity of these mechanistic categories. Y-linked genes (e.g. DDX3Y, UTY) exhibit robust and consistent male-biased expression across many cell types, reflecting their strict male specificity and the lack of dosage compensation on the Y chromosome (**Supp. Fig. 13A**). Interestingly, TBL1Y, which has an X-linked homolog (TBL1X), shows variable male-biased expression across cell types. This likely indicated cell-type-specific regulation and partial redundancy with TBL1X, which escapes X-inactivation in some contexts^128^. This identification highlights the complexity of Y-linked gene expression beyond binary sex specificity. Some additional Y-linked genes such as OFD1P12Y, XGY1, and GYG2P1 are annotated as pseudogenes, and showed inconsistent male-biased expression across cell types, probably due to their non-coding nature. X-inactivation escapees (e.g. XIST) show consistent female-biased expression patterns across a wide range of cell types. However, consistent with previous reports^128^, we also observe variability in escape status across cell types, indicating the dynamic nature of XCI escape, and its contribution to both consistent and cell-type-specific sex-biased expression (**Supp. Fig. 13B-C**). Hormone-responsive genes exhibit more cell-type specific sex-biased expression. In particular, we observed that androgen-responsive gene sets are enriched for male-biased expression in specific cell types, including bladder transitional epithelial cells, bone marrow hematopoietic cells, and bladder myeloid leukocytes. In contrast, estrogen-responsive gene sets were enriched for female-biased expression in cell types such as heart fibroblasts, muscle myeloid leukocytes, and others. This suggests that hormone signaling drives localized sex differences, in contrast to the broader, more consistent effects of Y-linked genes and X-inactivation escapees (**Supp. Fig. 13D-E**). We also identified the tissue and cell type specificity of expression of sex-biased transcription factors (**Supp. Fig. 13F**).

To further investigate X chromosome inactivation (XCI), we collected sex-differentially expressed genes at an FDR 1<% across tissues and cell types. We then curated a list of XCI-related genes from past study^128^, which classified X-linked genes into three categories: escape, variable, and inactive with respect to XCI status. By intersecting our significant sex-differential genes with these annotated XCI categories, we observed that genes known to escape XCI are significantly enriched among female-biased genes (**Supp. Fig. 14**). This trend was consistent at both the tissue and cell-type levels. These results support the role of XCI escapees in shaping female-biased expression and align well with previous findings from bulk RNA-seq datasets^117,128^.

The bulk nature of GTEx data poses challenges to identify cell type-specific effects. A significant limitation is that bulk gene expression averages out cell type-specific differences, masking the heterogeneity within tissue samples and resulting in small effect sizes that may obscure important cell type-specific variations^129^. Therefore, we compared the sex-biased genes identified from Tabula Sapiens with those from the GTEx project to identify both congruences and disparities. Tabula Sapiens has a greater number of gene transcripts (61,852 transcripts) compared to GTEx (35,431 transcripts). Our comparison revealed that 38.2% of differentially expressed (DE) genes identified in GTEx were also detected in the Tabula Sapiens dataset across 8 tissues common to both projects, although the overlap varied by tissue (**Supp. Fig. 15A-C**). Notably, 72.3% of chromosome X-linked DE genes identified in GTEx were also found in Tabula Sapiens, along with an additional 254 chromosome X-linked genes **(Supp. Fig. 15D)**. Interestingly, effect sizes for X chromosome genes identified in Tabula Sapiens were generally significantly larger than those reported by GTEx, with the notable exception of XIST (**Supp. Fig. 15D)**. Additionally, our single cell analysis revealed more ubiquitous sex dependent effects than seen in GTEx; for example, *TSIX* was identified in the bulk GTEx data as sex-dependent in only a few tissues, but was sex dependent in nearly all Tabula Sapiens tissues (**Supp. Fig. 15D**). Interestingly, sex dependent gene expression differences are clearly shared across related cell types independent of the tissue of origin. This can be seen by measuring the correlation of differential gene expression across cell types, highlighting the role of cell type in driving gene expression patterns (**Fig. 5C**). In comparing this single cell analysis with the bulk GTEx data, there is agreement in the correlation heatmap of sex-biased gene expression when focusing on tissue-specific differentially expressed (DE) genes, but the tissue-cell-type-specific analysis from Tabula Sapiens revealed a greater overlap of DE genes across tissues. This overlap further indicates that cell type-specific DE genes are shared across tissues, suggesting that sex bias is primarily driven by cell type specificity rather than tissue-level differences (**Supp. Fig. 15E).**

In addition to the recognition of sex-biased genes residing on the X and Y chromosome shared across more than 80 tissue-cell types, we also identified sex-biased protein-coding genes on autosomes. Notably, several genes, including *NABP1, HSPA1B* and *HSPA1A*, emerged as the most prevalent sex-biased genes across tissues such as vasculature, bladder, fat, bone marrow, as well as in immune cells such as myeloid leukocyte and T cells, and stromal cells such as contractile cells and fibroblast (**Supp. Fig. 16A**). Though the effect size for these genes were relatively small (**Fig. 5B**), they play ubiquitous roles in cellular signaling, hormonal regulation, immune response, and cellular stress responses. For example, *HSPA1A* is one of the heat shock proteins (HSPs), identifying as a lower concentration in female vasculature and potentially indicating higher risk in atherosclerotic disease^130^. *CTBP1-AS*, a corepressor of androgen receptor, is generally upregulated in prostate cancer in males^131^, while *BCL6* in male has been associated with maintaining sex-dependent hepatic chromatin acetylation, with a trend of overt fatty liver and glucose intolerance in males^132^. In contrast, genes such as *CIRBP*, *ABCA5* and *HLA-DPB1* were the most prevalent genes enriched in females. Other genes with large effect sizes, such as *LCN2, PTGDS, RNASE1* were also enriched in females (**Fig. 5B)**. HLA genes play a crucial role in immune responses, and recent studies suggest they contribute disproportionately to sex biased vulnerability in immune-mediated diseases, such as autoimmune disorders^133–135^. Additionally, sex differences have been linked to the extent to which HLA molecules influence the selection and expansion of T cells, as characterized by their T cell receptor variable beta chain^136^. Adipose LCN2 is frequently enriched in females, where it acts in an autocrine and paracrine manner to promote metabolic disturbances, inflammation, and fibrosis in female adipose tissue^137^. Conversely, several genes, including *SELENOH*, *USP53*, and *ZNF618*, were upregulated in males but not females, showing high effect sizes and abundance. This distinct expression pattern between sexes suggests that males and females may employ different molecular pathways to regulate key biological processes. The identification of these shared and significantly affected genes underscores their potential impact on sex-specific traits and disease susceptibilities. Furthermore, a density plot of concordance score across different gene types showed that chrY and chrX genes exhibit the most consistent sex effects, while mitochondrial genes demonstrated the least concordance (**Fig. 5D, Supp. Fig. 16B**).

Given the cell type specificity of sex differential expression, we further investigated which cell types exhibited the greatest variability in sex-based gene expression. Cell types with the largest number of sex-biased genes include myeloid leukocyte, innate lymphoid cell, epithelial cells, and granulocytes in tissues such as bone marrow, vasculature, and blood (**Supp. Fig. 16C-D**). Scatter plots of effect sizes in female and male donors indicated the highest mean effect size in fat, bone marrow tissues, particularly in stem cells, fibroblast and immune cells, such as lymphocytes and myeloid leukocyte (**Fig. 5E**). These findings align with previous studies that emphasize the importance of cell-type-specific analyses in uncovering sex differences in gene expression. For example, fibroblasts, which show the highest mean effect size, are known for their crucial roles in tissue homeostasis, processes that can be differentially regulated by sex hormones^138,139^. Immune cells such as lymphocytes, especially in the fat and bone marrow, reflect known sex differences in immune responses^140^. To link bulk and single-cell findings, we examined autosomal genes from GTEx that were most strongly co-expressed with X-linked genes and compared them to top-ranking tissue-cell types identified in Tabula Sapiens. These GTEx-derived autosomal gene modules were highly correlated with the TS-inferred sex-biased cell types (**Supp. Fig. 17**).

We summarized the representative functional enrichment for each cell type between male and female donors (**Fig. 5F**). In general, pathways are shared by both sexes but can be upregulated in a sex-dependent fashion in a highly cell type specific manner. In females, pathways related to cellular respiration and fatty acid metabolism are enriched in female fibroblast, while immune activity is enriched in granulocyte and T cells. In males, pathways associated with inflammatory response were predominantly enriched in endothelial cells and myeloid leukocytes, while androgen responses were predominantly enriched in epithelial, myeloid leukocyte and hematopoietic cells. These findings suggest that males and females leverage different molecular pathways to regulate key biological processes, which may contribute to sex-specific disease susceptibilities and physiological functions.

### Searchable Extended Metadata

The Tabula Sapiens 2.0 also includes de-identified medical records for all donors. These medical records are voluminous (an average of 13 pages per donor, 291 total pages) and the overhead associated with their interpretation is often prohibitive. Therefore, we developed ChatTS (https://singlecellgpt.com/chatTSP?password=chatTS; source code available at https://github.com/Harper-Hua/ChatTS)—a web application powered by a large language model—to facilitate use of Tabula Sapiens for the broader research community.

On the front end, a user asks a question about donors’ medical records. Then, on the back end, an external LLM is prompted to answer this question on a per-donor basis, using free-form text that was manually extracted from each donor’s medical record (including physical, imaging, and laboratory studies, as well as social and medical histories). Results are concatenated and returned as a table which can be downloaded and manually verified (**Fig. 6A**). Interestingly, we observed that the underlying LLM can interpret charts and offer conclusions that are not explicitly stated in the records (**Fig. 6B**). For example, when asked which donors have heart disease, the response did not require the explicit words “heart disease” in the medical records, but rather inferred it from symptoms and treatments.

**Figure 6.**
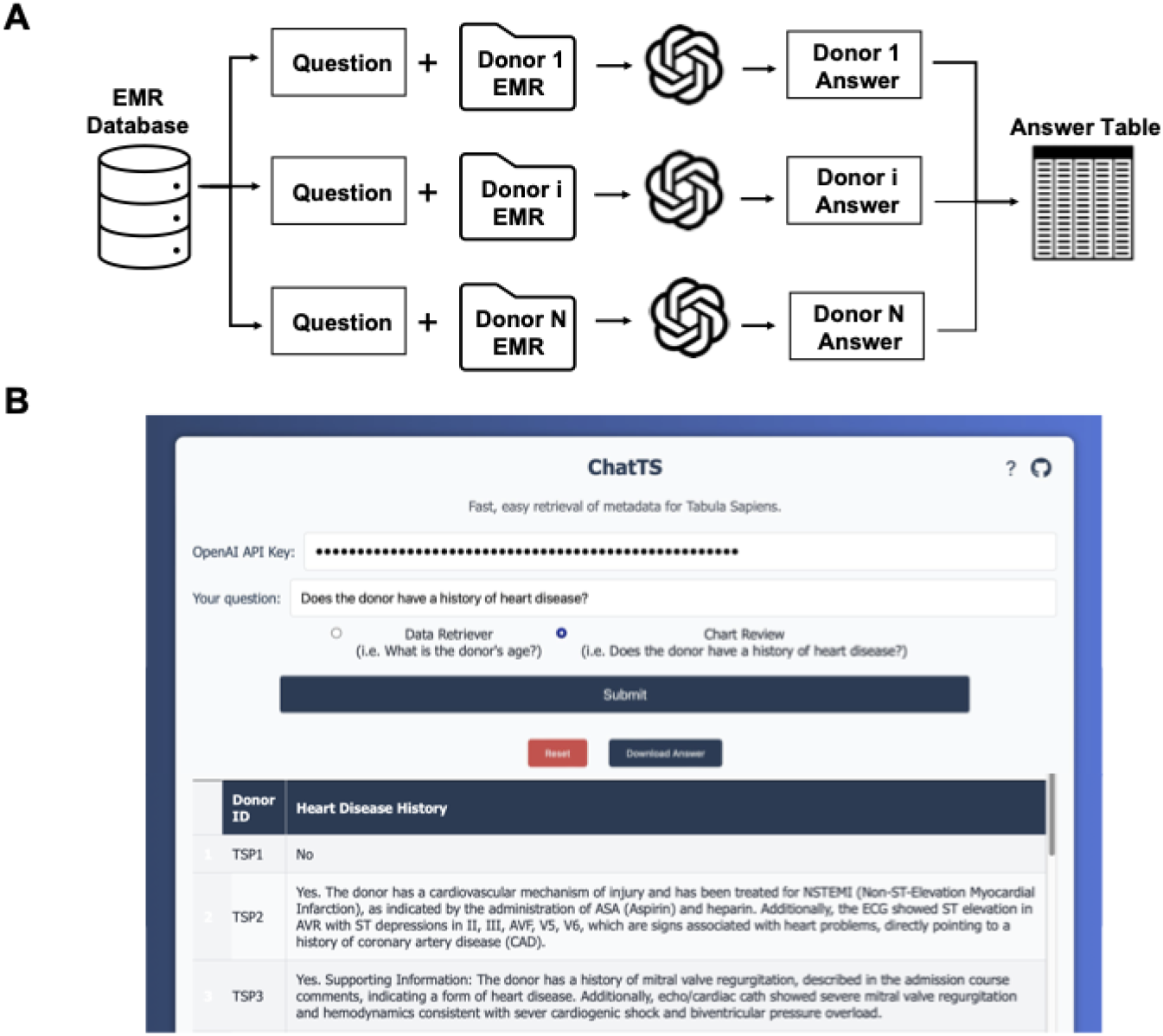
ChatTS web application overview. **A)** Schematic representation of the ChatTS web application architecture. **B)** Screenshot of the application, demonstrating inference based on information in charts.

We benchmarked ChatTS with a GPT-4o backend against 330 manually curated question-answer pairs regarding donor metadata. ChatTS showed satisfactory performance (83.2% agreement with manually curated data), and in many cases, the ChatTS-generated answers were more informative than the manually curated data, **Supp. Table 7**. Differences between manual and ChatTS answers tended to be related to ambiguities in the questions, their definitions, and information present in the medical records, and not to hallucination.

### Limitations of the Study

RNA abundance, while informative, does not fully capture protein levels or post-translational modifications, limiting to some extent inference on functional protein states and short time scale behavior of the cells. Cell type abundances measured by dissociation do not necessarily represent true biological abundance due to differential dissociation effects and because enrichment was performed by compartment (immune, epithelial, endothelial and stromal) to maximize the number of cell types captured. Conversely, very rare cell populations may be missing or due to the requirements of live single-cell RNA-seq, as samples were required to be processed quickly and with consistent methods overnight for each donor. As might be expected of any adult who has lived a full life, none of our donors were perfectly healthy. All donors are missing some tissues due to organ transplantation, which always had a higher priority than our study.

## Conclusions

The Tabula Sapiens 2.0, featuring over 1.1 million cells across 28 tissues from healthy donors aged 22 to 74 years, offers a comprehensive multiorgan dataset enabling systematic analysis of rare cell populations, while accounting for technical artifacts and donor-specific variations. Global analysis of transcription factor expression enabled a comprehensive analysis to identify which transcription factors are cell type specific and which are ubiquitously expressed. Collectively, we observe heterogeneous and cell type-specific senescence phenotypes and identify several novel and sometimes contradictory fates for cells that undergo cell cycle arrest. Understanding the diverse senescence phenotypes across different cell types offers promising avenues for developing targeted senolytics, tailored to address age-related diseases affecting specific tissues and cell types. Finally, sex-dependent gene expression analyses provide the beginnings of an understanding which genes and cell types are most strongly sexually dimorphic and the consequent implications for cellular and tissue functions. By comparing male and female gene expression profiles, we identified significant patterns and quantified the impact of sex on gene regulation, contributing to our understanding of biological differences and their consequences for health and disease.

Tabula Sapiens 2.0 is the most comprehensive human cell atlas reference to date, as measured by the breadth of tissues and cell types. The vignettes included in this publication illustrate the kind of exploratory analysis and hypothesis building that such a dataset enables. Following on the previous Tabula projects^1,21,141,142^, all data is available for the community to continue learning and improving as a reference atlas until we have a deeper understanding of the molecular representation of cell types and cell states. The community found many uses of Tabula Sapiens 1.0, ranging from being a tool to predict potential on target toxicity of drug candidates to being a reference standard used to evaluate bioinformatic tools and train AI models, and we expect that the expanded version described here will only increase such applications. For example, the discovery of cell type specific transcription factor usage will point the way for future biochemical studies to characterize the function of these transcription factors, as will the identification of previously undetected ubiquitous transcription factors which are found in all cell types. We have also already seen the use of Tabula Sapiens 2.0 as a hold out data set to test the zero shot capability of AI models which are trained on Tabula Sapiens 1.0.

## Resource availability

### Lead contact

Further information and requests about reagents and other resources used in this work should be directed to and will be fulfilled by the lead contact, Stephen R Quake.

### Data and code availability

● The entire dataset can be explored interactively at the Tabula Sapiens Data Portal^143^ (https://tabula-sapiens-portal.ds.czbiohub.org/).
● The code used for the analysis is available on the github repository for Tabula Sapiens (https://github.com/czbiohub-sf/tabula-sapiens/).
● Gene counts and metadata are publicly available from figshare^144^ and cellxgene^143^.
● The raw data files are available from a public AWS S3 bucket (https://registry.opendata.aws/tabula-sapiens/), and instructions on how to access the data have been provided in the project GitHub.
● To preserve the donors’ genetic privacy, we require a data transfer agreement to receive the raw sequence reads. The data transfer agreement is available in the data portal.
● Any additional information required to reanalyze the data reported in this paper is available from the lead contact upon request.

## Acknowledgements

This project has been made possible in part by grant nos. 2019-203354, 2020-224249, 2021-237288, 2021-006486, 2022-316725. from the Chan Zuckerberg Initiative DAF; an advised fund of Silicon Valley Community Foundation; and by support from the Chan Zuckerberg Biohub San Francisco.

We thank Donor Network West for procuring the organs and tissues from the donors for this project. We are also grateful to the UCSF Liver Center (funded by NIH P30DK026743) for assistance with the liver cell isolations, and B. Tojo for the original artwork in Fig. 1. We would like to express our gratitude and thanks to donor WEM and his family at Donor Network West, as well as to all the anonymous organ and tissue donors and their families for giving both the gift of life and the gift of knowledge through their generous donations. We would also like to thank Dr. Konstantin Kahnert in Dr. Emma Lundberg’s lab for extracting subcellular location information from the Human Protein Atlas website.

## STAR methods

### Key resources table

[will be moved here once manuscript is ready, for now please check the file STARmethods_KeyResourcesTable_TabulaSapiensv2.docx]

### Organ and tissue procurement

To maintain consistency in the overall Tabula Sapiens dataset, organ and tissue procurement followed the same procedure used in the first phase of the project. Donated organs and tissues were procured at various hospital locations in the Northern California Region through collaboration with a not-for-profit organization, Donor Network West (DNW, SanRamon, CA, USA). DNW is a federally mandated organ procurement organization (OPO) for Northern California. Recovery of non-transplantable organ and tissue was considered for research studies only after obtaining records of first-person authorization (i.e., donor’s consent during his/her DMV registrations) and/or consent from the family members of the donor. However, the pancreas from donor TSP9 was provided by Stanford University Hospital under appropriate regulatory procedures. The research protocol was approved by the DNW’s internal ethics committee (Research project STAN-19-104) and the medical advisory board, as well as by the Institutional Review Board at Stanford University which determined that this project does not meet the definition of human subject research as defined in federal regulations 45 CFR46.102 or 21 CFR 50.3.

Tissues were processed consistently across all donors. Each tissue was collected, and transported on ice, as quickly as possible to preserve cell viability. A private courier service was used to keep the time between organ procurement and initial tissue preparation to less than one hour. Single cell suspensions from each organ were prepared in tissue expert laboratories at Stanford and UCSF. For some tissues the dissociated cells were purified into compartment-level batches (immune, stromal, epithelial and endothelial) and then recombined into balanced cell suspensions in order to enhance sensitivity for rare cell types as described further below.

### Tissue preparation protocols and cell suspension preparation protocols

The methods for tissue dissociation and staining of cell suspensions for bladder, blood, bone marrow, eye, fat, heart, intestine, kidney, liver, lung, lymph node, mammary gland, pancreas, prostate, salivary gland, skeletal muscle, skin, spleen, tongue, trachea, thymus, uterus, and vasculature are described in the methods section of the Tabula Sapiens^1^.

### Inner Ear - Initial tissue preparation protocol

Whole organ utricles were collected from organ donors. Bilateral utricles were harvested from organ donors as previously described^145–147^. Briefly, a post-auricular incision was made followed by a modified transcanal approach to expose the middle ear. The tympanic membrane, malleus and incus were removed while keeping the stapes in situ. To expose the vestibular organs, the bony covering of the vestibule was thinned using a diamond burr on low speed, the stapes footplate removed, and the oval window widened. The utricle was harvested from the elliptical recess and placed in PBS on ice for single-cell RNA sequencing analysis.

### Inner Ear – 10x Genomics sample preparation

Utricles from organ donors were placed in DMEM/F12 (Thermo Fisher Scientific/Gibco, 11-039-021) with 5% FBS during transport to the laboratory. Tissues were then washed twice in DMEM/F12 and any debris and bone microdissected away. The whole utricle was then incubated with thermolysin (0.5 mg/mL; Sigma Aldrich, T7902) for 45 min at 37°C. Next, the whole utricles were digested using Accutase (Thermo Fisher Scientific, 00-4555-56) for 50 min at 37°C and single cell suspension was obtained by trituration using a 1 mL pipette. Single cell suspension was achieved using a 40 μm filter. DMEM/F12 media with 5% FBS was used for this step. Cells were then centrifuged at 300g for 5 min at 4°C. The supernatant was removed, and cells were resuspended in media. Number of cells per microliter was quantified using a hemocytometer

### Large and Small Intestines – Initial Tissue Preparation Protocol

Segments of the small intestine (duodenum and ileum) and large intestine (ascending and sigmoid colon) were transported on ice to Stanford University. Upon arrival, the tissues were sectioned into approximately 5 cm segments and rinsed multiple times with 35 mL of ice-cold PBS (Thermo Fisher Scientific 10010023) to remove residual lumen contents. The tissues were then opened with the mucosal side facing up, and 10 mL of ice-cold PBS was injected using a 21G needle. Mucosal blebs were then dissected, minced into approximately 2 mm pieces, and digested in 10 mL of digestion medium (DMEM/F12: Thermo Fisher Scientific 12634010, 1x Penicillin-Streptomycin-Neomycin: Sigma P4083, 1x Normocin: Invivogen ant-nr-2, 1x HEPES buffer: Corning 25-060-CI, 1x Glutamax: Thermo Fisher Scientific 35050061, and 10 µM ROCK inhibitor Y-27632: MedChem Express HY-10583) containing Collagenase III (300 U/mL: Worthington Biochemical Corporation LS004182) and DNase I (200 U/mL: Worthington Biochemical Corporation LS002139) at 37°C for 90 minutes. During digestion, the mixture was pipetted every 15 minutes with a 10 mL pipette to facilitate tissue breakdown. After digestion, 30 mL of intestinal wash buffer (HBSS: Corning 21-022-CV, 2% FBS, 1x Penicillin-Streptomycin-Neomycin, 1x Normocin, and 10 µM ROCK inhibitor Y-27632) was added to the suspension, which was then centrifuged at 1500 rpm for 5 minutes at 4°C. The supernatant was discarded, and the pellet was resuspended in 5 mL of ACK lysis buffer (Thermo Fisher Scientific A1049201) to lyse red blood cells and incubated for 5 minutes at room temperature. The suspension was centrifuged again, and the pellet was resuspended in intestinal wash buffer with DNase I, then filtered through a 100 µm cell strainer (Miltenyi Biotec 130-098-463). The final cell suspension was counted using trypan blue (Thermo Fisher Scientific 15250061) and adjusted to a concentration of 10^6^ cells/mL for direct single-cell analysis using the 10x Genomics platform or flow sorting.

### Ovary - Initial tissue preparation protocol

Ovaries were finely dissected to remove surrounding tissues. Ovaries were gently chopped using a razor blade and then transferred to a 50 mL Falcon^TM^ conical tube with a final volume of 30 mL 2 mg/mL collagenase type IV in DMEM/F12. Tubes were incubated at 37 °C shaking at 250 rpm for 20 min then centrifuged at 250 x g for 5 min. The supernatant was aspirated and 30 mL 0.25 % trypsin-EDTA was added. Tubes were incubated at 37 °C shaking at 250 rpm for 20 min then centrifuged at 250 x g for 5 min. The supernatant was aspirated, and the tissue was quenched with 30 mL 10 % FBS in DMEM/F12. Using a P1000, tissue was gently triturated to generate a single cell suspension. The cell suspension was filtered using a reversible 37 um sieve. The sieve was rinsed with 1 mL quench buffer (<37 um filtered flow through) and then flipped and placed in a new tube for rinsing with 1 mL quench buffer (>37 um unfiltered mature oocyte pool).

Single oocytes and/or follicles were handpicked from the cell suspension under magnification (2 to 6X) on a dissection microscope using an EZ-Grip pipette with a 125 um EZ-Tip and washed in 0.04% BSA-PBS. Single oocytes and/or follicles were then transferred in 0.6uL of 0.04% BSA-PBS to individual wells in a 96-well plate containing a lysis buffer master mix. Lysed oocytes and/or follicles were stored at -80 °C to enhance lysis prior to cDNA and sequencing library construction. Nine follicles, which were not dissociated into single-cell suspensions, represent the bulk transcriptomes of entire ovarian follicles.

### Ovary - FACS/SS2 sample preparation

Filtered cell suspensions were centrifuged at 250 x g for 5 min, aspirated, and washed in FACS Buffer (2 % FBS-PBS). The following antibodies were used at a 1:200 dilution in FACS Buffer: EPCAM-PE (Biolegend, 324206); CD45-FITC (Biolegend, 304038); CD31-APC (Biolegend, 303116). The cell suspensions were incubated 1 hr on ice for staining, then washed 3 times. Prior to sorting, Sytox Blue was added at a 1:1000 dilution (ThermoFisher, S34857) and live, single cells were sorted for EPCAM+ (oocytes), CD45+ (endothelial), CD45+ (immune), EPCAM-/CD45-/CD31- (stromal/granulosa) populations into 384-well lysis buffer plates following color compensation. Plates were stored at -80 °C until processing for sequencing.

### Stomach – Initial Tissue Preparation Protocol

Ligated whole stomachs were transported on ice to Stanford University. Upon arrival, a 6x3cm longitudinal strip was cut from the anterior aspect of the stomach antrum, 6 cm proximal to the pylorus. Stomach tissue was first rinsed in 1X PBS to remove clots and debris, and blotted dry to remove excess mucus. The tissue was then dissected, separating the mucosa/submucosa layer from the muscularis/serosa layer, and the layers were then weighed. Each layer was digested separately to maximize viability. The mucosa was first incubated with 5mM EDTA in HBSS buffer without Ca^2+^ for 10 minutes on a shaker at 37 °C (x3), with washes and collection of epithelial cells. After this, both mucosa and muscularis layers were minced with fine sharp scissors in digestion media with 0.8 mg/ml collagenase Type IV Worthington (Sigma) and 0.05 mg/mL DNase I (Roche) (10mL per 2.5 gram of tissue). Two serial mucosa digestion were done in an orbital shaker (300 rpm) at 37 °C, 20 min each, and for muscularis 40 min each (vortexing halfway). Digestions were stopped with cold R10-EDTA, and cells were filtered from undigested tissue, washed and then stained with CD45-PE (Biolegend, #304007) for cell counting and sorting. The Mucosa digestion and EDTA epithelial fraction were combined and DAPI stained (Thermofisher Cat# D1306). Live DAPI-, CD45+ and CD45- cells were then FAC-sorted on a BD Aria. Sorted cells from mucosa and muscularis were mixed back together at a 70:30 CD45+ to CD45- ratio, and used for 10X single-cell RNA sequencing.

### Testis- Initial Tissue preparation protocol

Testis tissues were dissected out from the tunica and lightly dissociated and minced using sanitized surgical forceps and razor blade. Two to three pieces of 0.5g testis tissue was placed in a 50mL Falcon tube with 10mL prewarmed collagenase solution at 32 °C (1 mg/mL collagenase I (Worthington Biochemicals #LS004196), 1 mM EDTA, 0.5ul/mL DNaseI (50mg/mL stock, Worthington Biochemicals #LS002139) in PBS). Testis tissue was first dissociated by vigorous pipetting and then incubated at 32 °C for 8 minutes with 2-3 times pipetting during the incubation to resuspend cells. After incubation, cells were dissociated again at room temperature by vigorous pipetting for 2-4 minutes. Cells were spinned down at 250xg for 5min at room temperature and then the supernatant was removed. The dissociation steps were repeated one more time before 5mL of TripLE (Thermo Fisher Scientific #12604013) plus 1uL/mL DNaseI (50mg/mL stock) was added to the dissociated cells and incubated at 32 °C for 15 min with pipetting every 5 min. Cells were then washed with 30mL PBS and filtered through a 70 µm cell strainer and then a 40 µm cell strainer. Cell number was recorded before cells were spin down at 250xg for 10min. Then the supernatant was removed, and cells were resuspended at 1000 cells/uL for cell capture in microfluidic droplets with the 10x Genomics platform.

### Testis – FACS/SS2 sample preparation

After cells were spun down at 250xg for 10min and after passing the cell strainer, cells were washed once in 3mL FACS buffer (2%FBS (Thermofisher #A5670701), 1mM EDTA (Invitrogen #AM9260G) in PBS(Gibco #10010023)), spin down at 25xg for 5min, and resuspended in 400uL of FACS buffer. Antibodies cocktail (2 µL of EPCAM-PE (Biolegend, 324206), 2uL of CD45-FITC (Biolegend, 304038), 2uL of CD31-APC (Biolegend, 303116)) was added to the cell suspension, and cells were incubated on ice for 30min, with mix by tapping every 10min. After incubation, cells were washed twice with 2mL FACS buffer and spin down at 250xg for 5min, and resuspended in 1mL FACS buffer. Sytox blue was added before sorting.

### Thymus– Initial Tissue Preparation Protocol

The tissue dissociation and staining of cell suspensions for thymus is described in the methods section of the Tabula Sapiens^1^. Our analysis revealed the presence of cells originating from fat and intra-thymic lymph nodes among those attributed to the thymus, likely due to difficulties in tissue collection caused by the natural involution of the thymus with age. Consequently, we excluded the thymus from our analysis of sex-biased gene expression.

### 10x Genomics protocol

10x Genomics kits used were Chromium Next GEM Single Cell 3′ Kit v3.1 or Chromium Next GEM Single Cell 5ʹ Kit v2. The protocols provided by the manufacturer were followed. In general, two 10X Chromium channels were loaded per tissue with 7000 cells, with the goal of obtaining data for 10,000 viable cells per tissue.

### Organ and cell coverage

Our goals were to characterize the gene expression profile of 10,000 cells from each organ and detect as many cell types as possible. As explained in detail for each organ, about ⅔ of the organs employed a MACS based enrichment strategy, either to balance cell types between four compartments; epithelial, endothelial, immune, and stromal. This ensured abundant cell types in one compartment did not mask rare cell types in another. Two 10x reactions per organ were loaded with 7,000 cells each with the goal to yield 10,000 QC-passed cells. Four 384-well Smartseq2 plates were run per organ. In most organs, one plate was used for each compartment (epithelial, endothelial, immune, and stromal), however, to capture rare cells, some organ experts allocated cells across the four plates differently. The use of two 10x reactions enabled some flexibility to distinguish in the data the anatomical position of the sample or allowed enrichments other than epithelial, endothelial, immune, and stromal. It also served as insurance against losing an entire organ due to a clog of the 10X chip.

### Flow Sorting

Details of the sorting can be found in Tabula Sapiens^1^. Briefly, after dissociation, the single cells from each organ and tissue that were destined for plates were isolated into 384 plates via FACS. On some cell suspensions destined for 10x tissue compartments were enriched by FACS to balance the cell types as described in the tissue specific methods. Most sorting was done with SH800S (Sony) sorters. The last column of each 384 well plate was intentionally unsorted so that ERCC’s controls could be used as a plate processing control. Immediately after sorting, plates were sealed with a pre-labelled aluminum seal, centrifuged, and flash frozen on dry ice to ensure full cell lysis. Typical sort times were 4 to 9 minutes per 384-well plate.

### Smart-seq2 Protocol

Plate-based sequencing of tissues used the 384 plate modification of Smart-seq2^148^ as described in Tabula Sapiens^1^ and Tabula Muris Senis^142^.

### Smart-seq2.5 Protocol

Ovary plates in the object denoted “SS3” were processed into cDNA by a modified protocol called “Smart-seq2.5,” which is a revision of the Smart-seq2 protocol that takes advantage of the improved reagents and reaction conditions used in Smart-seq3xpress^149^. Briefly, lysis plates were prepared by dispensing 0.5 µL lysis buffer (0.15% Triton X100 (Sigma-Aldrich, 93443-100ML), 10% polyethylene glycol 8000 (Sigma-Aldrich, P1458-50ML), 0.6 u/µL Recombinant RNase Inhibitor (Takara Bio, 2313B), 1 mM of each dNTP (Roche, NTMIXKB), 0.25 µM biotinylated oligo-dT30VN (Integrated DNA Technologies, 5’-biotin-AAGCAGTGGTATCAACGCAGAGTACT30VN-3′), and 1:600,000 ERCC spike-in RNA (Thermo Fisher Scientific, 4456740)) into 384-well hard-shell PCR plates (Bio-Rad HSP3901) using a Mantis liquid handler (Formulatrix). The lysis plates were then sealed with AlumaSeal CS films (Sigma-Aldrich, Z722634), spun down, snap-frozen on dry ice, and stored at -80 °C until sorting. Dissociated cells were sorted into thawed lysis plates by FACS as described above, after which the plates were spun down, snap-frozen on dry ice, and stored at -80 °C until further processing.

For cDNA synthesis, plates were first incubated at 72 °C for 10 min to ensure cell lysis and RNA denaturation. Then 0.5 µL RT mix (50 mM Tris-HCl pH 8.0 (Thermo Fisher Scientific, BP1758-100), 60 mM NaCl (Invitrogen, AM9760G), 5 mM MgCl_2_ (Invitrogen, AM9530G), 2 mM GTP (Thermo Scientific, R1461), 16 mM mM DTT (Promega, P117A), 1.5 mM TSO (Integrated DNA Technologies, 5’-AAGCAGTGGTATCAACGCAGAGTGAATrGrGrG-3’), 0.5 u/µL Recombinant RNase Inhibitor (Takara Bio, 2313B), and 4 u/µL Maxima H Minus Reverse Transcriptase (Thermo Scientific Scientific, EP0751)) was dispensed into each well using a Mantis liquid handler. Reverse transcription was then carried out on a Bio-Rad C100 x384 thermal cycler using the following program: 1) 42 °C for 90 minutes, 2) 10 cycles of 50 °C for 2 min and 42 °C for 2 min, and 3) 85 °C for 5 min. The resulting cDNA was amplified by dispensing 1.5 µL pre-amplification mix (1.67x SeqAmp PCR buffer (Takara Bio, 638526), 0.083 µM ISPCR primer (Integrated DNA Technologies, 5’-AAGCAGTGGTATCAACGCAGAGT-3’), and 0.042 u/µL SeqAmp DNA polymerase (Takara Bio, 638504); concentrations were chosen to reach 1x SeqAmp PCR buffer, 0.5 µM ISPCR primer, and 0.025 u/µL SeqAmp DNA polymerase in the 2.5 µL final reaction) using a Mantis liquid handler and then performing PCR on a BioRad C1000 384-well thermal cycler (Bio-Rad) with the following program: 1) 95 °C for 1 min, 2) 13-19 cycles of 98 °C for 10s, then 65 °C for 30 s, and 68 °C for 4 min, and 3) 72 °C for 10 min. The resulting cDNA was diluted 1:5 by adding 10 uL of buffer EB (Qiagen Cat#19086) using a Mantis liquid handler. Sequencing libraries were prepared and pooled from the diluted cDNA plates using the same protocol as for Smart-seq2.

### Sequencing

All Sequencing was done by the CZBiohub San Francisco Sequencing Team. 10X libraries were loaded on Illumina NovaSeq 6000 S4 flow cells in sets of 16 libraries with the goal of generating 50,000 to 75,000 reads per cell. SmartSeq libraries were run in sets of 20 plate libraries with a target of 1M reads per cell.

### Biobanking and Histology

Where possible, additional tissue samples were collected from the vicinity of sequenced specimens for frozen and fixed biobanking. Frozen samples were washed and flash frozen with liquid nitrogen. Fixed samples were fixed in 10% buffered formalin and paraffin embedded (FFPE). Hematoxylin and eosin (H&E) stained slides were generated from the FFPE samples using standard methods, digitally scanned using Leica Aperio AT2 23AT2100. Images are available on the Tabula Sapiens portal.

### scRNAseq data extraction

Sequences from the NovaSeq 6000 were de-multiplexed using bcl2fastq version 2.20.0.4.22. Reads were aligned to the Gencode Reference version 41 (GRCh38) genome using STAR^150^ version 2.7.11b with parameters TK. Gene counts were produced using HTSEQ^151^ version 2.0.5 with default parameters, except ‘stranded’ was set to ‘false’, and ‘mode’ was set to ‘intersection-nonempty’. Sequences from the microfluidic droplet platform were de-multiplexed and aligned using CellRanger version 7.0.1, available from 10x Genomics with default parameters.

### scRNAseq data pre-processing and cell type annotations

Gene count tables were combined with the metadata variables using the Scanpy^152^ Python package version 1.9.6. We first roughly filtered the dataset to remove any cells with fewer than 100 genes and 1000 counts (unique molecular identifiers or UMIs). In order to filter out reads from ambient RNA we ran DecontX^153^ (implemented in R package celda v1.16.1) separately for each 10X run, using default parameters given the full background and the filtered cells. After the DecontX filtering step, we re-filtered the dataset more strictly using a minimum threshold of 200 non-mitochondrial genes and 2500 non-mitochondrial counts for the droplet (10X) cells and 500 non-mitochondrial genes and 5000 non-mitochondrial counts for the smartseq FACS sorted cells. This filtered gene-count matrix was then used for the analysis.

Ambient RNA and barcode swapping^154^ are known problems in 10x sequencing. After using DecontX to remove ambient RNA, we removed all cells sharing both the cell and transcript barcode but not the same sample barcode in each sequencing run in order to remove cells generated by barcode swapping.

In the analysis step, we first integrated the multiple batches of data from each donor to generate a unified visualization of the cells using scVI^155^ from scvi-tools^156^ release 0.20.0. For training the variational autoencoder neural network, we used the following hyper parameters: n_latent=50, n_layers=5, dropout_rate=0.1. We allowed each gene to have its own variance parameter by setting dispersion="gene". We trained the scVI model for 50 iterations with all available data and corrected the batch effect associated with donor and technology. scVI generated a harmonized latent space that was then projected to a 2D space using UMAP. This process was done individually for each organ, and we then shared the harmonized data along with the reduced dimensional latent space in a h5ad format data object compatible with both Scanpy and CELLxGENE. CELLxGENE is a data exploration and visualization tool that allows users to interactively explore any scRNAseq dataset^9,10^. Manual annotation was performed by tissue experts using CELLxGENE. Each data object contained three main components: gene count data, cell-wise metadata, and gene-wise metadata for their organ of interest. CELLxGENE allows the user to color cells by any cell metadata such as donor and compartment. Cells can also be colored by gene expression data. The user can also select cells based on any meta data features, or using a lasso tool. Following each organ and/or tissue manual annotation procedure, a data object containing the new annotations was generated using the same scVI parameters and the annotations were regularized to follow the cell ontology^157^. We did not correct for batch effects associated with organ even though each organ is sequenced separately because of concerns of removing biological variation by over-correction. Cell types missing in the current public version of the cell ontology were added to the provisional Tabula Sapiens cell ontology.

Since Tabula Sapiens was annotated by a large number of experts, quality control (QC) was performed on the manual annotations by using the automatic annotation tool popV^158^. PopV was applied to all organs in Tabula Sapiens donors 1 and 2 and predictability scores were generated for all cells by running a 5-fold cross validation. For donors 3 to 15 a draft automated annotation was generated using PopularVote. This was followed by manual inspection and annotation of all tissues in this set. This process was iterated in this manuscript by adding manual expert annotations for most organs from donors 17-25 before running popV on all the remaining unannotated cells and donors and cleaning up the labels manually as described above.

### Quality control for contamination correction

To address potential barcode mixtures arising from sequencing experiments where different samples from each donor were loaded in the same 10X Genomics strips, we implemented a contamination correction filter. Specifically, we checked for overlapping barcodes across samples within the same donor. Cells sharing the same barcode identified in multiple samples from the same donor were considered potential contaminants and were excluded from further analysis. This process reduced the total number of cells from 1,150,192 to 1,105,354, enhancing the specificity of downstream analyses.

### Transcription factor activity analysis

Transcription factor activity within each cell type was analyzed by running pySCENIC^159,160^ (v0.12.1) on gene expression data from each cell type separately. The analysis was based on hg38-refseq_r80-mc_v10_clust-gene_based databases downloaded from resources.aertslab.org. Outputs from the ctx step of pySCENIC were used to infer downstream targets of transcription factors while outputs of the aucell step were used for transcription factor activity.

### Transcription factor analysis across cell types using the ***τ*** statistic

τ analysis was done using only the droplet data for consistency across organs. This included 175 cell types of the 180 total cell types identified. The mean of the log normalized expression of each TF was computed for each cell type. Mean expression ranged from 0.0 to 4.9. The τ statistic (2,3) was used to generate a cell type specificity value for each of the 1635 transcription factors from these mean expression values.

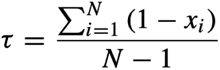

Every TF has a vector, x, of its mean expression in each cell type of the 175(N) cell types. TF vectors were normalized by their maximum expression value and then inverted by subtracting them from 1 to create a vector for each TF of values ranging from 0 to 1 with low values in cell types where it is highly expressed and values near 1 where it is lowly expressed. This vector is summed to make a single value for each TF. These sums are normalized to make τ range from 0 to 1. The Tspex^161^ package, downloaded from https://apcamargo.github.io/tspex/, version-0.6.3 was used to perform this by calling “tspex.TissueSpecificity(TF_exp, ’τ’, log = False)”. The resulting τ values range between 0.4 and 1.0 where low τ values are ubiquitous transcription factors and high τ values are cell type specific. Heatmaps were made using the seaborn clustermap function in seaborn package (v-0.12.2). In the large heatmaps with all cell types, “vmax” was set to the 99.5 percentile, clipping the max value near 1.0 in order to improve visualization. Based on the distribution of τ values in the mouse atlas and a similar cutoff used in the fly cell atlas, we chose a τ threshold of 0.85 as the demarcation between specific (860 TF’s) and non-specific (745 TF’s). There was no clear transition in the τ distribution to distinguish ubiquitous from non-specific TF’s. We attempted to find one by incorporating a lower bound for the mean expression and/or for the fraction of cells expressing the TF, but a principled cutoff criteria did not become clear. We then looked for protein expression of 100 lowest τ TF’s in the HPA to get a different window on ubiquity.

### Transcription factor Gene Ontology Analysis

Curated TF gene names were obtained from the “Collection of known and likely humanTFs (1639 proteins)” data set, downloaded from the University of Toronto transcription factor database. (https://humantfs.ccbr.utoronto.ca/allTFs.php.) 1639 of these gene names were found in our GTF file of which 1635 TF’s showed expression in our dataset. SHOX and ZBED1 had zero expression. The 745 non-specific (τ < 0.85) transcription factors were analyzed for gene set enrichment using the GSEApy package^162^ (v-0.10.5) with GO_Biological_Process_2021 geneset against a background of all 1635 transcription factors. 69 GO terms had an adjusted P-val < 0.02. These were organized manually into broad cellular functions and related terms were placed into subgroups.

### Analysis of the senescent cells phenotypes

Single-cell gene expression data from Tabula Sapiens 2.0 were used to identify senescent cells. Tissues from the sex with only a single donor were excluded from the analysis. After removing 52,149 cells from six tissues, the final dataset included 1.08 million cells from 25 tissues, collected from 21 different donors. Senescent cells were defined as CDKN2A+ MKI67- cells in the filtered dataset. A total of 48,114 senescent cells (∼4.4% of all cells) were identified, spanning 145 cell types grouped into 34 broad categories. These cells were analyzed across different donors, tissues, and cell types. To identify senescence-associated genes (SAGs), differential gene expression analysis (DGEA) was performed by comparing senescent cells to non-senescent cells from the same cell type, donor, and tissue. To detect robust and statistically significant SAGs, we performed differential gene expression analysis (wilcoxon double-sided test) using the Scanpy package (v-1.10.1) with the following thresholds: log fold-change > 0.5, adjusted p-value < 0.001, and minimum percent of non-zero expression in senescent cell group = 0.5. SAGs from different donor-tissue-cell types were then grouped to select SAGs enriched in at least 50% of the donors across tissue-cell types. These enriched SAGs were combined into a final list of 3792 genes, which were enriched in at least 50% donors for at least one tissue and one broad cell type. To visualize the cell type prevalence of SAGs or senescence associated hallmark genes, we reperformed differential gene expression analysis at cell type-level for each of the 3,792 SAGs by comparing all senescent and non-senescent cells across broad cell types and calculated log_2_ fold changes for genes with statistically significant differences (adjusted p-value < 0.01) between senescent and non-senescent cells for a given cell type. We then used these cell type-level log_2_ fold changes as enrichment scores to assess the cell type prevalence for these SAGs. The number of cell types in which each SAG was enriched in senescent cells with log_2_ fold-change > 0.5 was considered to be the cell type prevalence for that SAG.

To identify coordinated transcriptional programs, expression data for 3792 senescence-associated genes (SAGs) were analyzed across senescent cells from multiple donors, tissues, and cell types. Mean senescence-associated transcriptomic profiles of senescent cells were generated and used for visualaization of senescent transcriptomes on a UMAP. Single-cell senescence-associated transcriptomic profiles of senescent cells (48114 cells x 3792 genes) were used for co-expression analysis using the cNMF consensus non-negative matrix factorization implemented in the cNMF package (v-1.5.4)^163^. To identify co-expressed gene modules, consensus non-negative matrix factorization (cNMF) was applied to the raw gene expression count matrix across a range of matrix ranks (K = 10 to 50), with 10 iterations per K. The optimal number of factors, K = 34, was determined by evaluating the trade-off between reconstruction error and stability across K values. All factorization outputs from the 10 iterations at K = 34 were aggregated and subjected to density filtering, yielding 34 robust gene modules. Each module comprised the top 100 genes most strongly associated with the corresponding factor.

For each cNMF factor, the top 100 genes with the highest weights were selected for gene ontology (GO) analysis using the GSEApy package^162^ (v-1.1.3) and the GO_Biological_Processes_2023 database. For 100 genes in each cNMF factor, pathways with an adjusted p-value < 0.01 were retained. Among these, up to 10 pathways were selected per factor based on having at least five overlapping genes and ≤ 25% overlap with already selected pathways within the same list. A second round of filtering was performed to remove pathways with > 25% gene overlap across different gene lists to remove redundant pathways across factors. Pathway names, overlaps, GO-IDs, adjusted p-values, and associated genes per cNMF factors were then assembled. This analysis characterized the biological processes associated with each SAG module. For the 34 cNMF factors, a total of 31 distinct pathways were significantly associated with known biological processes (adjusted p-value < 0.01). Gene set enrichment analysis (GSEA) was then used to quantify enrichment of senescent associated pathways across different broad cell types. First, cell type-level log fold changes between senescent and non-senescent cells were calculated for all broad cell types. Gene sets corresponding to the ontologies representing senescence-associated pathways were then compiled using Biomart in the GSEApy package. These gene sets were then filtered to contain genes identified within the 3792 SAGs. Pre-ranked GSEA was then performed with cell type-level log fold changes and the filtered senescence-associated gene sets to characterize the enrichment of senescence-associated gene sets across cell types. GSEA results were filtered for FDR < 0.05 to select 17 senescence-associated pathways that were significantly enriched in one of the cell types. Normalized enrichment scores and the lead genes within each pathway were visualized on a Heatmap.

### Preprocessing for sex difference analysis

*Smart-seq Data Exclusion.* We filtered out cells associated with the Smart-seq protocol, reducing the overall cell numbers from 1,136,218 to 1,093,048..

*Low statistical power exclusion*. To ensure reliable comparison of sex differences across tissue and cell type reliably, we excluded tissue-cell type pairs that met either of the following criteria: (a) only one donor was represented, or (b) fewer than 100 cells were present in either the female or male group. This threshold was set to ensure sufficient statistical power for differential expression analyses.

*PCA density filter.* Outliers in the data were identified and removed using a Principal Component Analysis (PCA) density filter. Cells falling outside the high-density regions in the PCA space were considered outliers and excluded. This step ensured that extreme variations not representative of the underlying biological processes did not skew the results.

*Anatomical Position Consideration*: For each tissue and cell type, we checked for variability in anatomical position to avoid biases introduced by spatial heterogeneity across samples. If multiple anatomical positions were present for a given tissue-cell type, we checked for overlap between male and female donors. Only tissue-cell type pairs with shared anatomical positions between sexes were included in the analysis. If common anatomical positions were found, the data was further filtered to retain only samples from those positions. Additionally, a minimum of two male and two female donors was required to maintain sufficient statistical power. This step ensured that sex differences observed in gene expression were not confounded by anatomical variability across samples. The organ number for a rigor statistical power decreased from 28 to 13 organs.

Focusing on broad cell types for the sex-differential analysis, the number of tissue-cell type pairs decreased from 329 to 81 adequately represented groups as a result.

### Pseudo-Bulk differential gene expression analysis for sex differences

To assess differential gene expression between male and female samples at the tissue-cell type level, we employed a pseudo-bulk approach with edgeR (v-4.0.1)^164,165^, which has been recommended as an effective method to prevent false discoveries in datasets with covariates^166^. This method aggregates single-cell data to create bulk-like profiles, enhancing statistical power while accounting for variability among donors and other covariates. For each tissue-cell type pair, we extracted the subset of cells corresponding to that specific tissue and cell type from the filtered dataset. A PCA density filter was applied to remove outliers by performing PCA on the scVI latent space representations and calculating a density estimate. Cells falling below the density cutoff were excluded to ensure that extreme variations did not affect the results. We then processed the data at the donor level. For each donor within the tissue-cell type subset, we included only those donors with at least 50 cells to ensure sufficient representation. For donors meeting this criterion, we randomly sampled 500 cells with replacement from the donor’s cells to standardize the sample size across donors and reduce computational load. Clustering was performed on the donor-specific data using the Leiden algorithm with 30 nearest neighbors, 50 principal components and resolution 0.8, based on the scVI representations. This allowed us to capture intra-donor heterogeneity. For each cluster within the donor’s data, we aggregated the raw UMI counts to create pseudo-bulk expression profiles. This resulted in multiple pseudo-bulk samples per donor, capturing intra-donor heterogeneity. Each pseudo-bulk sample was then annotated with metadata including donor ID, tissue type, cell type, age, sex, ethnicity, and a unique cluster ID combining sex, donor ID, and Leiden cluster label. Genes were filtered to retain those expressed in at least 10% of the cells within the dataset, ensuring that lowly expressed genes did not skew the analysis. Key genes of interest, such as sex chromosome-linked genes, were retained regardless of their expression levels. The pseudo-bulk samples from all donors for each tissue-cell type pair were concatenated to create a comprehensive dataset for differential expression analysis.

Differential expression between male and female pseudo-bulk samples was assessed using edgeR’s generalized linear model (GLM) framework, which models count data using negative binomial distributions. The model included covariates such as donor age and batch effects where applicable to account for potential confounding factors. Genes with an adjusted p-value (Benjamini-Hochberg correction) less than 0.05 and an absolute log_2_ fold change greater than 0.25 were considered significantly differentially expressed.

### Comparison between Tabula Sapiens and GTEx

To directly compare the differentially expressed genes identified from the Tabula Sapiens and the Genotype-Tissue Expression (GTEx) project, we performed a pseudo-bulk analysis of the Tabula Sapiens data at the tissue level. This approach is similar to the pseudo-bulk differential gene expression analysis described previously but focuses on aggregating gene expression data per tissue rather than per tissue-cell type pair.

For the GTEx data, we obtained publicly available results for differentially expressed (DE) genes between male and female samples across various tissues. These results were sourced directly from the GTEx project, which provides pre-processed differential expression analyses performed on bulk RNA-seq data from healthy human tissues. By using the publicly available DE gene lists from GTEx, we ensured that the comparison between Tabula Sapiens and GTEx was based on consistent and standardized analyses performed by the GTEx consortium. To compare the differentially expressed genes identified from Tabula Sapiens and GTEx, we matched genes based on their gene symbols and aligned tissues between the two datasets. Overlaps between the lists of differentially expressed genes were assessed using Venn diagrams.

### Searchable extended metadata

Electronic Medical Records (EMRs) for 22 of 24 donors were obtained as PDF files from each hospital by Donor Network West, who manually de-identified each record. Our goal in developing ChatTS was to create a searchable database of extended donor metadata by feeding this information to an LLM to generate customized metadata tables for the user.

The EMRs had a mean±SD of 13±2 pages per donor, totaling 291 pages across all donors with EMRs available. Charts contained medical and social history taken at admission, and the full course of treatment following admission, until the donor was pronounced brain dead. This included positive and negative test results, values recorded for tests, other tabular and time-series information, as well as doctor notes. The initial size, scope, and format of the database was beyond current interpretability capabilities of LLMs, including the ability to understand tabular and time series data. Furthermore, their initial size was beyond context window limits, seemed to lead to sporadic information retrieval, and quickly overwhelmed API rate limits. Thus, to decrease complexity and size, we opted to create a database consisting of only free text present in the EMRs. Therefore, we manually extracted all free text from the EMRs, dropping tabular or time series data. To further decrease database size, we removed negative test results, and missing values (entered in the EMR as “NA” or “?”). The final size of this database is roughly 352KB and it contains a single .txt file per-donor.

### ChatTS Application Architecture

A single .txt file is stored for each donor. A user is required to supply their own OpenAI API key for billing purposes, and then asks a question. The user can ask questions in two different modes, Data Retrieval and Chart Review. In either case, the input question is taken from the user and fed through an LLM in a first pass to transform the question into the form of asking for a single donor (for example, “Does anyone have a history of lupus” would be transformed to something like “Does this donor have a history of lupus”). Then, this question is added to an input prompt independently per-donor. Each input prompt contains 1) the user’s transformed question, 2) a single donor’s EMR as manually curated text, and 3) a set of instructions on how to answer the question. In Data Retrieval mode, this set of instructions is to simply return yes, no, or a single number. Meanwhile, in Chart Review mode, the instructions request the LLM to provide a detailed answer with supporting information from the EMR text. We also parallelize the API requests to decrease total processing time experienced by the user. In summary, when the user asks a question, 22 independent prompts are created (one per donor) and asked to the LLM in parallel. The results are then aggregated and added to a central metadata table that can be downloaded as a csv file. While we use OpenAI, we built the LLM integration flexibly and could use any LLM as backend.

### ChatTS Application Server

The application is written in Python and HTML and hosted on an Amazon EC2 instance to allow flexibility in scaling to meet user demand. The application backend makes requests to GPT-4o through OpenAI’s API. We have deposited the source code online at https://github.com/Harper-Hua/ChatTS.

## Supplementary Figures

**Supplementary Figure 1:**
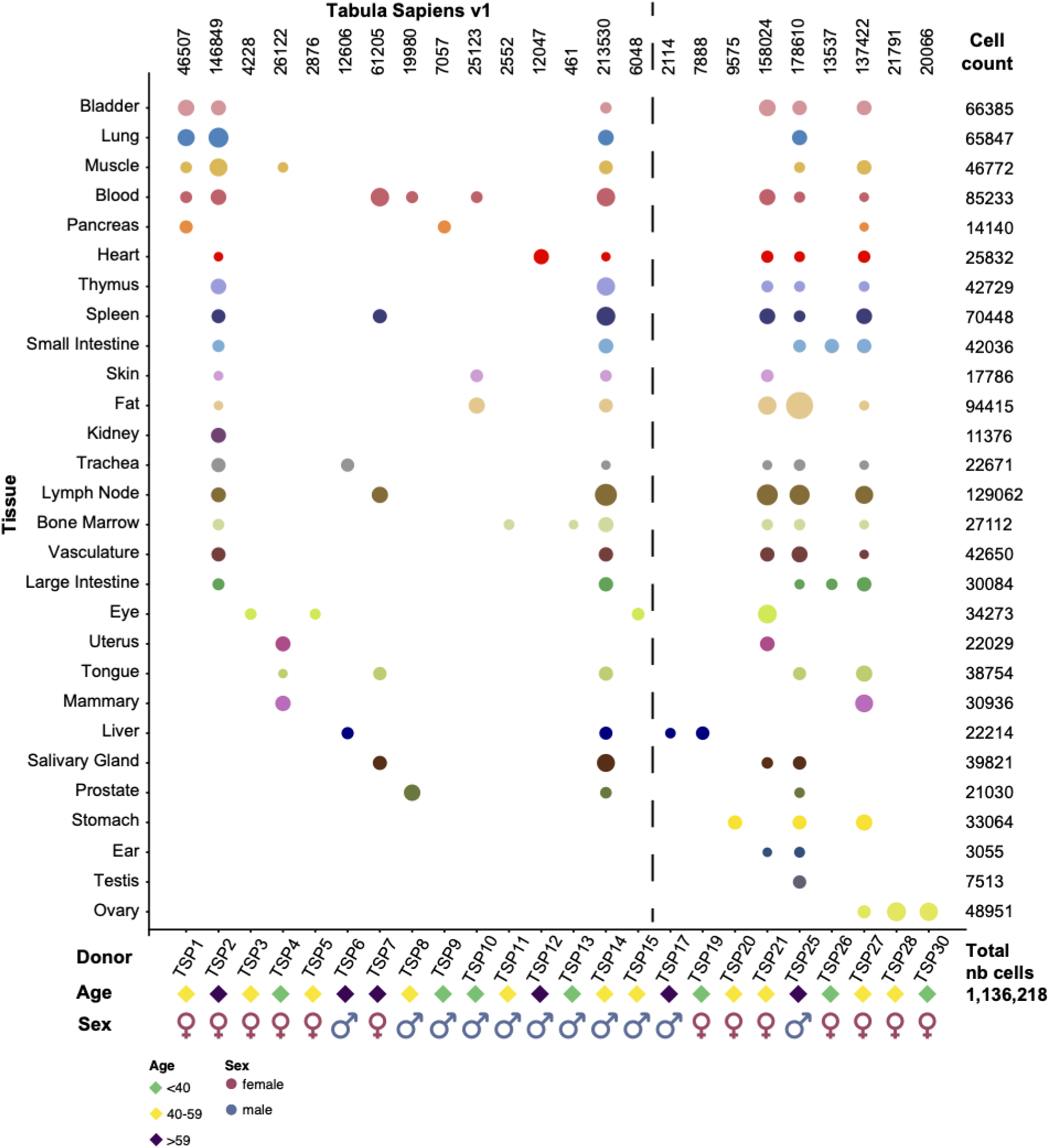
Bubble plot summarizing the distribution of cells across various tissues, donors, ages, and sexes from the Tabula Sapiens 2.0. Each row corresponds to a different tissue type, with tissues listed on the y-axis. The x-axis represents individual donors (TSP1 to TSP30), with symbols at the bottom indicating donor sex (pink circles for females, blue circles for males) and age group (green diamonds for donors under 40 years old, yellow diamonds for donors aged 40-59, and purple diamonds for donors over 59). The size of the bubbles represents the number of cells sampled from each tissue for each donor, and the color of the bubbles reflects the sex of the donor. On the far right, the total cell count for each tissue is listed, with the sum across all tissues amounting to 1,136,335 cells.

**Supplementary Figure 2.**
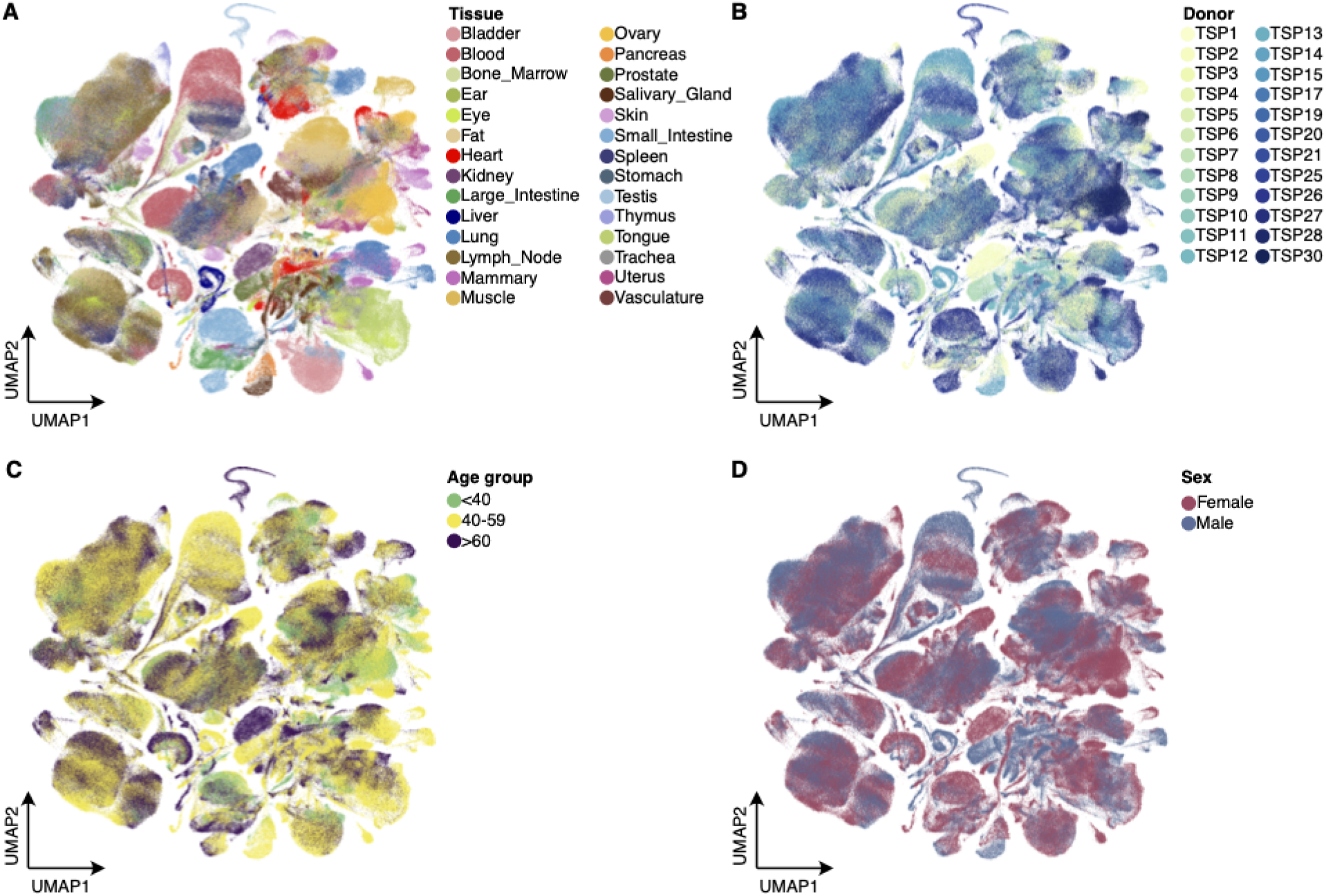
UMAP overview of key metadata variables in Tabula Sapiens 2.0. UMAP plots of the expanded Tabula Sapiens 2.0 colored by **A)** Tissue, **B)** Donor, **C)** Age, and **D)** Sex.

**Supplementary Figure 3.** Data Quality Control.

**Supplementary Figure 4.** Data Quality Control.

**Supplementary Figure 5.** Data Quality Control.

**Supplementary Figure 6.** Data Quality Control.

**Supplementary Figure 7.**
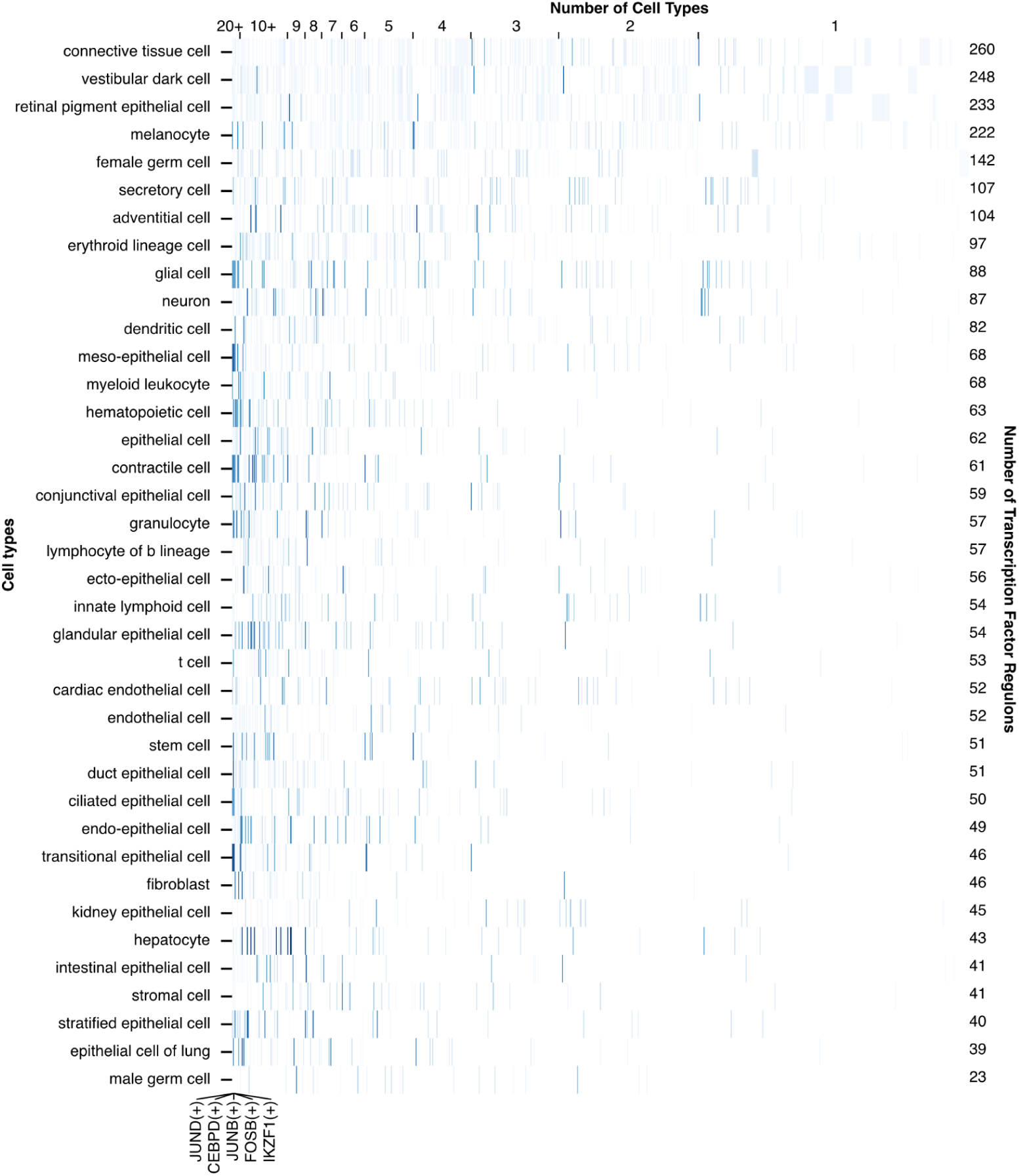
Transcription factor activity across cell types. 839 transcription factor regulons identified by SCENIC are shown in descending order by cell type specificity from left to right and cell types are listed in descending order by number of regulons identified for them. Darker shades correspond to higher average activity of a transcription factor within cells of a cell type.

**Supplementary Figure 8:**
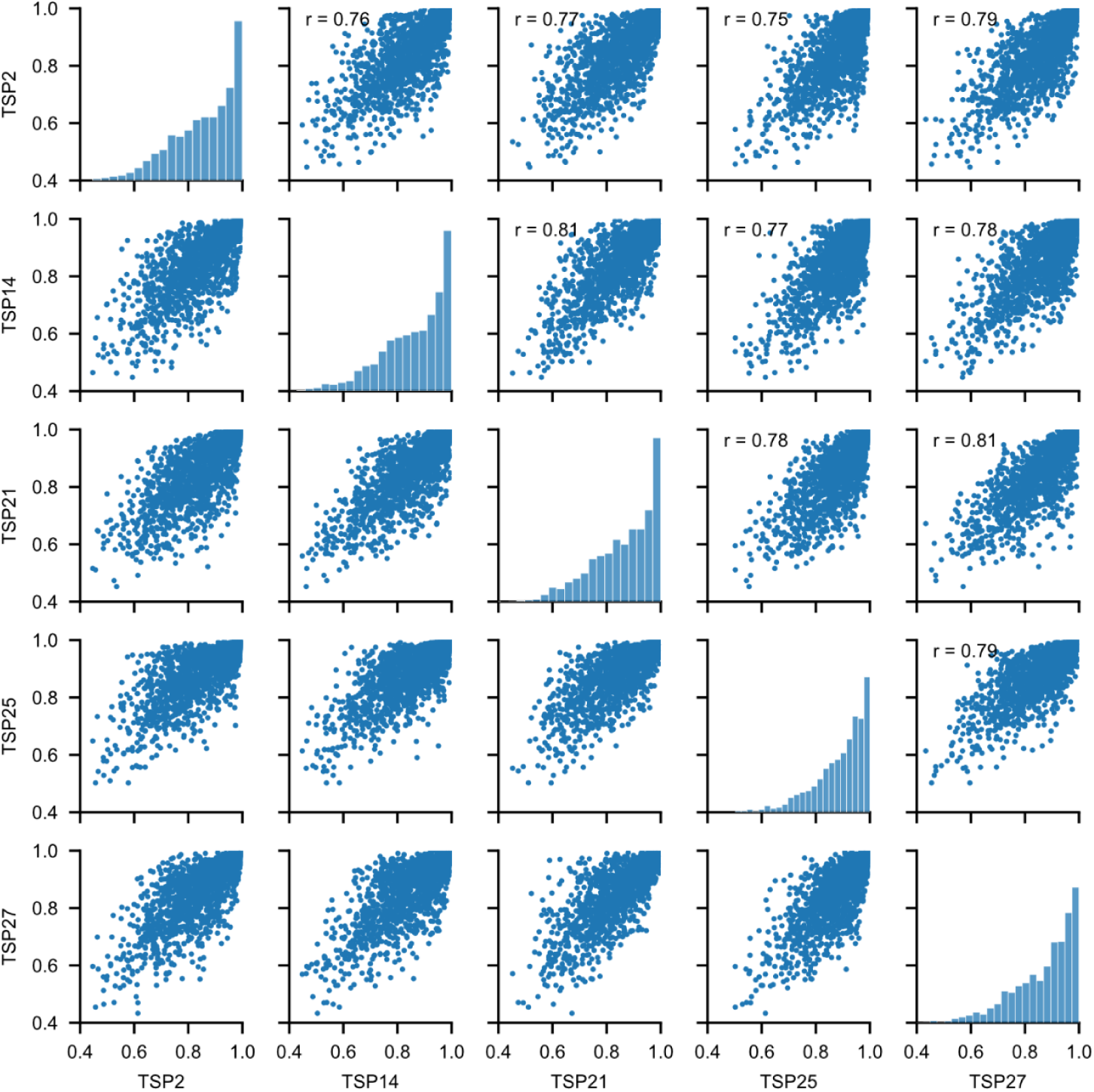
Inter-donor pearson correlation of τ specificity. Mean cell type expression was computed on each of donors TSP2, TSP14, TSP21, TSP25 and TSP27, followed by computation of τ cell type specificity for each TF and the pearson correlation between donors.

**Supplementary Figure 9:**
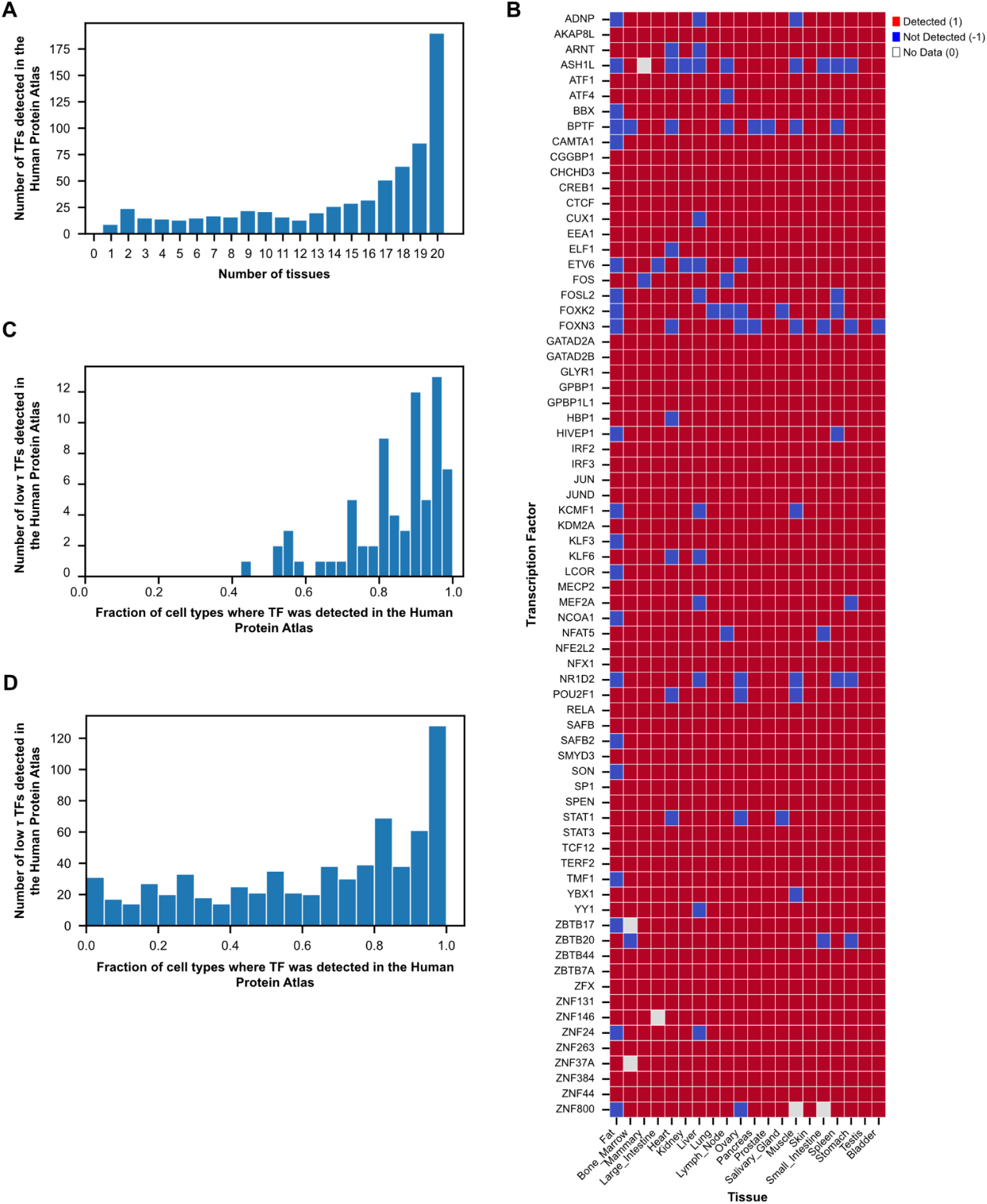
Human Protein Atlas immuno detection of transcription factors on a cell-type basis. A) Distribution of the number of tissues in which each of the 699 common TF proteins were detected. **B**) Binary heatmap of detected vs not detected for 72 of the lowest 100 τ TF’s in the 20 common tissues. **C)** Distribution of 72 of the 100 lowest τ TF’s across the fraction of 32 common cell types in which the TF protein was detected. **D)** The distribution of all 699 TF’s tested across the fraction of cell types.

**Supplementary Figure 10:**
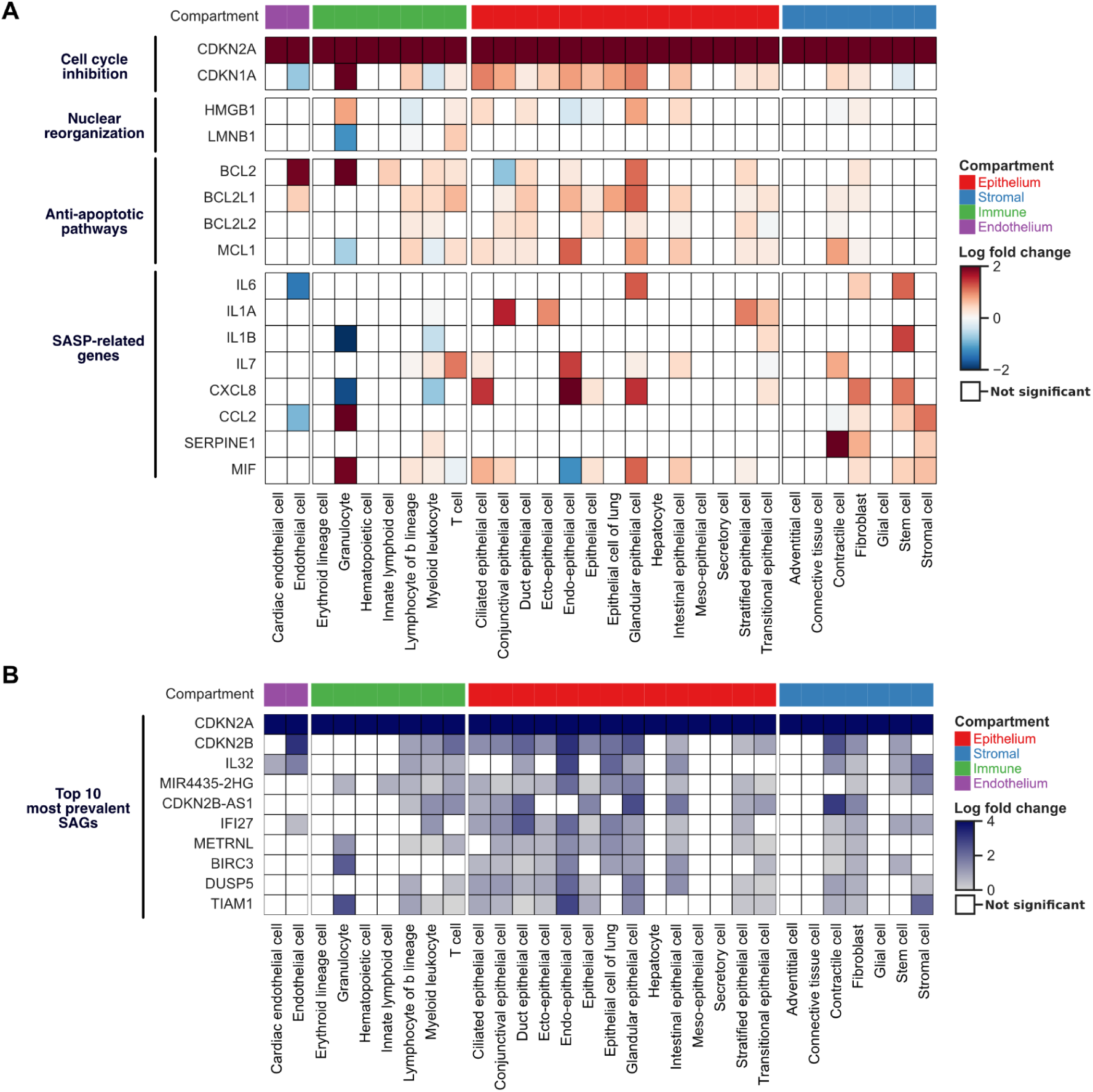
Molecular phenotype of senescent cells across human tissues. **A)** Heat map showing the log_2_ fold changes for senescence hallmark genes, which were statistically different between senescent and non-senescent cells across broad cell types (adjusted *p*-value < 0.01). The list of hallmark genes was compiled according to SenNet guidelines. Genes are grouped into four distinct hallmarks of senescence, labeled on the left. Broad cell types were grouped into four compartments, as indicated by the color bar at the top. **B)** Heat map showing the log_2_ fold changes for the most universal senescence-associated genes (SAGs), which were statistically different between senescent and non-senescent cells across broad cell types (adjusted *p*-value < 0.01). Broad cell types were grouped into four compartments, as indicated by the color bar at the top.

**Supplementary Figure 11:**
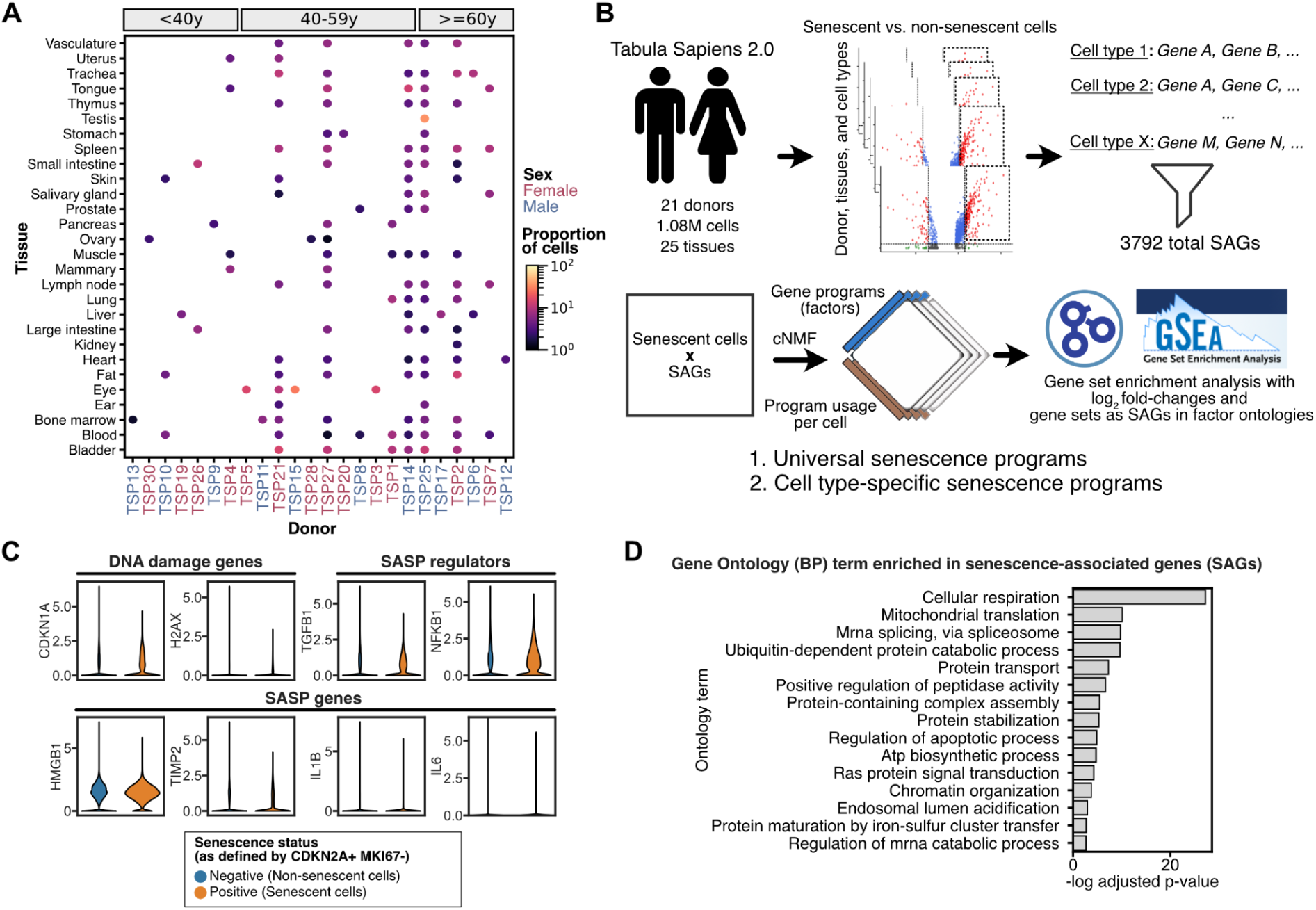
Senescent cells distribution and phenotypes across human tissues. **A)** Dot plot showing the proportion of senescent cells, defined as *CDKN2A*+ *MKI67*- cells, across all tissues and donors. The donor names were colored by sex and were split by age in the annotation bar on top. **B)** Workflow chart showing the methodological steps used to characterize universal and non-universal senescence-associated genes (SAGs). **C)** Violin plots showing the expression of various canonical SAGs in senescent (*CDKN2A*+ *MKI67*-) and non-senescent cells (*CDKN2A*-) in Tabula Sapiens 2.0 dataset. **D)** Bar plot showing top 15 ontology terms enriched for 3972 senescence-associated genes.

**Supplementary Figure 12:**
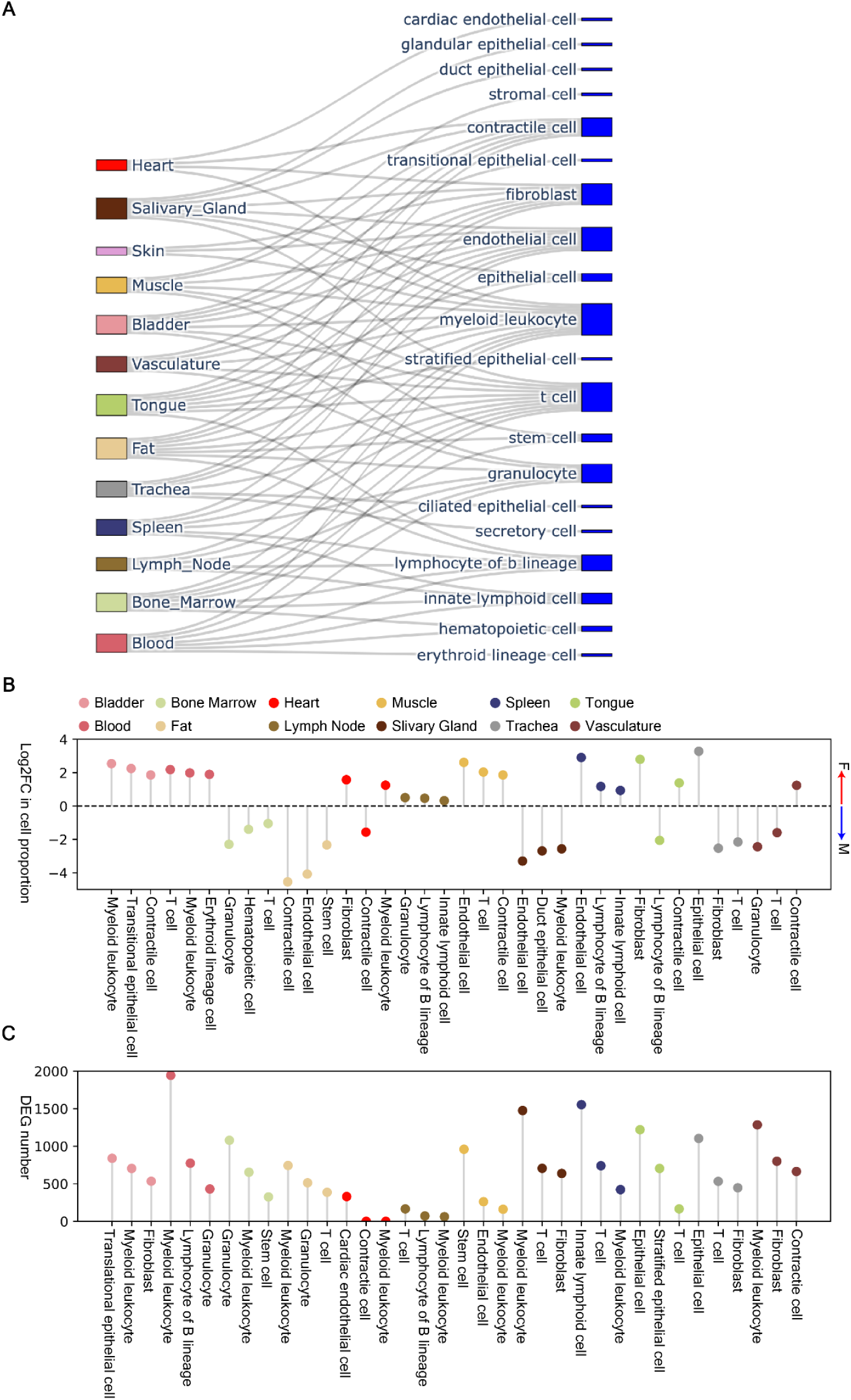
Overview of Tabula Sapiens 2.0 data for sex-biased analysis. **A)** Sankey diagram showing the flow of tissue types (left) into specific cell types (right) analyzed for sex-biased expression. Tissues are color-coded and connected to major cell classes, such as cardiac endothelial cells, fibroblasts, contractile cells, epithelial cells, and granulocytes. **B)** Lollipop plot of the log fold change (logFC) for sex-biased gene expression, comparing male and female expression levels across different cell types. Positive logFC values indicate female-biased genes (red), while negative logFC values indicate male-biased genes (blue). **C)** Corresponding plot displaying the total number of differentially expressed genes (DEGs) for each cell type. The color of the dots represents the tissue origin, matching the color legend from panel B.

**Supplementary Figure 13.**
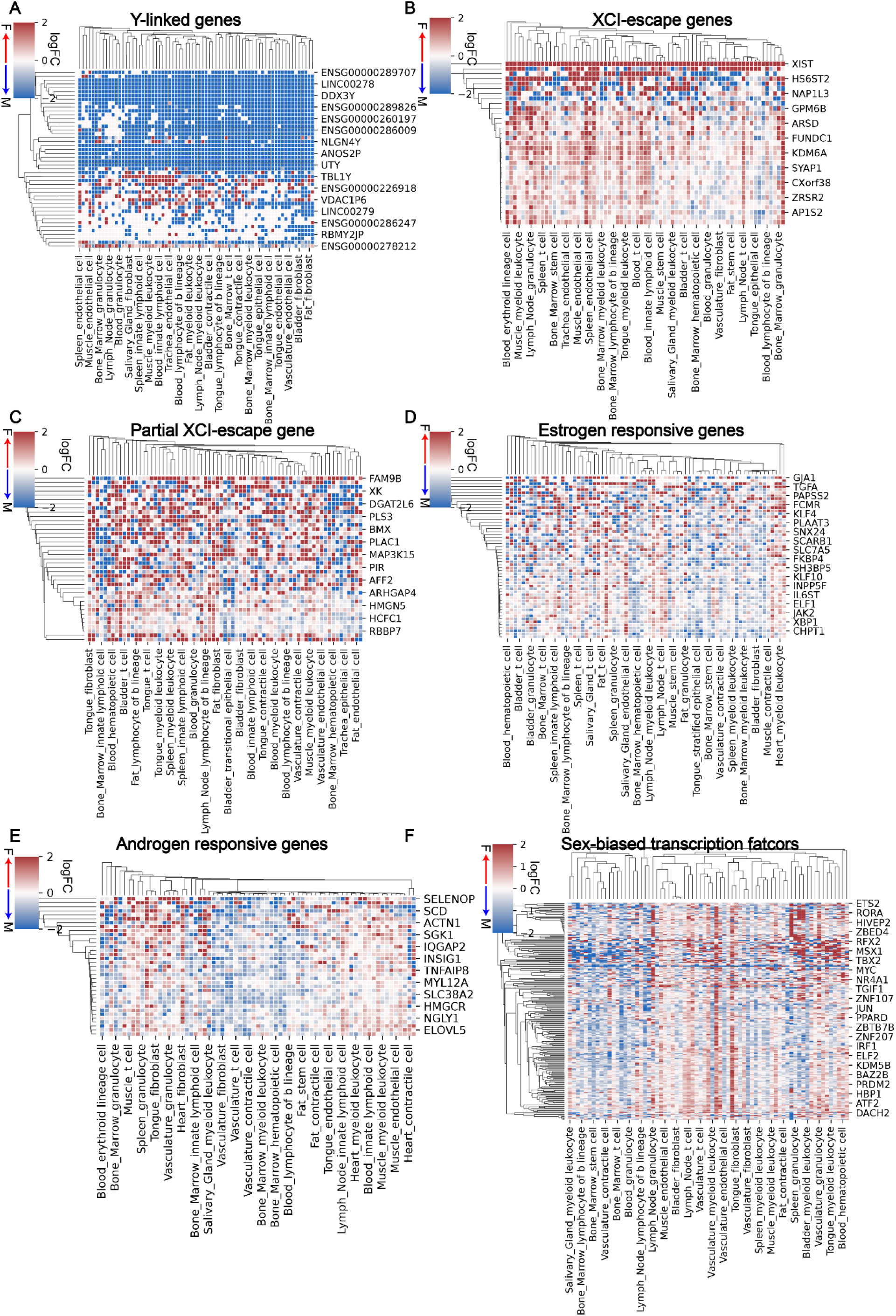
Mean log fold change value of differential gene expression between female to male across. **A)** Y-linked genes, **B)** chrX inactivation escape genes, **C)** partial escape genes, **D)** estrogen responsive genes and **E)** androgen responsive genes, **F)** sex-biased transcription factors.

**Supplementary Figure 14.**
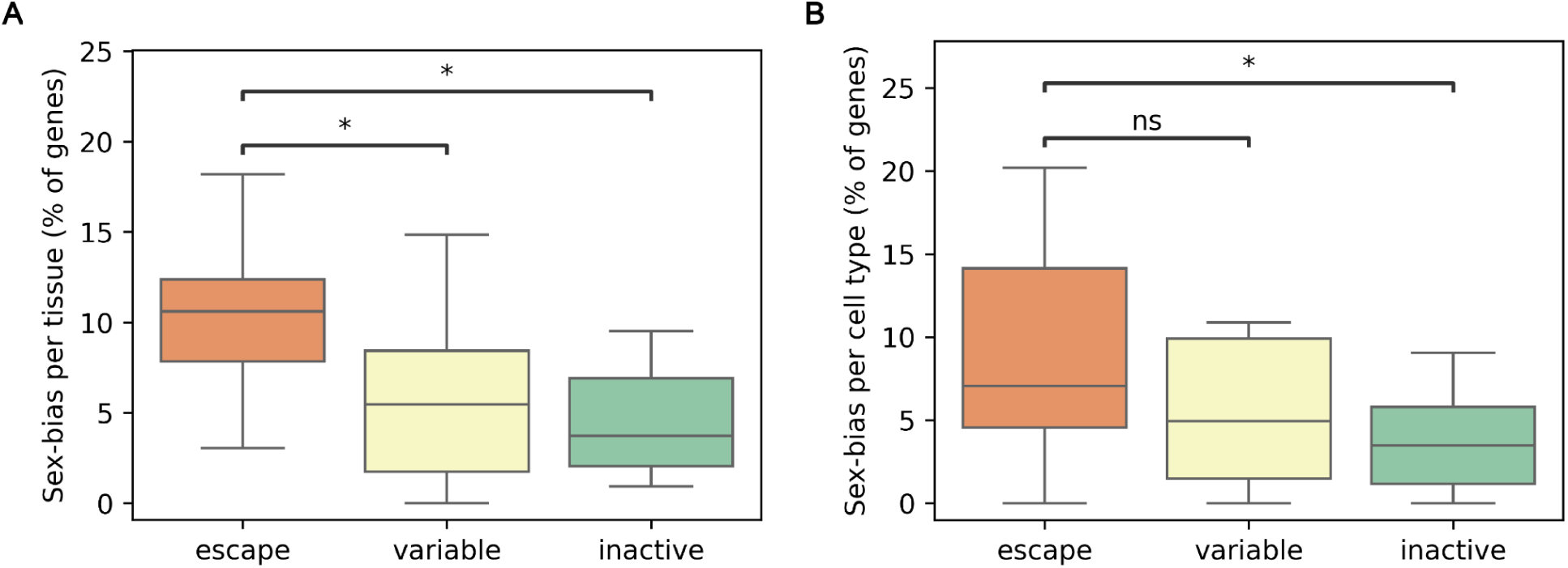
Barplot of sex-biased per (A) tissue and (B) cell type across XCI escape, variable and inactive status.

**Supplementary Figure 15:**
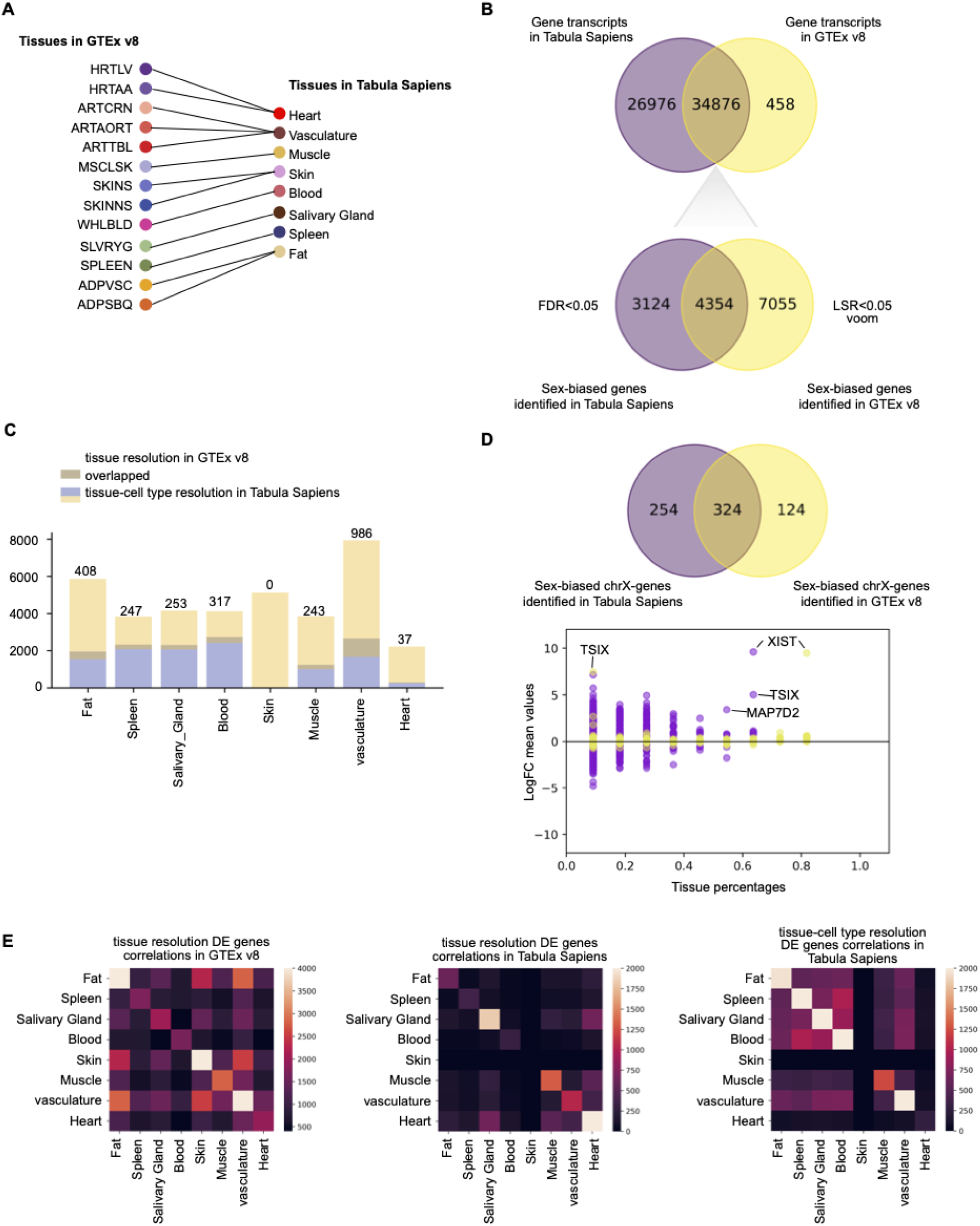
Comparison of differential gene expression across the Tabula Sapiens 2.0 and GTEx Projects. **A)** Mapping of overlapping tissues between GTEx v8 and Tabula Sapiens 2.0. Each tissue present in GTEx (left) is connected to its corresponding tissue in Tabula Sapiens (right), including tissues like heart, vasculature, muscle, skin, blood, and fat. **B)** Venn diagrams comparing gene transcripts between Tabula Sapiens 2.0 and GTEx v8. Top: The number of gene transcripts shared between both datasets. Bottom: Comparison of sex-biased genes identified at an FDR < 0.05 in Tabula Sapiens and GTEx using voom analysis. **C)** Bar plot showing the number of sex-biased genes identified in GTEx (tissue resolution) versus Tabula Sapiens (tissue resolution and tissue-cell type resolution). The orange bars represent genes identified in Tabula Sapiens, yellow bars show genes from GTEx, and the gray bars represent the overlap between the two datasets. **D)** Top: Venn diagram comparing sex-biased X chromosome genes identified in Tabula Sapiens and GTEx, including notable genes such as TSIX and XIST. Bottom: Log fold change (logFC) values for X-linked genes across tissues, highlighting TSIX and MAPT02. **E)** Heatmaps comparing the correlations of differentially expressed genes at the tissue resolution between GTEx v8 (left) and Tabula Sapiens v2 (center), and at the tissue-cell type resolution in Tabula Sapiens 2.0 (right).

**Supplementary Figure 16.**
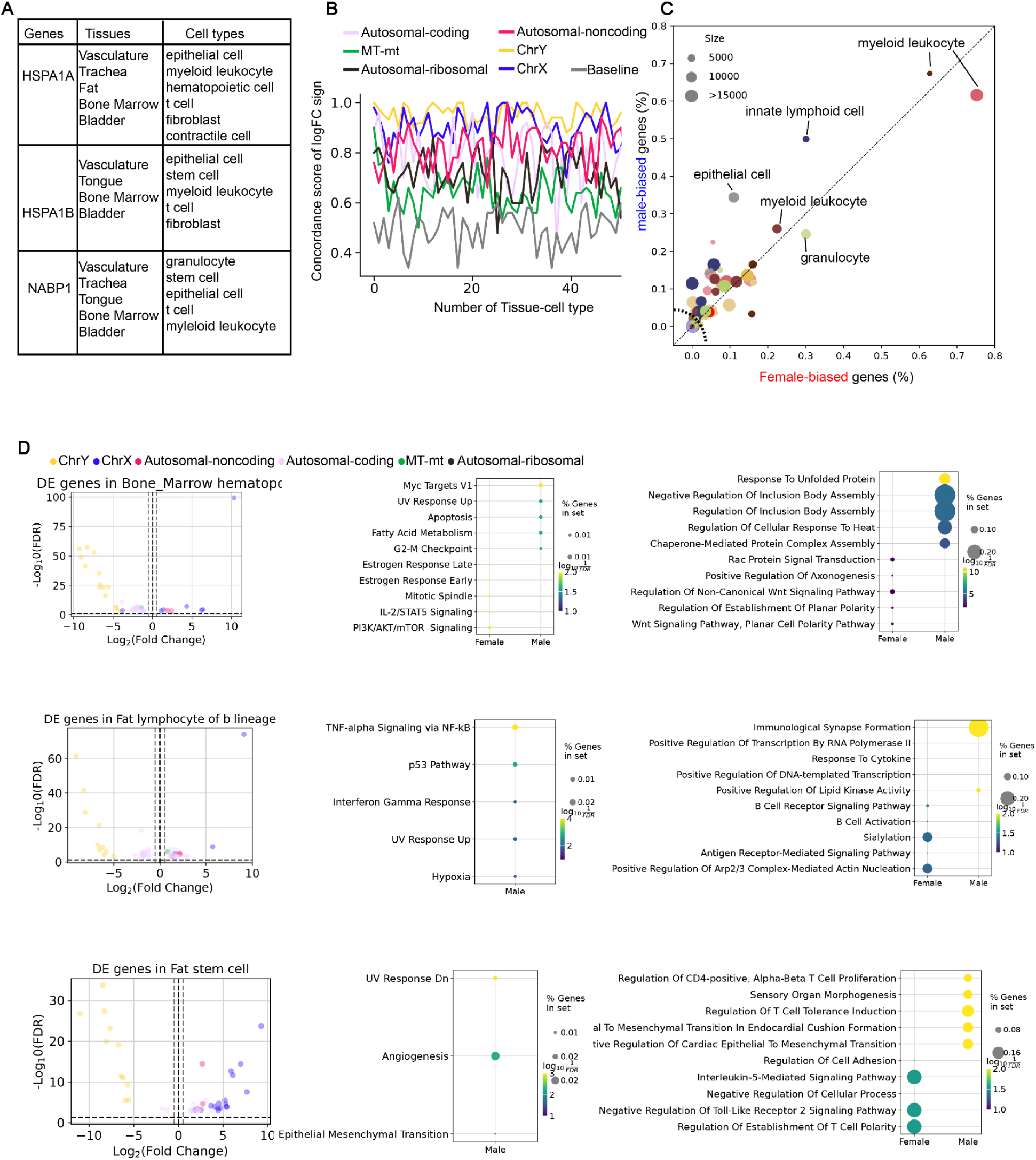
Analysis of sex-biased gene expression across tissue-cell types. **A)** Table showing representative sex-biased genes identified across various tissues and cell types in the Tabula Sapiens dataset. Tissues include blood, heart, lymph node, salivary gland, and spleen; cell types include B lymphocytes, fibroblasts, contractile cells, and myeloid leukocytes. **B)** Line plot showing concordance scores of differentially expressed genes (DEGs) across tissue-cell types. Concordance is measured for various gene categories, including autosomal-coding, autosomal-noncoding, mitochondrial, chromosome Y (chrY), and chromosome X (chrX). The baseline score is included for comparison. **C)** Scatter plot displaying the percentage of male-biased versus female-biased genes across cell types. The size of the dots corresponds to the number of sex-biased genes identified in each cell type. Major cell types include B lymphocytes, fibroblasts, contractile cells, and T cells. The plot highlights the distribution of male-biased genes (in blue) and female-biased genes (in red). **D)** Left: Volcano plots illustrating sex-biased genes (logFC) and statistical significance (-log10(FDR)) for select cell types, including T cells, lymphocytes of B lineage, and heart contractile cells. Right: Bubble plots displaying enriched biological pathways for male-and female-enriched genes, with pathway terms related to oxidative phosphorylation, mitotic spindle, UV response, and cellular respiration. Bubble size corresponds to the number of genes involved, and the color represents the logFC direction (female or male).

**Supplementary Figure 17:**
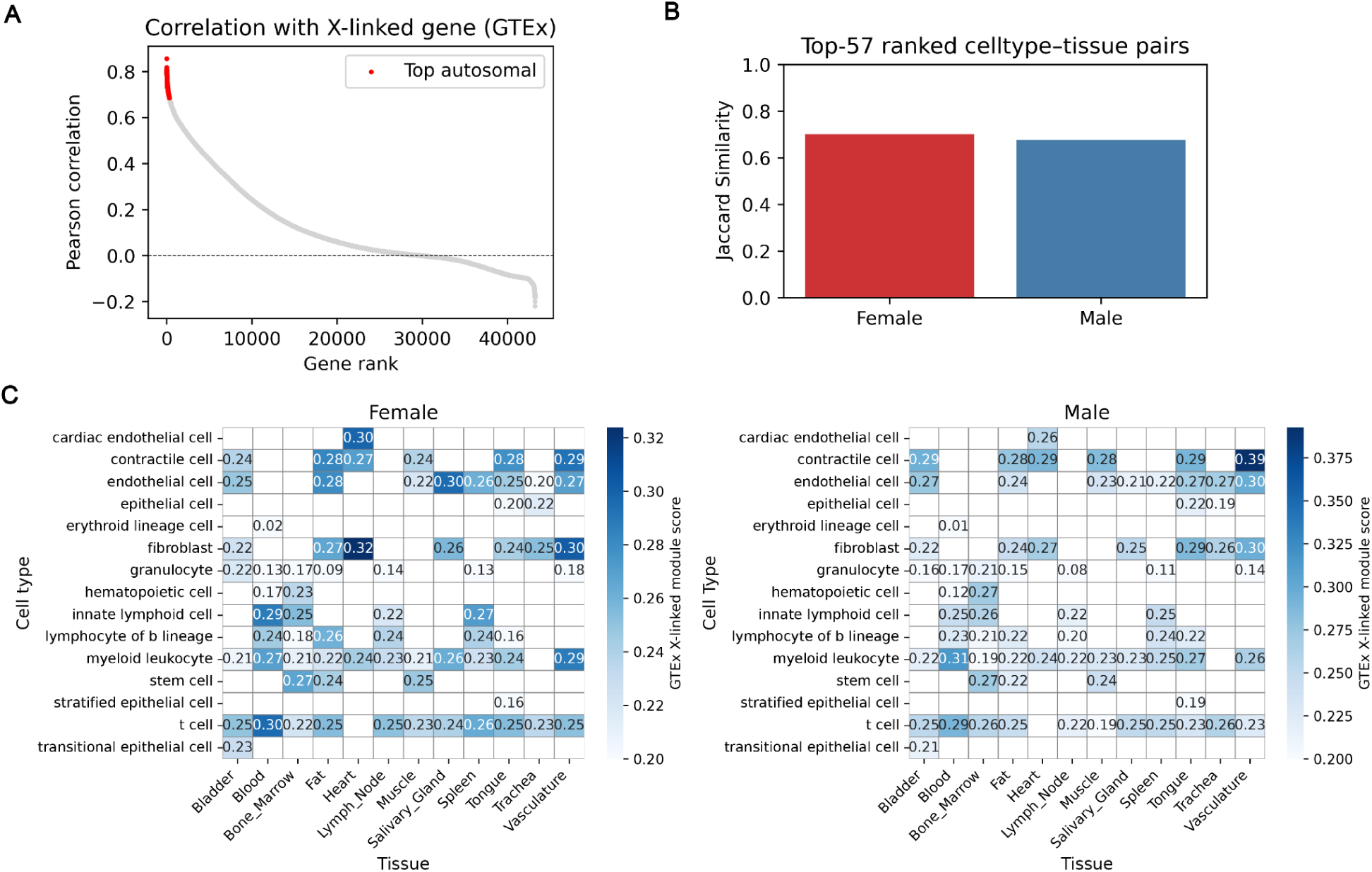
Autosomal genes co-expressed with X-linked genes across tissues and cell types identified in GTEx are correlated with sex-biased tissue-cell type pairs in Tabula Sapiens 2.0. **A)** Top-ranking autosomal genes among the top 300 genes in GTEx coexpressed with X-linked genes. **B)** Jaccard similarity between the top 57 sex-biaed tissue-cell type pairs in GTEx and Tabula Sapiens. **C)** Heatmap of gene expression in Tabula Sapiens 2.0 for autosomal genes identified in GTEx co-expressed with X-linked genes across tissues and cell types.

## Full author list

### Overall project direction and coordination

Robert C Jones^1^

Mark Krasnow^2,3^

Angela Oliveira Pisco^4^

Stephen R. Quake^1,5,6^

Julia Salzman^2,7^

Nir Yosef^4,8,9^

### Manuscript writing

George Crowley^1^

Siyu He^1,7^

Robert C Jones^1^

Madhav Mantri^1^

Angela Olivera Pisco^4^

Stephen R. Quake^1,5,6^

Anton Thieme^1^

### Donor recruitment

Jessie Aguirre^10^

Ron Garner^10^

Sal Guerrero^10^

William Harper^10^

Resham Irfan^10^

Sophia Mahfouz^10^

Ravi Ponnusamy^10^

Bhavani A. Sanagavarapu^10^

Ahmad Salehi^10^

Ivan Sampson^10^

Chloe Tang^10^

### Surgeons

Alan G. Cheng^11^

James M. Gardner^12,13^

Burnett Kelly^10,14^

Thurman Slone^10^

Zifa Wang^10^

### Logistical coordination

Anika Choudhury^1,4^

Sheela Crasta^1^

George Crowley^1^

Chen Dong^1,4^

Marcus L. Forst^1^

Douglas E. Henze^1^

Jaeyoon Lee^1^

Maurizio Morri^4^

Angela Oliveira Pisco^4^

Serena Y. Tan^15^

Sevahn K. Vorperian^16,17^

Lynn Yang^1,4^

### Organ processing

Marcela Alcántara-Hernádez^18^

Julian Berg^19^

Dhruv Bhatt^20^

Sara Billings^11^

Andrès Gottfried-Blackmore^18,21^

Jamie Bozeman^20^

Simon Bucher^22^

Elisa Caffrey^23^

Amber Casillas^24^

Rebecca Chen^23^

Anika Choudhury^1,4^

Matthew Choi^22^

Sheela Crasta^1^

Rebecca N. Culver Ivana Cvijovic^1,5^

Ke Ding^26^

Hala Shakib Dhowre^27^

Hua Dong^26^

Kenneth Donaville^22^

Lauren Duan^19^

Xiaochen Fan^20^

Mariko H. Foecke^24^

Francisco X. Galdos^19^

Eliza A. Gaylord^24^

Karen Gonzales^20^

Wiliam R. Goodyer ^28^

Michelle Griffin^29^

Yuchao Gu^2,30,31^

Shuo Han^26^

Jun Yan He^23^

Paul Heinrich^19^

Rebeca Arroyo Hornero^18^

Keliana Hui^23^

Juan C. Irwin^24^

SoRi Jang^2^

Annie Jensen^19,32^

Saswati Karmakar^15,25^

Jengmin Kang^33^

Hailey Kang^22^

Soochi Kim^33^

Stewart J. Kim^2,30,31^

William Kong^26^

Mallory A. Laboulaye^26^

Daniel Lee^19^

Gyehyun Lee^34^

Elise Lelou^22^

Anping Li^26^

Baoxiang Li^27^

Wan-Jin Lu^26^

Hayley Raquer-McKay^18^

Elvira Mennillo^34^

Lindsay Moore^35^

Elena Montauti^23^

Karim Mrouj^26^

Shravani Mukherjee^27^

Patrick Neuhöfer ^2,30,31^

Saphia Nguyen^22^

Honor Paine^22^

Jennifer B. Parker^26,29^

Julia Pham^23^

Kiet T. Phong^36^

Pratima Prabala^20^

Zhen Qi^26^

Joshua Quintanilla^19,32^

Iulia Rusu^34^

Ali Reza Rais Sadati^19^

Bronwyn Scott^27^

David Seong^18^

Hosu Sin^37^

Hanbing Song^38^

Bikem Soyur^24^

Sean Spencer^18,21^

Varun R. Subramaniam^27^

Michael Swift^1^

Aditi Swarup^27^

Greg Szot^12,13^

Aris Taychameekiatchai^22^

Emily Trimm^20^

Stefan Veizades^19,32^

Sivakamasundari Vijayakumar^26^

Kim Chi Vo^24^

Tian Wang^35^

Timothy Wu^2^

Yinghua Xie^19,32^

William Yue^22^

Zue Zhang^2^

### Sequencing

Angela Detweiler^4^

Honey Mekonen^4^

Norma F. Neff^4^

Sheryl Paul^4^

Amanda Seng^4^

Jia Yan^4^

### Biobanking and Histology

Deana Rae Crystal Colburg^39^

Chen Dong^1,4^

Balint Laszlo Forgo^15^

Luca Ghita^18^

Frank McCarthy^40^

Serena Y. Tan^15^

Lynn Yang^1,4^

### cDNA and library prep

Aditi Agrawal^4^

Anika Choudhury^1,4^

Sheela Crasta^1^

Chen Dong^1,4^

Alina Isakova^1^

Jaeyoon Lee^1^

Maurizio Morri^4^

Kavita Murthy^1^

Alexandra Psaltis^1^

Wenfei Sun^1^

Jia Yan^4^

### Data Analysis

Kyle Awayan^4^

Pierre Boyeau^41^

Robrecht Cannoodt^42–44^

George Crowley^1^

Leah Dorman^4^

Samuel D’Souza^4^

Can Ergen^8,41^

Siyu He^7^

Justin Hong^45^

Harper Hua^7^

Robert C Jones^1^

Jaeyoon Lee^1^

Madhav Mantri^1^

Erin McGeever^4^

Antoine de Morree^33,46,47^

Angela Oliveira Pisco^4^

Luise A. Seeker^1^

Alexander J. Tarashansky^4^

Anton Thieme^1^

### Expert cell type annotation

Marcela Alcántara-Hernádez^18^

Simon Bucher^22^

Ivana Cvijovic^1,5^

Hala Shakib Dhowre^27^

Ke Ding^26^

Hua Dong^26^

Xiaochen Fan^20^

Mariko H. Foecke^24^

Eliza A. Gaylord^24^

Astrid Gillich^2^

Wiliam R. Goodyer ^28^

Andrès Gottfried-Blackmore^18,21^

Rebeca Arroyo Hornero^18^

Taha A. Jan^48^

SoRi Jang^2^

Soochi Kim^33^

Saswati Karmakar^15,25^

Stewart J. Kim^2,30,31^

William Kong^26^

Mallory A. Laboulaye^26^

Angela Ling^48^

Wan-Jin Lu^26^

Antoine de Morree^46^

Abhishek Murti^22^

Patrick Neuhöfer^2,30,31^

Kiet T. Phong^36^

Hayley Raquer-McKay^18^

Nikita Sajai^22^

Ryan M. Samuel^49^

David Seong^18^

Hosu Sin^37^

Hanbing Song^38^

Michael Swift^1^

Aris Taychameekiatchai^22^

Stefan Veizades^19,32^

Sivakamasundari Vijayakumar^26^

Juliane Winkler^50,51^

Timothy Wu^2^

Zue Zhang^2^

### Tissue expert principal investigators

Steven E. Artandi^2,30,31^

Philip A. Beachy^26,32,52^

Alan G. Cheng^11^

Mike F. Clarke^26^

Zev Gartner^4,53^

Linda C. Giudice^54^

Franklin W. Huang^38,55^

Juliana Idoyaga^18,56^

Michael G. Kattah^34^

Mark Krasnow^2,3^

Christin S. Kuo^57^

Diana J. Laird^24^

Michael T. Longaker^26,58^

Patricia Nguyen^19,32,59^

David Y. Oh^23^

Stephen R. Quake^1,5,6^

Thomas A. Rando^33^

Kristy Red-Horse^20^

Bruce Wang^22^

Albert Y. Wu^27^

Sean M. Wu^19,32^

Bo Yu^37^

James Zou^7,60^

### Affiliations

1. Department of Bioengineering, Stanford University; Stanford, CA, USA.

2. Department of Biochemistry, Stanford University School of Medicine, Stanford, CA, USA.

3. Howard Hughes Medical Institute, USA.

4. Chan Zuckerberg Biohub, San Francisco, CA, USA.

5. Department of Applied Physics, Stanford University, Stanford, CA, USA.

6. The Chan Zuckerberg Initiative, Redwood City, CA, USA.

7. Department of Biomedical Data Science, Stanford University, Stanford, CA, USA.

8. Center for Computational Biology, University of California Berkeley, Berkeley, CA, USA.

9. Ragon Institute of MGH, MIT and Harvard, Cambridge, MA, USA.

10. Donor Network West, San Ramon, CA, USA.

11. Department of Otolaryngology-Head and Neck Surgery, Stanford University School of Medicine, Stanford, California, USA.

12. Department of Surgery, University of California San Francisco, San Francisco, CA, USA.

13. Diabetes Center, University of California San Francisco, San Francisco, CA, USA.

14. DCI Donor Services, Sacramento, CA, USA.

15. Department of Pathology, Stanford University School of Medicine, Stanford, CA, USA.

16. Department of Chemical Engineering, Stanford University, Stanford, CA, USA.

17. Sarafan ChEM-H, Stanford University, Stanford, CA, USA.

18. Department of Microbiology and Immunology, Stanford University School of Medicine, Stanford, CA, USA.

19. Stanford Cardiovascular Institute, Stanford CA, USA.

20. Department of Biology, Stanford University, Stanford, CA, USA.

21. Division of Gastroenterology, Department of Medicine, Stanford University School of Medicine, Stanford, CA, USA.

22. Department of Medicine and Liver Center, University of California San Francisco, San Francisco, CA, USA.

23. Division of Hematology/Oncology, Department of Medicine, University of California San Francisco, San Francisco, CA, USA.

24. Department of Ob/Gyn and Reproductive Sciences, Eli and Edythe Broad Center for Regeneration Medicine and Stem Cell Research, University of California, San Francisco.

25. Department of Genetics, Stanford University School of Medicine, Stanford, CA, USA.

26. Institute for Stem Cell Biology and Regenerative Medicine, Stanford University School of Medicine, Stanford, CA, USA.

27. Department of Ophthalmology, Stanford University School of Medicine, Stanford, CA, USA.

28. Department of Pediatrics, Division of Cardiology, Stanford University School of Medicine, Stanford, CA, USA.

29. Department of Surgery, Division of Plastic and Reconstructive Surgery, Stanford University School of Medicine, Stanford, CA, USA.

30. Stanford Cancer Institute, Stanford University School of Medicine, Stanford, CA, USA.

31. Department of Medicine, Division of Hematology, Stanford University School of Medicine, Stanford, CA, USA.

32. Department of Medicine, Division of Cardiovascular Medicine, Stanford University, Stanford, CA, USA.

33. Department of Neurology and Neurological Sciences, Stanford University School of Medicine, Stanford, CA, USA.

34. Division of Gastroenterology, Department of Medicine, University of California, San Francisco, San Francisco, CA, USA.

35. Division of Pediatric Otolaryngology Stanford University School of Medicine, Stanford, CA, USA.

36. Department of Bioengineering and Therapeutic Sciences, University of California, San Francisco, San Francisco, CA, USA.

37. Department of OB/GYN Stanford University, Palo Alto, CA, USA.

38. Division of Hematology and Oncology, Department of Medicine, Bakar Computational Health Sciences Institute, Institute for Human Genetics, University of California San Francisco, San Francisco, CA, USA.

39. Stanford Health Care, Stanford CA, USA.

40. Mass Spectrometry Platform, Chan Zuckerberg Biohub, Stanford, CA, USA.

41. Department of Electrical Engineering and Computer Sciences, University of California Berkeley, Berkeley, CA, USA.

42. Data Intuitive, Flanders, Belgium.

43. Data Mining and Modelling for Biomedicine group, VIB Center for Inflammation Research, Ghent, Belgium.

44. Department of Applied Mathematics, Computer Science, and Statistics, Ghent University, Ghent, Belgium.

45. Department of Computer Science, Columbia University; New York, NY, USA.

46. Department of Biomedicine, Aarhus University, Aarhus, Denmark.

47. Paul F. Glenn Center for the Biology of Aging, Stanford University School of Medicine, Stanford, CA, USA.

48. Department of Otolaryngology, Vanderbilt University Medical Center, Nashville, TN, USA.

49. Department of Cellular Molecular Pharmacology, University of California, San Francisco, San Francisco, CA, USA.

50. Department of Cell & Tissue Biology, University of California San Francisco, San Francisco, CA, USA.

51. Center for Cancer Research Medical University of Vienna Borschkegasse 8a 1090 Vienna, Austria.

52. Department of Urology, Stanford University School of Medicine, Stanford, CA, USA.

53. Department of Pharmaceutical Chemistry, University of California San Francisco, San Francisco, CA.

54. Center for Reproductive Sciences, Department of Obstetrics, Gynecology and Reproductive Sciences, University of California San Francisco, San Francisco, CA, USA.

55. Division of Hematology/Oncology, Department of Medicine, San Francisco Veterans Affairs Health Care System, San Francisco, CA, USA.

56. Pharmacology and Molecular Biology Departments, Schools of Medicine and Biological Sciences, University of California, San Diego, CA, USA.

57. Department of Pediatrics, Division of Pulmonary Medicine, Stanford University, Stanford, CA, USA.

58. Department of Surgery, Stanford University School of Medicine, Stanford, CA, USA.

59. Veterans Affairs Palo Alto Health Care System, Palo Alto, CA, USA.

60. Department of Computer Science, Stanford University, Stanford, CA.

