## Supplementary figures and images for "Tabula Sapiens reveals transcription factor expression, senescence effects, and sex-specific features in cell types from 28 human organs and tissues"

### Supp. Figure 1

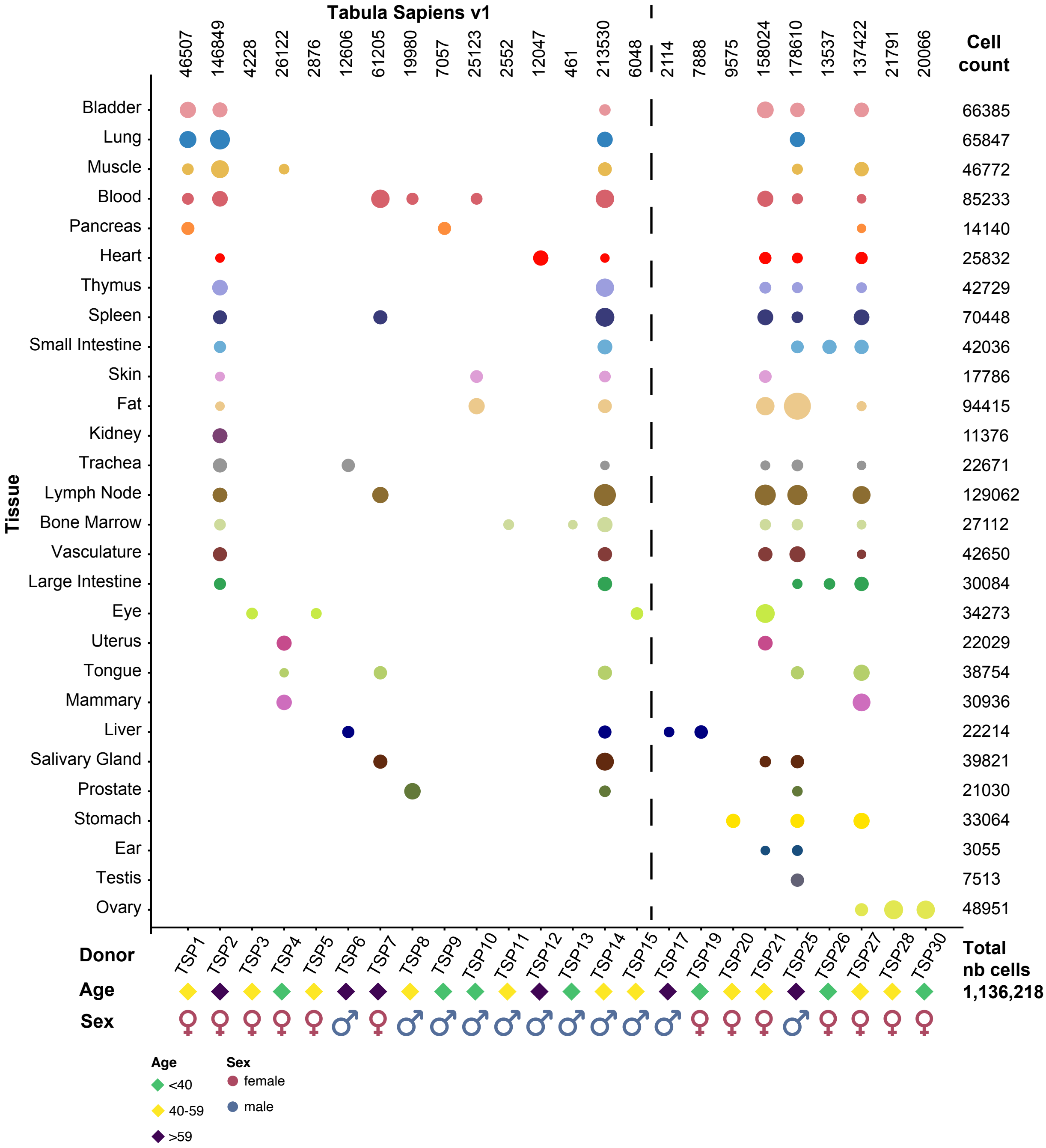

### Supp. Figure 2

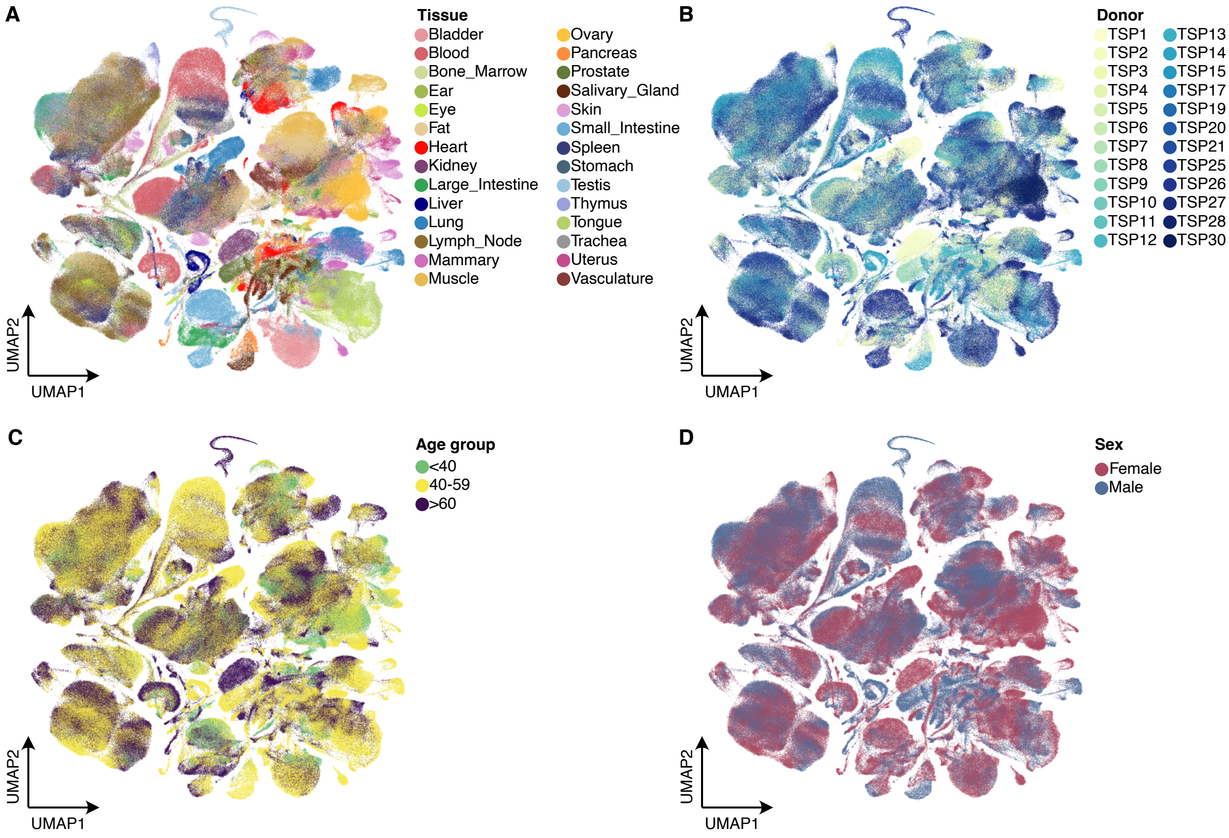

### Supp. Figure 3

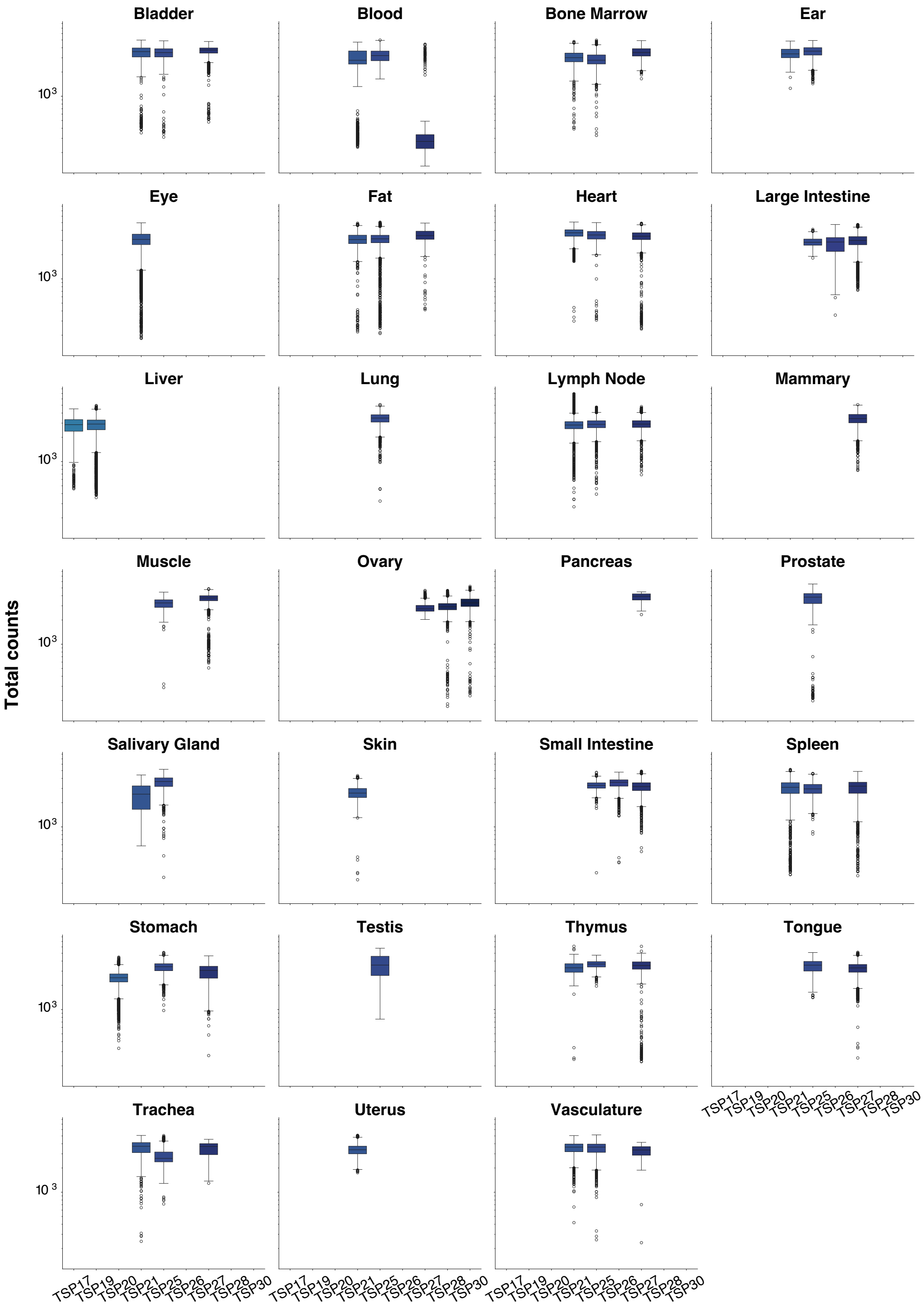

### Supp. Figure 4

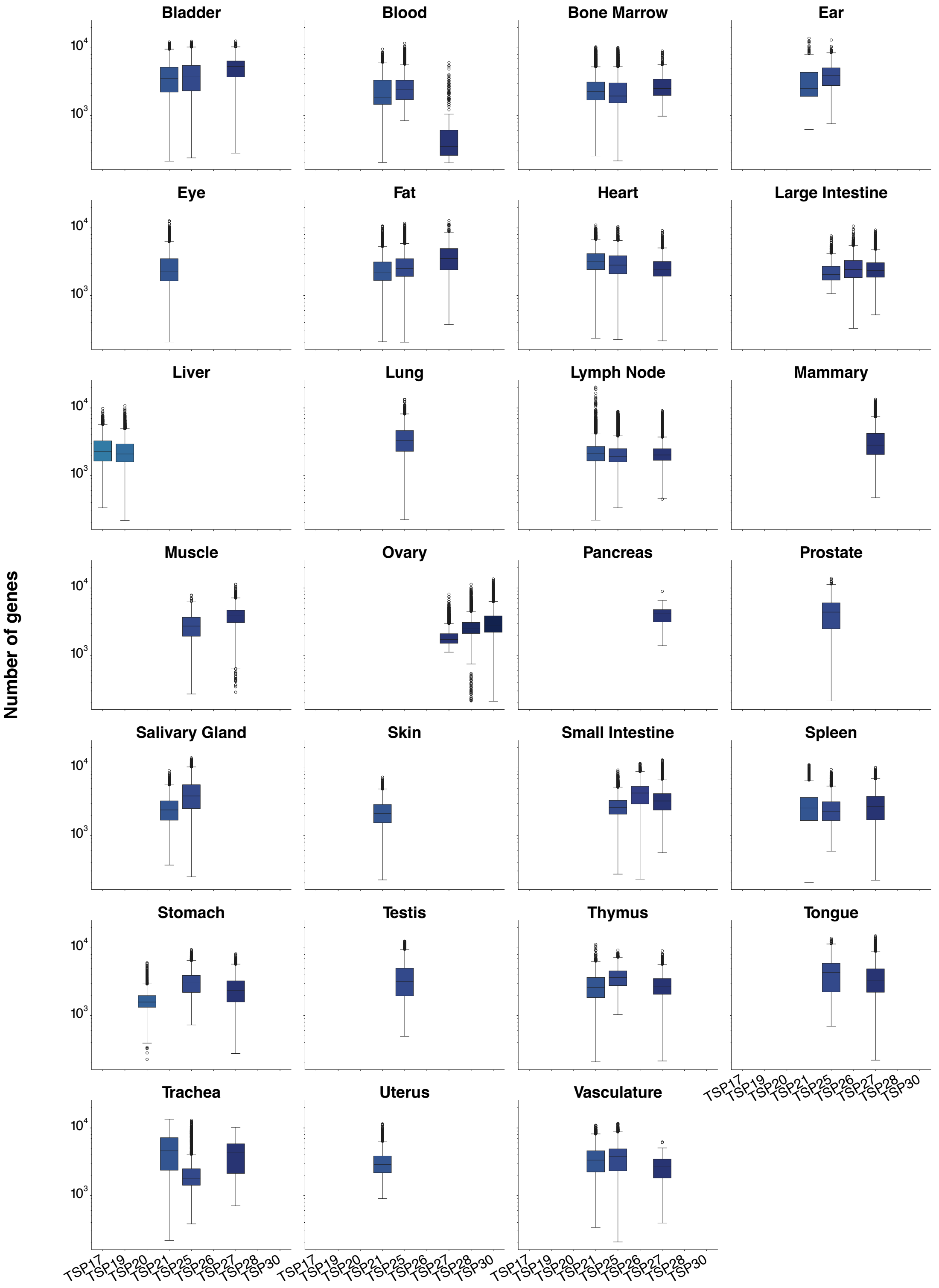

### Supp. Figure 5

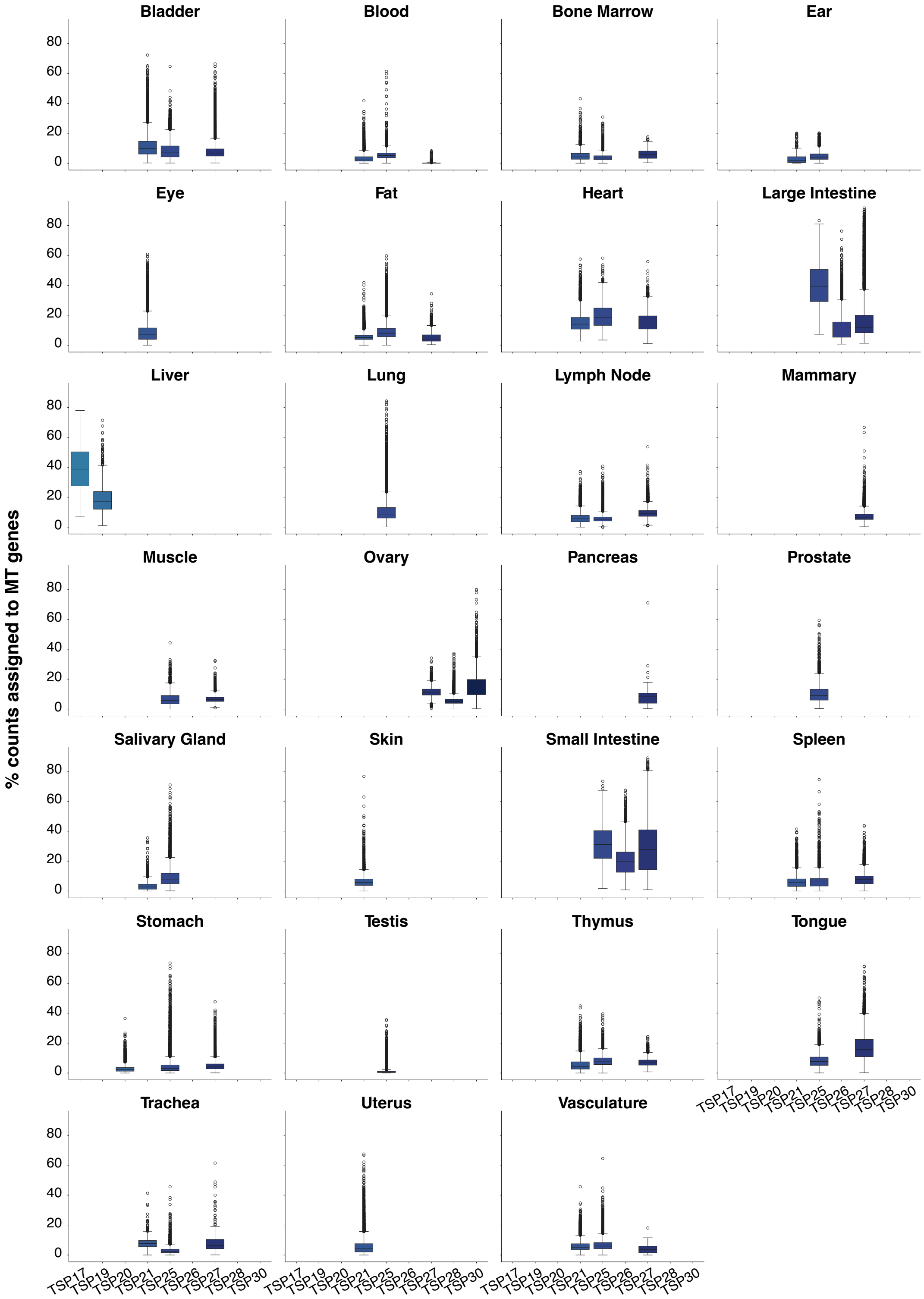

### Supp. Figure 6

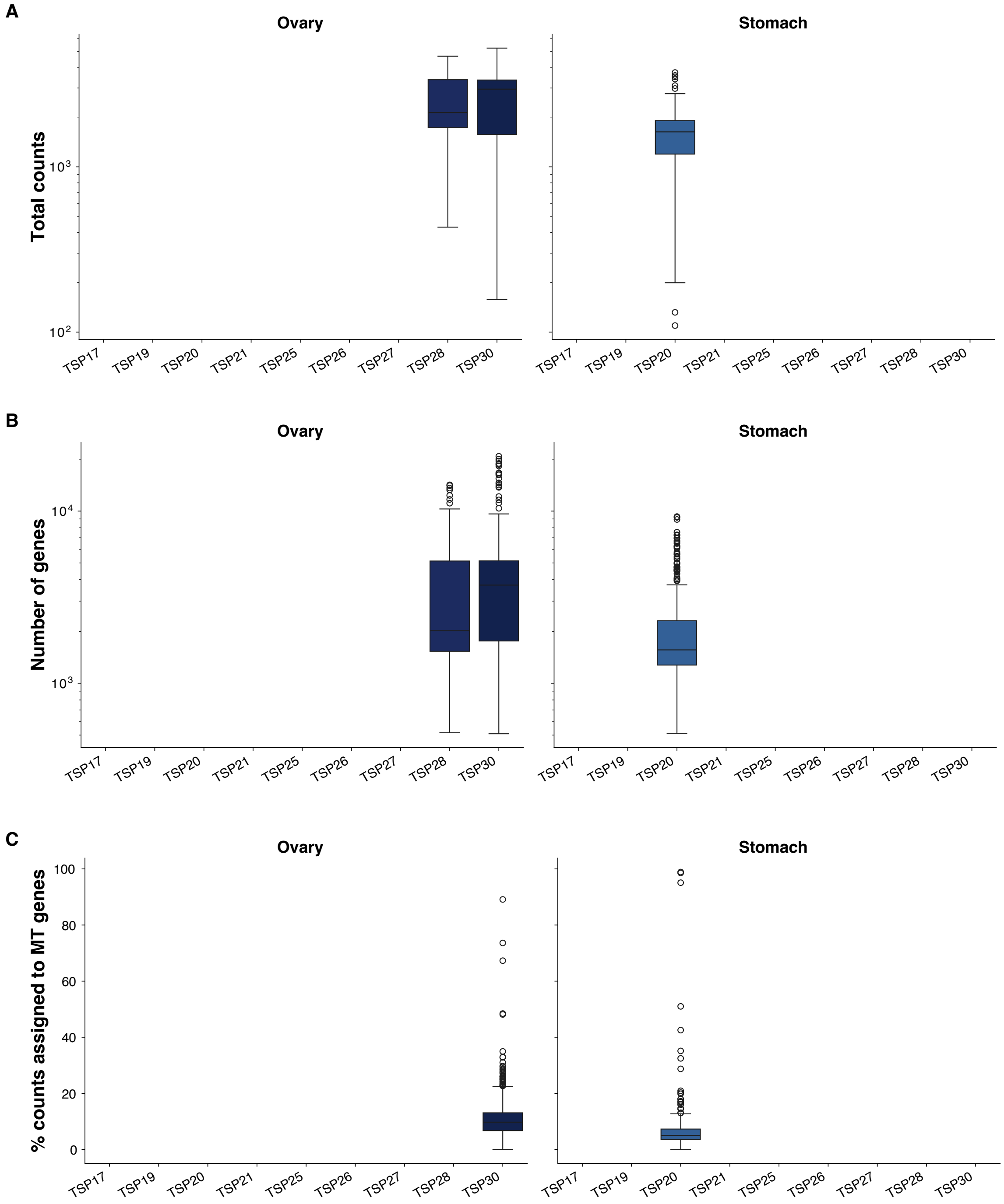

### Supp. Figure 7

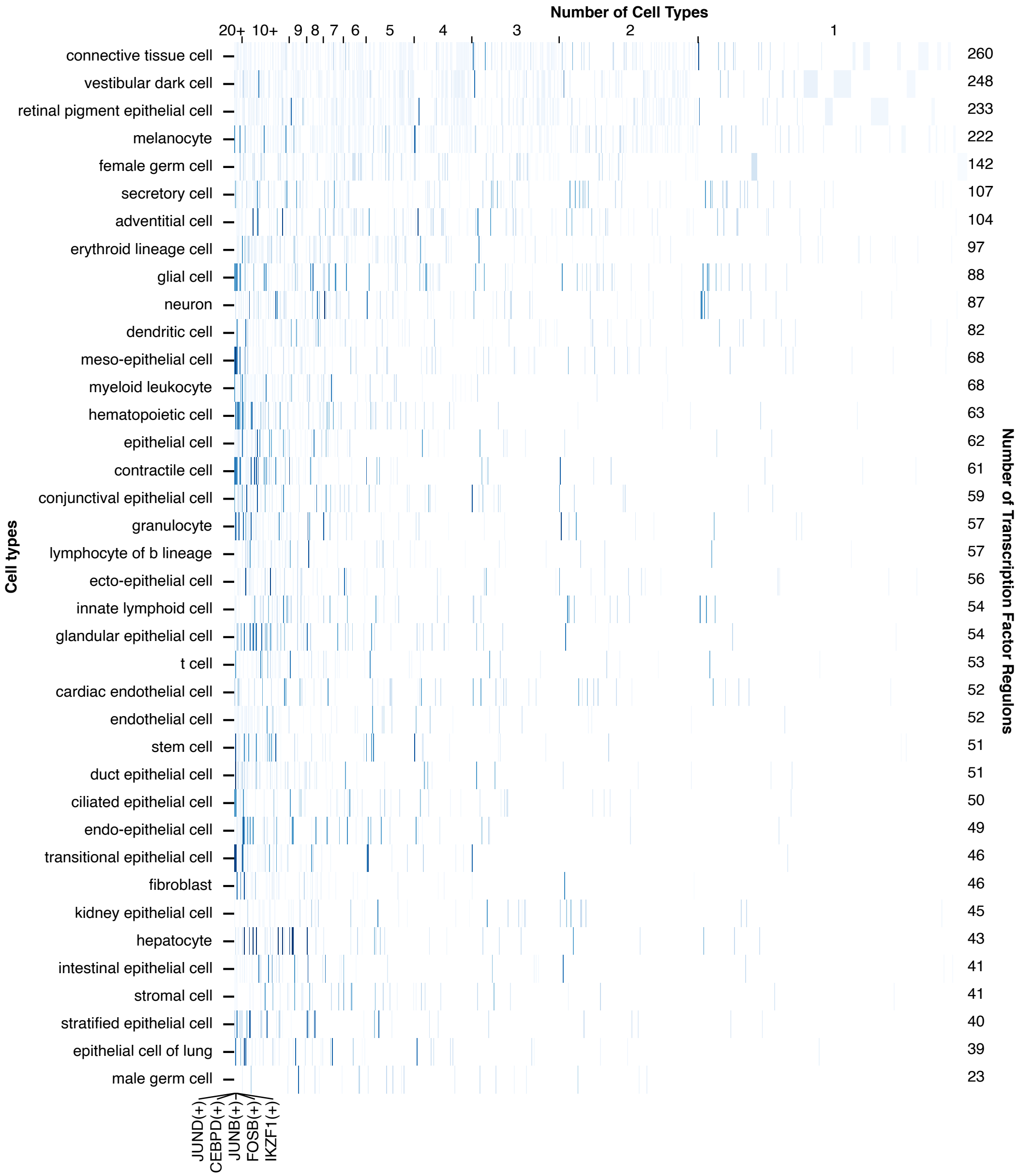

### Supp. Figure 8

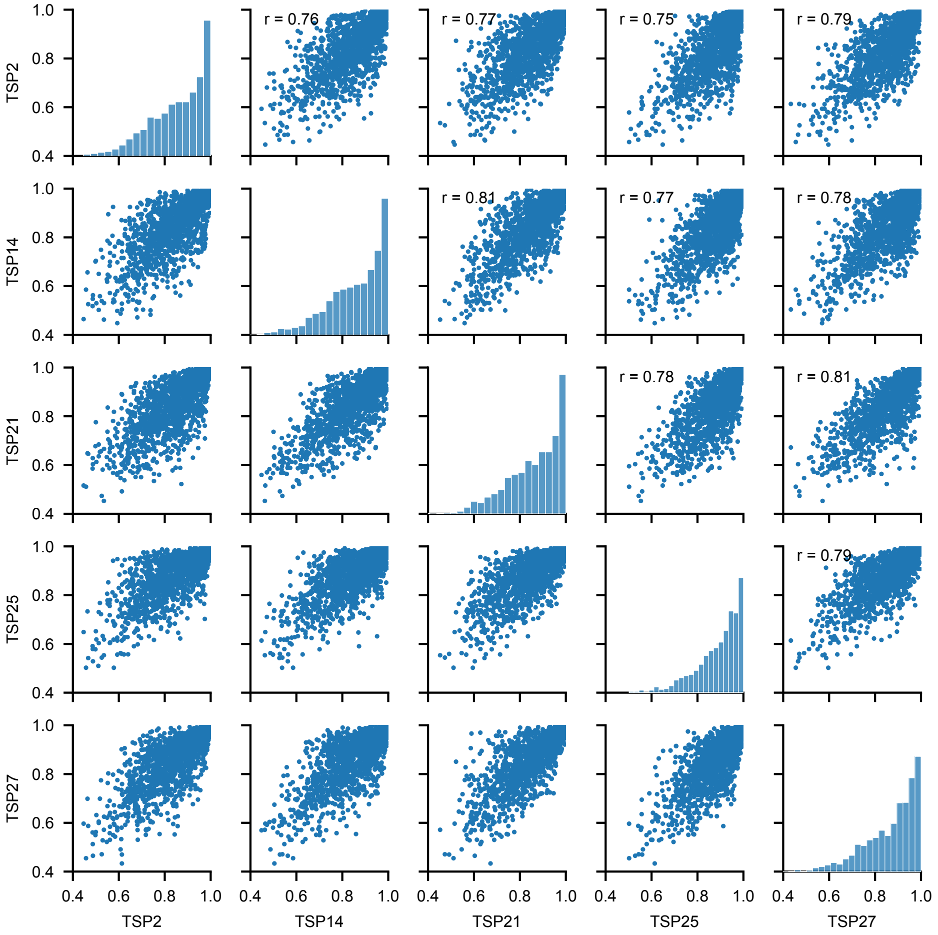

### Supp. Figure 9

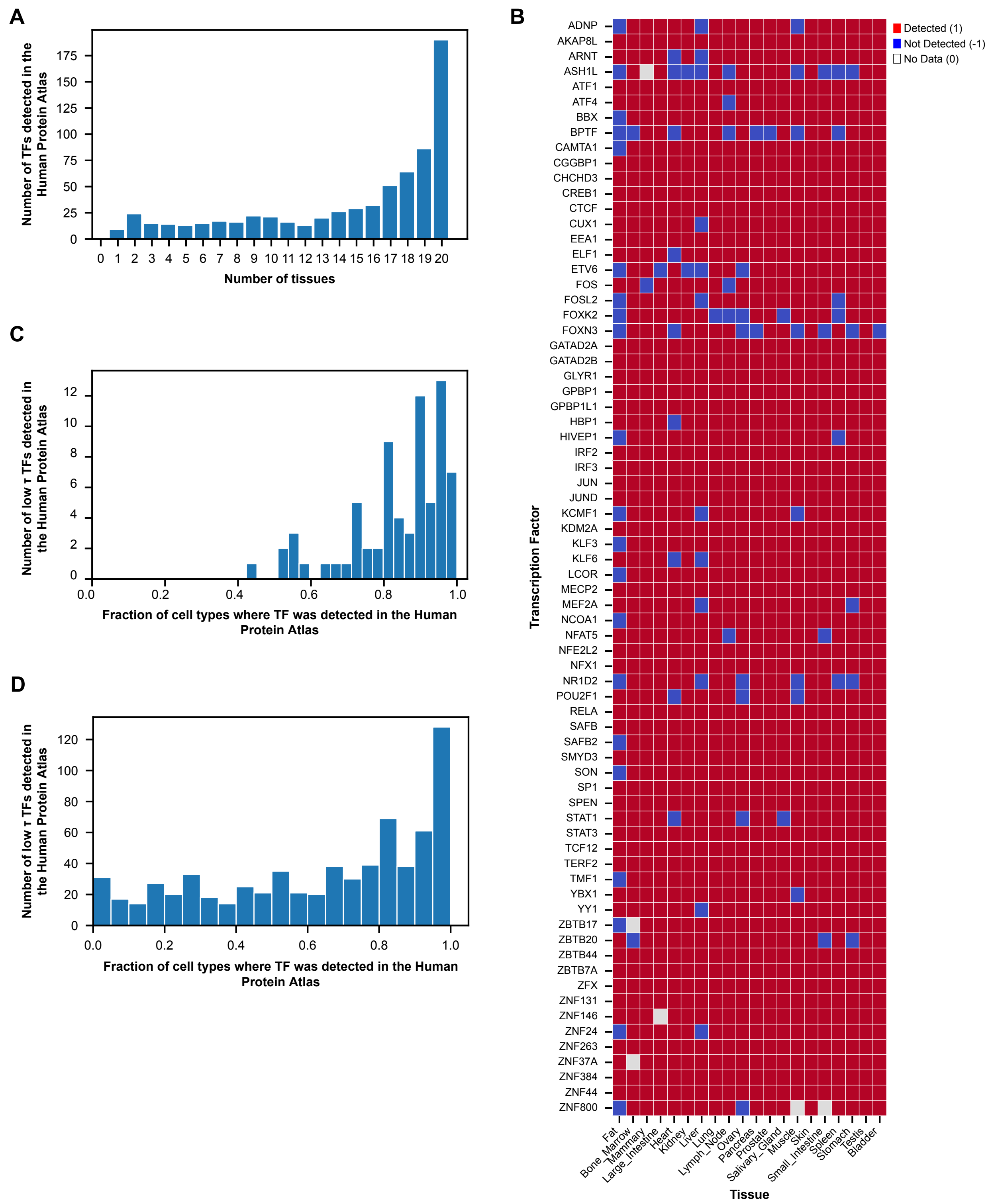

### Supp. Figure 10

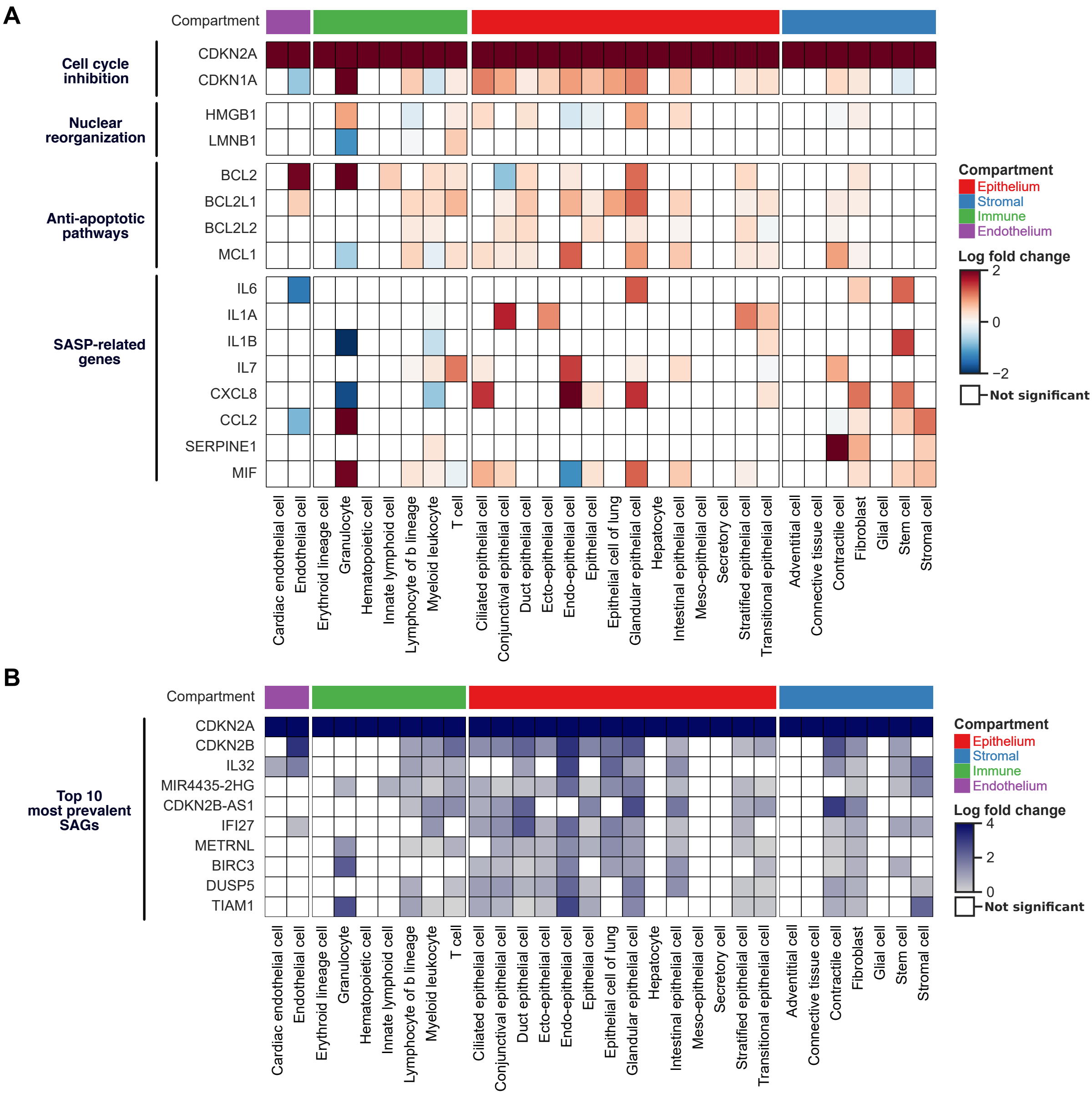

### Supp. Figure 11

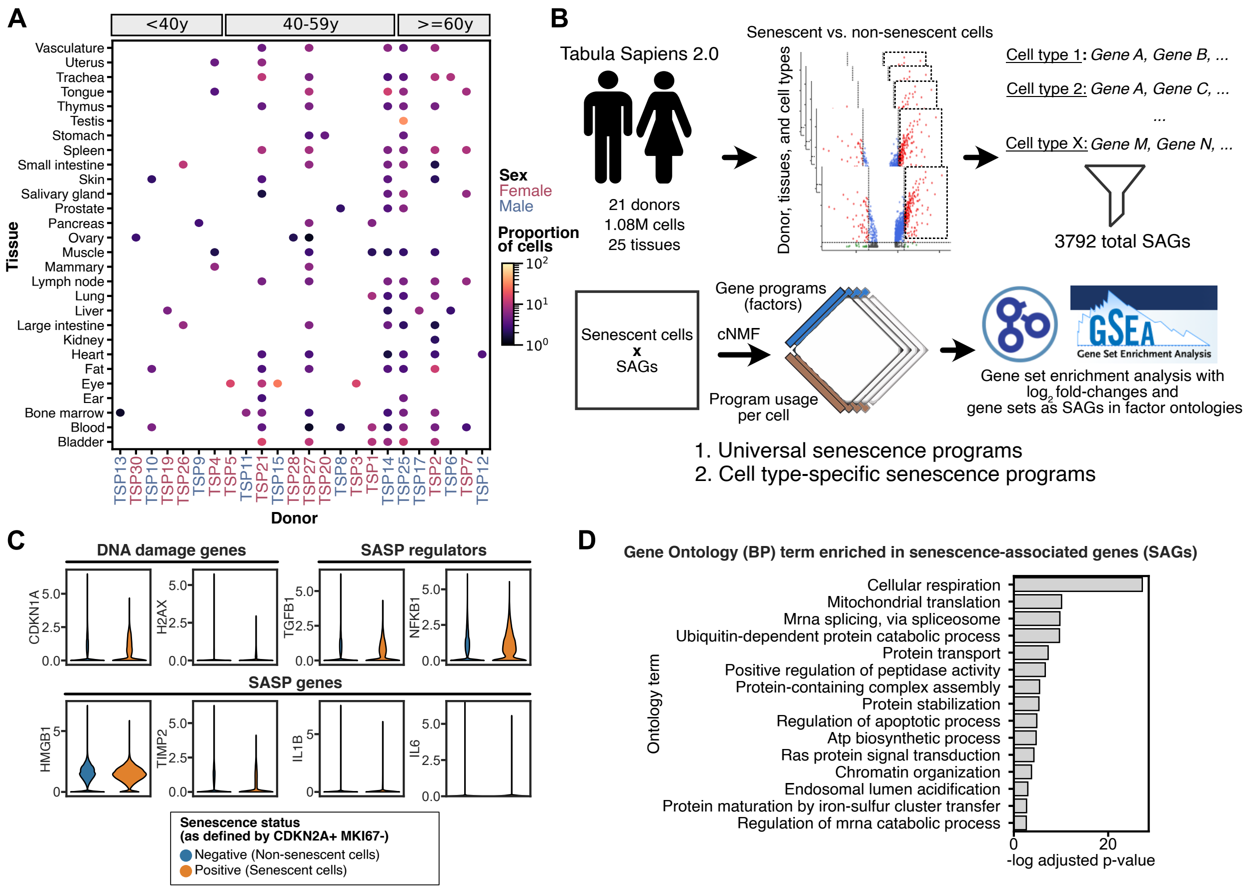

### Supp. Figure 12

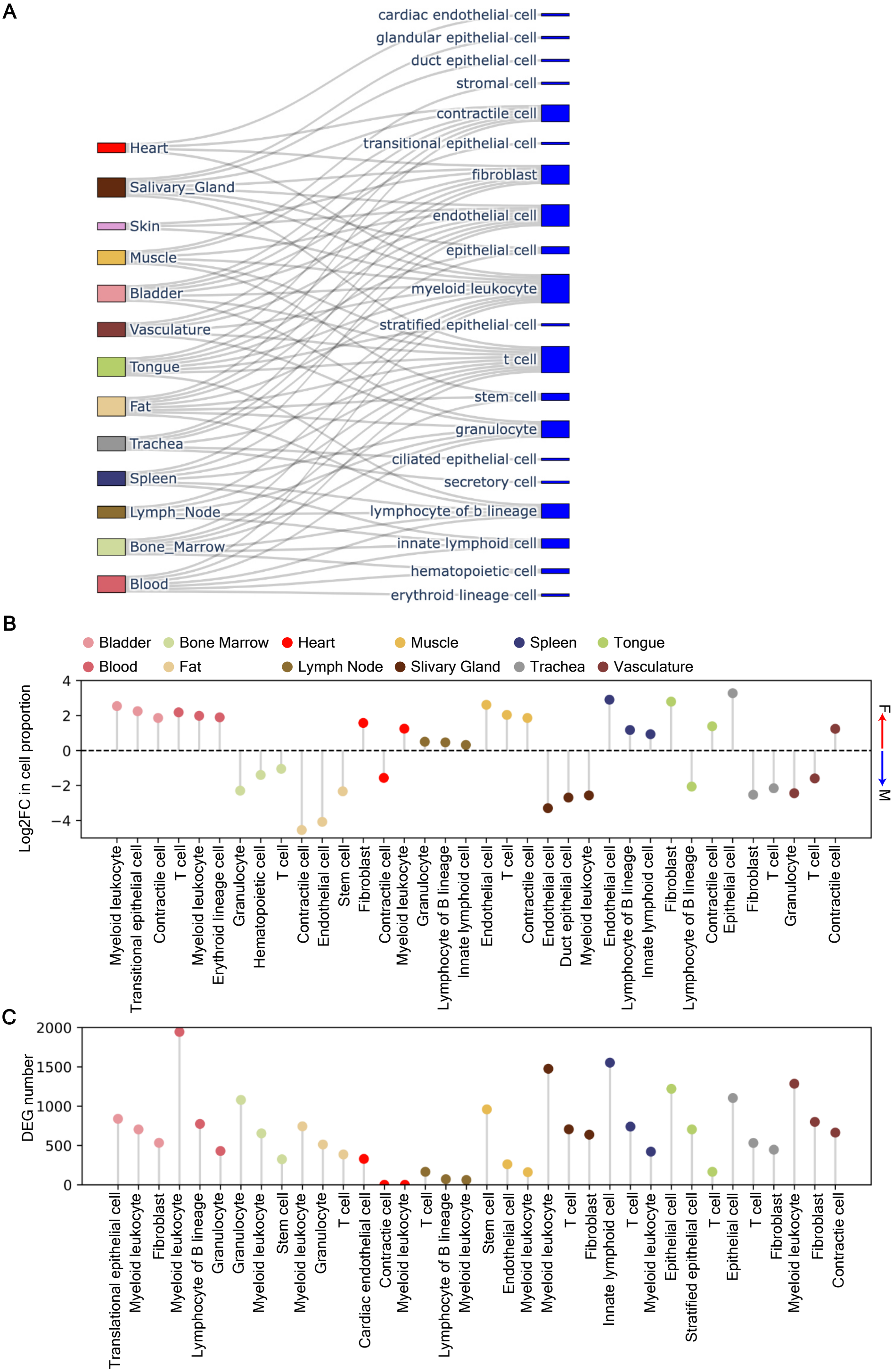

### Supp. Figure 13

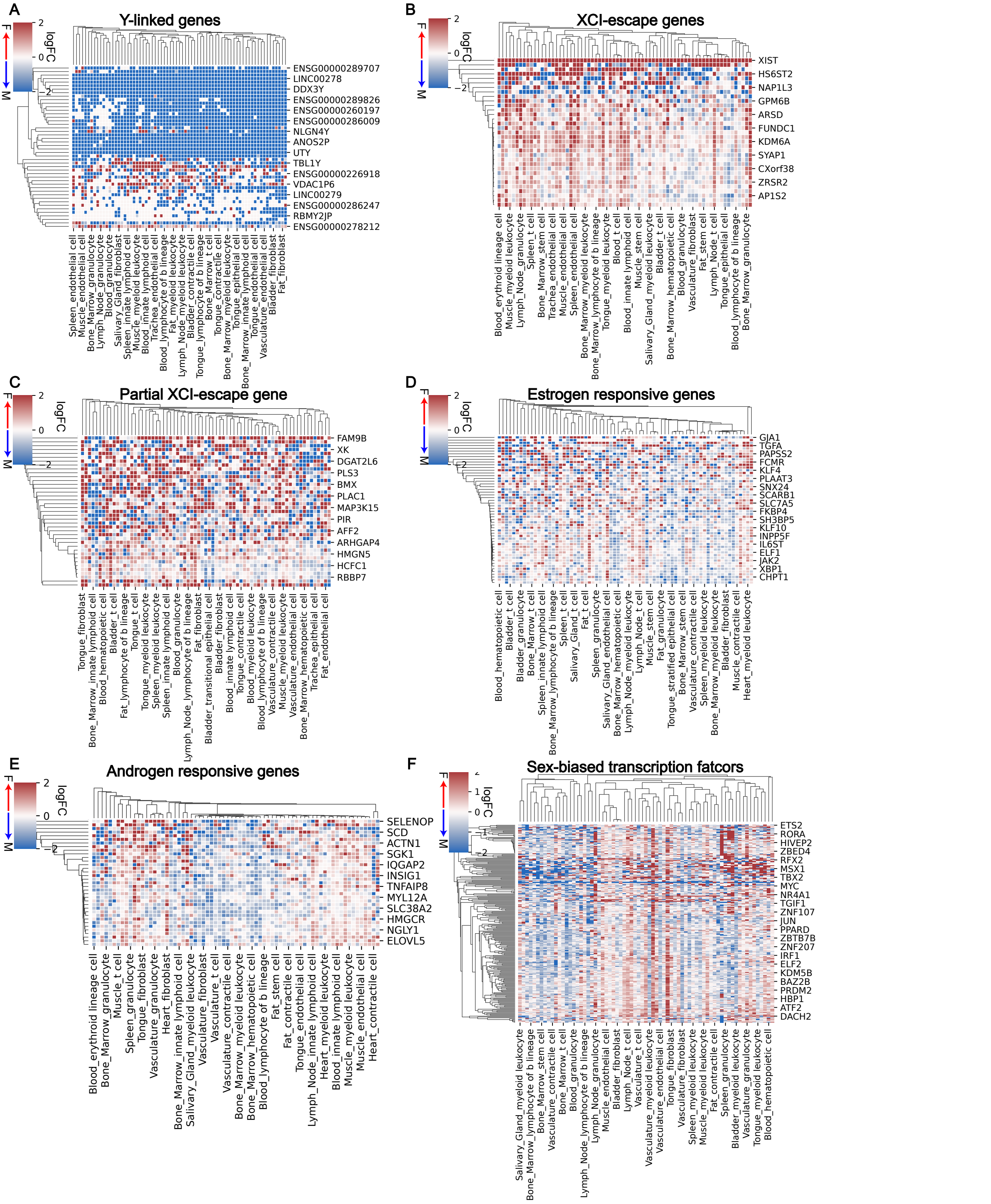

### Supp. Figure 14

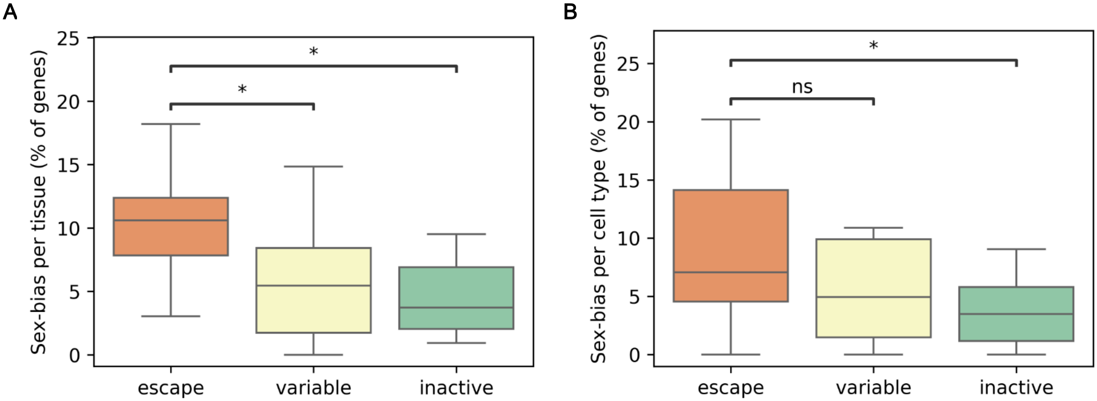

### Supp. Figure 15

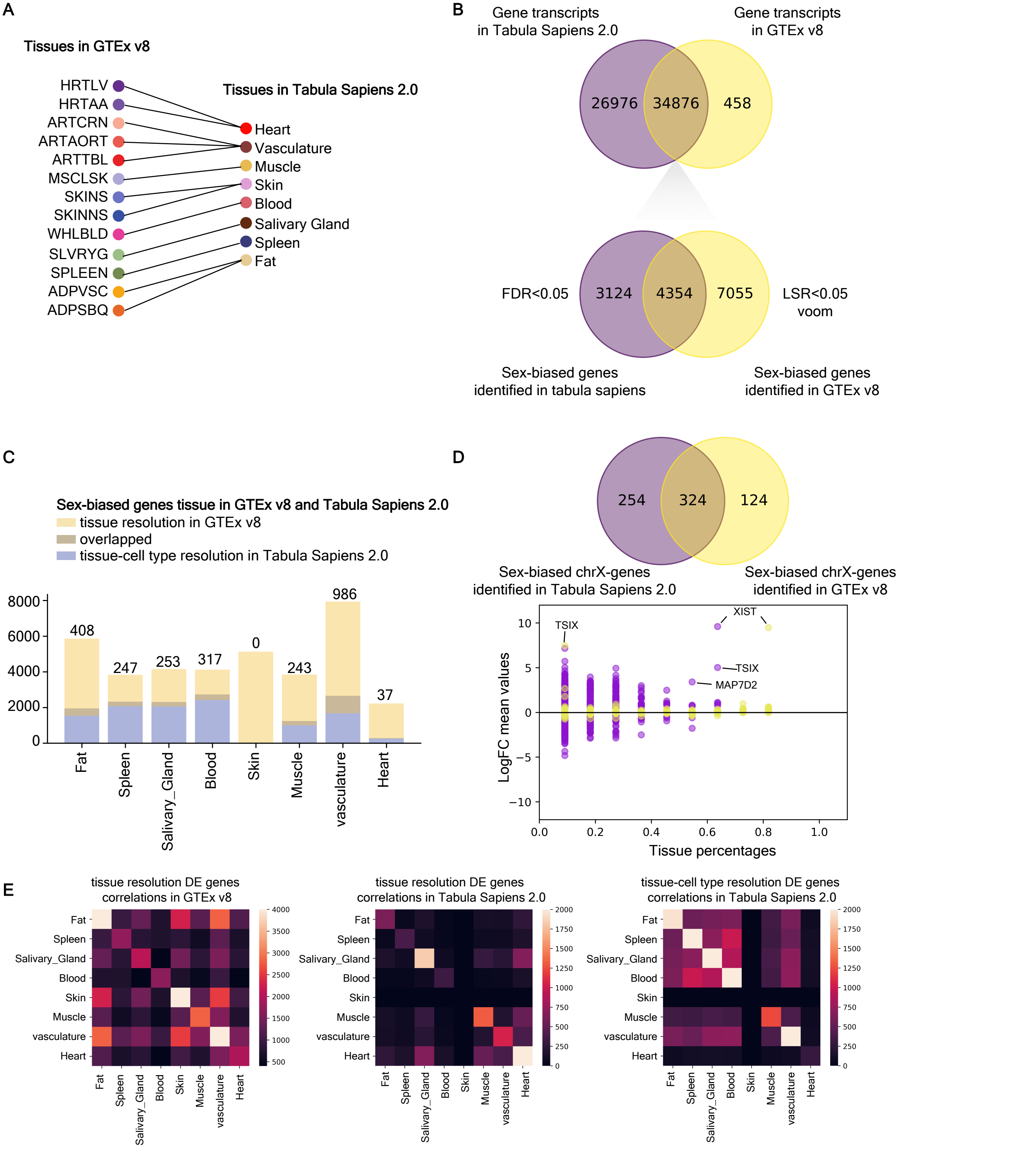

### Supp. Figure 17

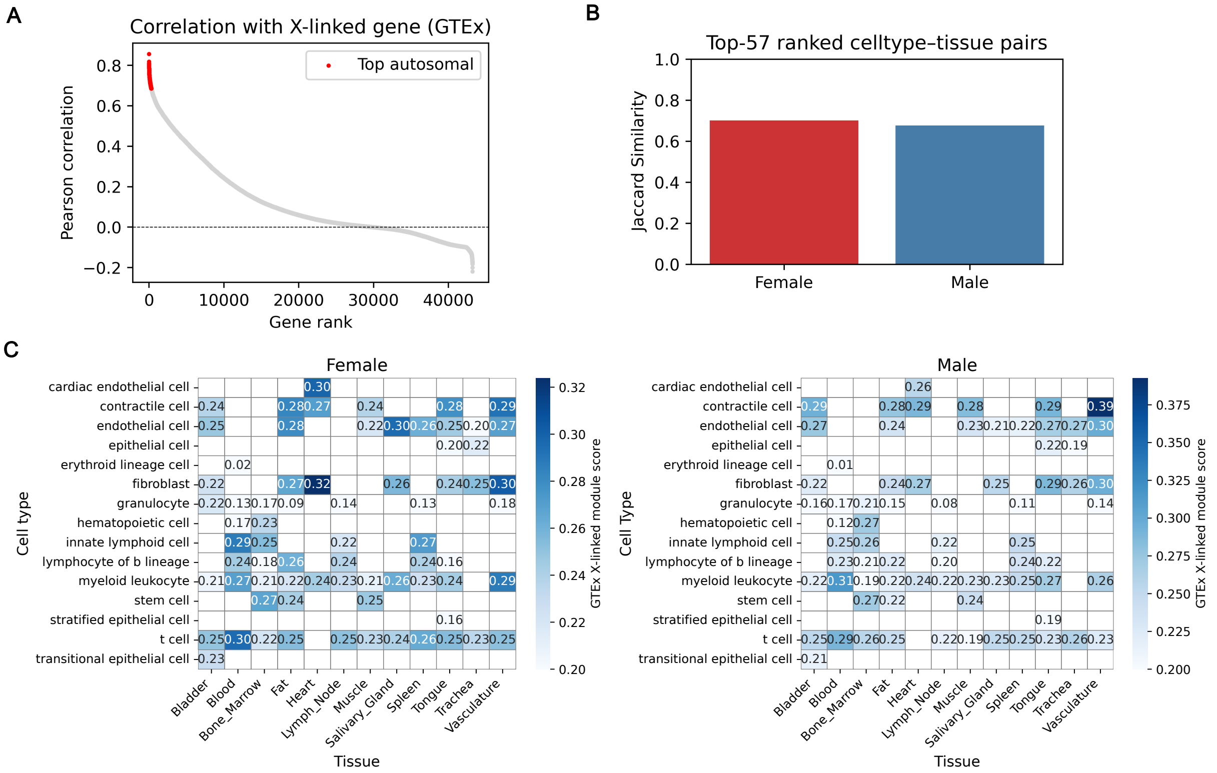
