## Supplementary material for "Tabula Sapiens reveals transcription factor expression, senescence effects, and sex-specific features in cell types from 28 human organs and tissues": Supp. Figure 16

A

| Genes | Tissues | Cell types |
|---|---|---|
| HSPA1A | Vasculature<br>Trachea<br>Fat<br>Bone Marrow<br>Bladder | epithelial cell<br>myeloid leukocyte<br>hematopoietic cell<br>t cell<br>fibroblast<br>contractile cell |
| HSPA1B | Vasculature<br>Tongue<br>Bone Marrow<br>Bladder | epithelial cell<br>stem cell<br>myeloid leukocyte<br>t cell<br>fibroblast |
| NABP1 | Vasculature<br>Trachea<br>Tongue<br>Bone Marrow<br>Bladder | granulocyte<br>stem cell<br>epithelial cell<br>t cell<br>myeloid leukocyte |

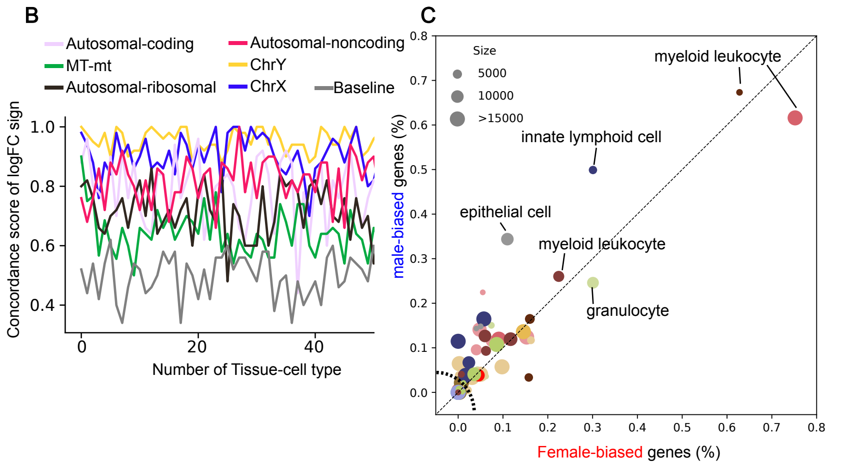


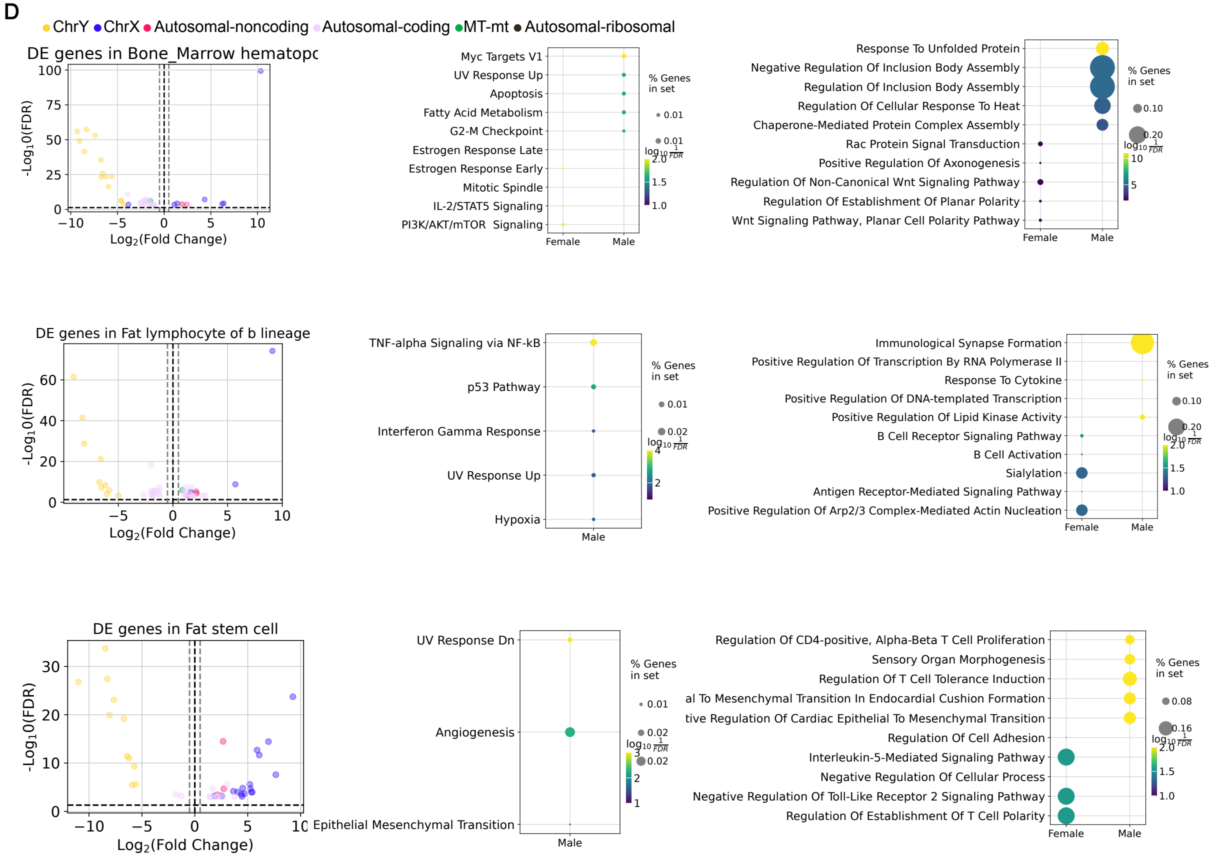
